# Structure of the Commander endosomal trafficking complex linked to X-linked intellectual disability/Ritscher-Schinzel syndrome

**DOI:** 10.1101/2023.01.25.525455

**Authors:** Michael D. Healy, Kerrie E. McNally, Rebeka Butkovic, Molly Chilton, Kohji Kato, Joanna Sacharz, Calum McConville, Edmund R.R. Moody, Shrestha Shaw, Vicente J. Planelles-Herrero, Sathish K.N. Yadav, Jennifer Ross, Ufuk Borucu, Catherine S. Palmer, Kai-En Chen, Tristan I. Croll, Ryan J. Hall, Nikeisha J. Caruana, Rajesh Ghai, Thi H.D. Nguyen, Kate J. Heesom, Shinji Saitoh, Imre Berger, Christiane Schaffitzel, Tom A. Williams, David A. Stroud, Emmanuel Derivery, Brett M. Collins, Peter J. Cullen

**Author notes:** Joint senior/corresponding authors. Joint 1^st^ authors (Michael D. Healy, Kerrie E. McNally, Rebeka Butkovic).

## Abstract

The Commander complex is required for endosomal recycling of diverse transmembrane cargos and is mutated in Ritscher-Schinzel syndrome. It comprises two subassemblies; Retriever composed of VPS35L, VPS26C and VPS29, and the CCC complex which contains ten subunits COMMD1-COMMD10 and two coiled-coil domain-containing (CCDC) proteins CCDC22 and CCDC93. Combining X-ray crystallography, electron cryomicroscopy and *in silico* predictions we have assembled a complete structural model of Commander. Retriever is distantly related to the endosomal Retromer complex but has unique features preventing the shared VPS29 subunit from interacting with Retromer-associated factors. The COMMD proteins form a distinctive hetero-decameric ring stabilised by extensive interactions with CCDC22 and CCDC93. These adopt a coiled-coil structure that connects the CCC and Retriever assemblies and recruits a sixteenth subunit, DENND10, to form the complete Commander complex. The structure allows mapping of disease-causing mutations and reveals the molecular features required for the function of this evolutionarily conserved trafficking machinery.

## INTRODUCTION

Trafficking of integral membrane proteins through the endosomal network is a central feature of eukaryotic cell biology. Proteins entering endosomal compartments are sorted between degradation within lysosomes, or retrieval and recycling to organelles that include the cell surface and the biosynthetic and autophagic compartments (1). Several protein machineries are known to be essential for sequence-specific cargo transport including the well characterised Retromer complex, and the more recently identified Commander complex (2–5). Commander regulates Retromer-independent endosomal retrieval and recycling of hundreds of proteins including integrins and lipoprotein receptors (6), and mutations in its subunits are causative for X-linked intellectual disability (XLID) and Ritscher-Schinzel syndrome (RSS), a multi-system developmental disorder characterised by craniofacial features, cerebellar hypoplasia, and stunted cardiovascular development (7–14).

Commander is composed of sixteen subunits, arranged in two subassemblies termed the CCC and Retriever complexes. Retriever is a trimer of VPS35L, VPS26C and VPS29 that shares distant homology to the well characterised Retromer endosomal trafficking machinery. Retromer is also a trimeric complex containing VPS29, either VPS26A/B (paralogues with VPS26C) and VPS35 (2, 15). The CCC complex is named for its twelve components, the two coiled coil domain-containing proteins <u>C</u>CDC22 and <u>C</u>CDC93 and the ten members of the <u>C</u>OMMD (Copper Metabolism Murr1 (Mouse U2af1-rs1 region 1) Domain) family COMMD1-COMMD10 (2, 16–19) (**Fig. 1A**). The sixteenth subunit of Commander is the DENN (differentially expressed in normal and neoplastic cells) domain protein DENND10 (also called FAM45A) (6, 16, 20–22), although its precise relationship to either Retriever or CCC is unknown.

**Fig. 1.**
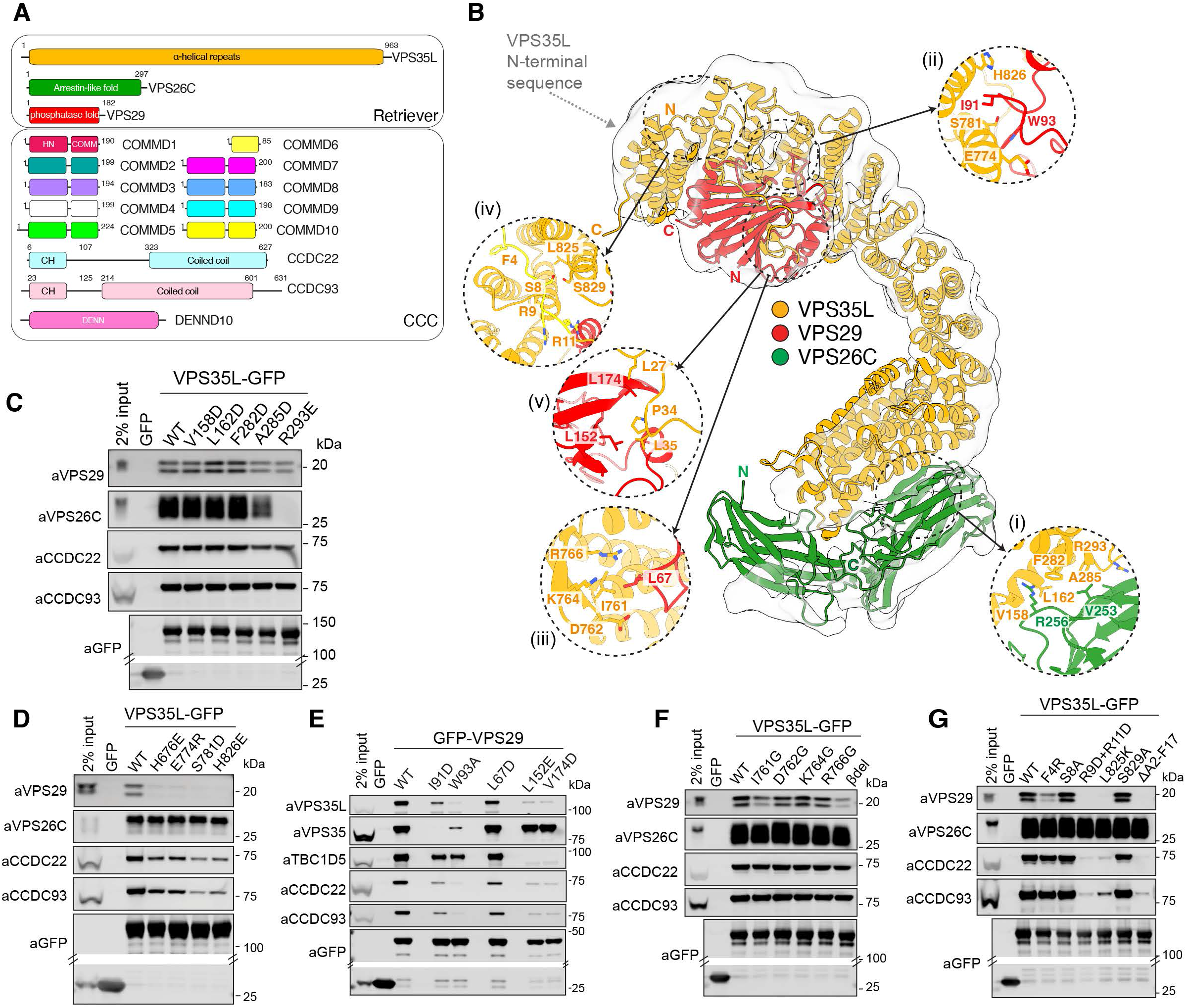
Architecture of the human Retriever complex. (A) Schematic illustration of the sixteen subunits of the Retriever and CCC sub-complexes that form the Commander assembly. **(B)** Low resolution CryoEM envelope of the human Retriever complex with docked AlphaFold2 model (**Fig. S1K**). Insets show details of specific interfaces. (*i*). VPS35L:VPS26C interface; (*ii*). VPS35L:VPS29 interaction; (*iii*). β-hairpin of VPS35L makes interactions with VPS29; (*iv*). The N-terminus of VPS35L makes an intramolecular interaction with its C-terminus; (*v*). PL motifs in the N-terminus of VPS35L interacting with the hydrophobic surface of VPS29. **(C, D)** HEK293T cells were transfected with GFP-tagged VPS35L wild type (WT) and mutants targeting the interface with **(C)** VPS26C and **(D)** VPS29 and subjected to GFP-nanotrap. **(E)** HEK293T cells were transfected to express GFP-tagged VPS29 wild type (WT) and mutants targeting the major interfaces within the Retriever assembly. **(F)** HEK293T cells were transfected to express GFP-tagged VPS35L wild type (WT) and mutants targeting the β-hairpin and subjected to GFP-nanotrap. **(G)** HEK293T cells were transfected to express GFP-tagged VPS35L wild type (WT) or mutants targeting the N-terminal sequence mediating intramolecular interactions with the VPS35L C-terminus and subjected to GFP-nanotrap. **(C-G)** All blots are representative of three independent experiments (*n* = 3). **Fig. S15** shows quantified band intensities.

Despite its importance in diverse eukaryotic species and its role in human disease, the structure of Commander is completely unknown. Even fundamental questions such as the stoichiometry of its subunits and whether it is a single stable complex remain unanswered. Its subunits are conserved in eukaryotes from humans to single-celled choanoflagellates and are primarily localized to the outer leaflet of early endosomal compartments, where the complex is required for the plasma membrane recycling of many different cargos. Most of these, including α5β1 integrin, the amyloid precursor protein (APP) and various members of the lipoprotein receptor family such as LRP1 and LDLR, contain ΦxNxx[YF] sequence motifs (where Φ is a hydrophobic amino acid) that are recruited to Commander via the adaptor protein, sorting nexin 17 (SNX17) (6, 20, 23-29). Other known transmembrane cargos such as the copper transporters ATP7A and ATP7B however, are trafficked via unknown mechanisms (30–32). As well as causing XLID and RSS developmental disorders, mutations in Commander subunits CCDC22 and VPS35L lead to hypercholesterolemia linked to reduced trafficking of LDLRs (7-10, 12-14). The Commander complex is also required for cellular infection by human papilloma virus (HPV) (6), and CRISPR knockout screens have shown that Commander is essential for SARS-CoV-2 infection, potentially by regulating endosomal cholesterol homeostasis (33–36). In addition to its role in trafficking, early studies of individual Commander subunits such as COMMD1 and COMMD7 implicated these proteins in regulating NF-κB levels and subsequent transcription pathways through interactions with distinct cullin-containing ubiquitin ligases (12, 37–43).

The ten COMMD proteins are core components of Commander and the constituent CCC subassembly. These all possess a C-terminal COMM domain approximately 70-80 residues in length, as well as an α-helical N-terminal (HN) domain (44, 45). The HN domain, while relatively divergent in sequence, has a conserved globular structure composed of six α-helices α1-α6 as seen in the structure of COMMD9 (45, 46). The C-terminal COMM domain has relatively high sequence similarity across the ten proteins and is composed of three anti-parallel β-strands, β1-β3, and a C-terminal α-helix α7 (45). Notably, the structure of the COMM domain requires it to form obligate dimers, where the β-strands and α-helix of two monomers are tightly interlocked in a ‘left-handed handshake’ topology that buries otherwise solvent exposed hydrophobic sidechains (45). Although the COMM domains were found to form promiscuous homo and heterodimers when co-expressed in bacteria, it is assumed that COMMD proteins have preferred, yet undescribed, dimeric partnerships *in vivo*. Apart from COMMD9 only the structure of the VPS29 subunit of Commander (shared with Retromer) has been determined (47–54). Overall, the structure of Commander is almost completely uncharacterised, and the stoichiometry of the different subunits are unclear.

Here we present a complete model of the sixteen subunit Commander complex, using a combination of *in vitro* reconstitution, X-ray crystallography, electron cryomicroscopy (cryoEM), machine learning-based molecular modelling and extensive validation by cellular knockouts, structure-based mutagenesis, and rescue studies. We show that Retriever, despite a superficial similarity to the distantly related Retromer, forms a heterotrimer with unique features that mediate its divergent function. The ten COMMD proteins assemble into a remarkable structure not previously seen in any other protein complex, forming a hetero-decameric ring from five specific heterodimers, with a precise and evolutionarily conserved order of subunit organisation. The CCDC22 and CCDC93 proteins stabilise the CCC complex, with natively unstructured N-terminal sequences forming extensive interactions that wind through the decameric COMMD ring. AlphaFold2 modelling and extensive validation allowed us to build an overall model of the fully assembled complex that shows how CCDC proteins link the COMMD decameric ring to the Retriever trimer, as well as recruiting the DENND10 subunit through a central coiled-coil structure. Finally, our work allowed us to structurally map all known missense mutations that cause X-linked intellectual disability and 3C/RSS. Many of these are found near interfaces between subunits, and our unbiased proteomic studies confirm that they perturb efficient complex formation. These studies provide key insights into the molecular basis for the assembly and function of the entire Commander assembly required for endosomal recycling of many essential transmembrane cargos.

## RESULTS

### Structure of the trimeric Retriever complex

Recombinant human Retriever (3xStrepII-VPS26C, VPS35L and VPS29-6xHis) was expressed in Sf21 insect cells using the biGBac system/MultiBac BEVS (55, 56) and isolated using TALON affinity purification and size-exclusion chromatography similar to our previous protocol (**Fig. S1A**) (6). The resultant peak contained VPS26C, VPS35L and VPS29 in a stable 1:1:1 heterotrimer (**Fig. S1B-D**). Dispersed particles with an elongated ‘footprint’-like morphology were observed in negative stain electron microscopy (**Fig. S1E**), and single particle 2D/3D cryo-EM classes were dominated by the front view of the ‘footprint’, with limited other orientations (**Fig. S1F-J**). Attempts to address the preferred particle orientation were not successful (including grid support and tilted datasets; data not shown). Due to the preferential orientation of the particles, gold-standard Fourier Shell Correlations (FSCs) are overestimated, and the 3D reconstruction was insufficient for *ab initio* model building (**Table S1**). However, a high confidence model of Retriever generated with AlphaFold2 (57–59) aligned very well with the low-resolution 3D cryo-EM envelope (**Fig. 1B; Fig. S1K; Movie S1**).

Analogous to Retromer (60–65), VPS35L has an extended α-solenoid structure that binds VPS26C and VPS29 at the amino-terminal and carboxy-terminal ends of the solenoid respectively (**Fig. 1B**; **Fig. S2A**). VPS35L has very little sequence similarity to Retromer subunit VPS35 (<21% identity) but both are comprised of sixteen HEAT-like α-helical repeat structures. Unlike VPS35, VPS35L also has an additional conserved N-terminal sequence (∼180 residues) that is mostly natively unstructured apart from elements that are consistently predicted to engage both the last three α-helical repeats of VPS35L as well as the VPS29 subunit (discussed below). The major interactions of the subunits within the Retriever complex were validated by mutagenesis of GFP-tagged VPS35L or VPS29 expressed in HEK293T cells and isolated using GFP-nanotrap prior to quantitative Western analysis of Retriever and CCC complex assembly. Analogous to VPS26A/B binding to VPS35 (61, 63, 64), VPS26C associates with the second and third α-helical repeats in VPS35L via its C-terminal β-sandwich subdomain (**Fig. 1C**). Structure-based mutant VPS35L(R293E) (see inset *(i)* in **Fig. 1B**) induced a >95% decrease in VPS26C binding but retained binding to VPS29 and the CCC complex (**Fig. 1C**). The major interaction of VPS29 with VPS35L is supported by the carboxy-terminal region of the VPS35L α-solenoid partially wrapping around VPS29, similar to VPS35 in Retromer (60). Mutagenesis of key binding residues, VPS35L(H826E) and VPS35L(S781D) (see inset *(ii)* in **Fig. 1B**), resulted in >95% loss of VPS29 binding (**Fig. 1D**), and reciprocal mutations VPS29(I91D) and VPS29(W93A) also decreased binding to VPS35L (**Fig. 1E**). Interestingly, reduced VPS29 interaction also correlated with a decrease in association with the CCC complex, suggesting that its binding to VPS35L is important to stabilize Retriever and CCC assembly.

### N-terminal VPS35L sequences bind a conserved VPS29 site that prevents interaction with Retromer-associated proteins

Two additional interactions unique to Retriever further promote VPS29 binding and regulate its subsequent function. Firstly, a small β-sheet extension at the base of VPS35L contacts VPS29 (see inset *(iii)* in **Fig. 1B**). Mutation of this interface with VPS35L(I761G) or complete deletion of the β-hairpin induced >50% reduction in VPS29 interaction (with CCC complex retained) (**Fig. 1F**), while reciprocal mutant VPS29(L67D) reduced binding to VPS35L by approximately 30% (**Fig. 1E**). The second unique interaction with VPS29 predicted by AlphaFold2 involves the first ∼40 residues of the extended N-terminal region of VPS35L that is not present in the Retromer VPS35 subunit (see inset *(iv)* in **Fig. 1B**). The first 17 residues form an intramolecular interaction with the carboxy-terminal region of the VPS35L α-solenoid (**Fig. 1B**), with the same structure predicted for all VPS35L orthologues across species (not shown). Deleting these residues leads to a near complete loss of VPS29 *and* CCC complex binding without affecting VPS26C association, which is replicated with both VPS35L(R9D/R11D) and VPS35L(L825K) site-directed mutants (**Fig. 1G**).

The intramolecular association of VPS35L N- and C-terminal regions serves to tether and orient two Pro-Leu (PL) motifs, ^26^PL^27^ and ^34^PL^35^ of VPS35L for binding to a contiguous hydrophobic surface on VPS29 (see inset *(v)* in **Fig. 1B**). A synthetic VPS35L peptide (residues Glu16-Ile38) representing the predicted VPS29-binding sequence interacted strongly with purified VPS29 with a *K*_d_ of 1.8 ± 0.8 μM as measured by ITC (**Fig. 2A**). We determined the X-ray crystal structure of the VPS29-peptide complex at a resolution of 1.35 Å, which unambiguously confirmed that Leu27 to Leu35 of VPS35L interact with two hydrophobic pockets on VPS29 exactly as predicted by AlphaFold2 (**Fig. 2B; Table S2**). The conserved ^34^PL^35^ side chains of VPS35L bind the VPS29 pocket defined by Val174 and Leu152 respectively, mutations L27D and L35D in VPS35L block peptide binding by ITC (**Fig. 2A**), and immunoprecipitations showed reduced binding to VPS29 and the CCC complex (**Fig. 2C**). Moreover, VPS29(L152E) and VPS29(V174D) retained Retromer association but displayed a >95% decrease in binding to VPS35L (and CCDC proteins), confirming the central importance of the ^34^PL^35^-VPS29 association for Retriever assembly (**Fig. 1E**).

**Fig. 2.**
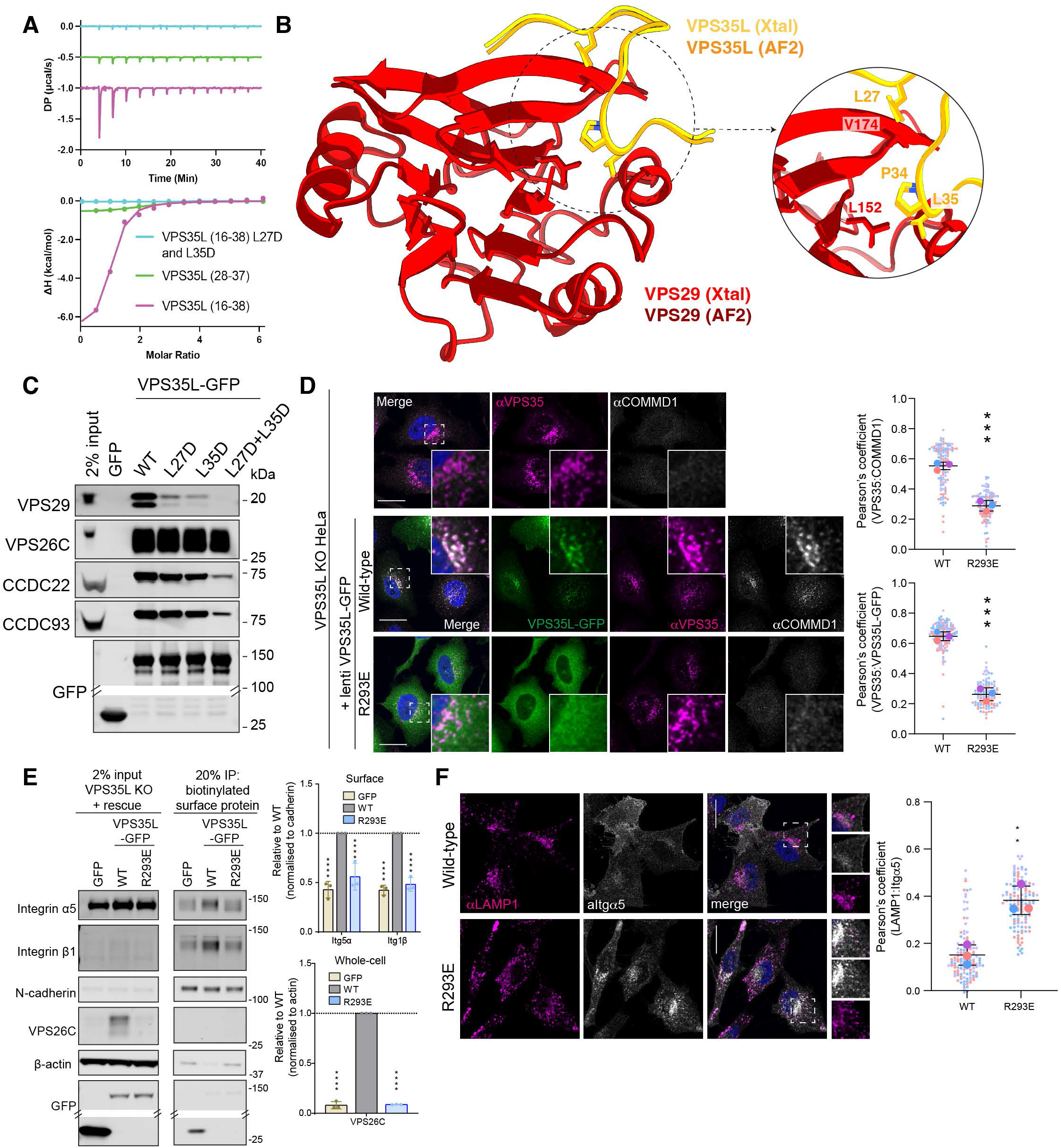
A unique structure in VPS35L regulates VPS29 interaction. (A) VPS35L peptides were titrated into VPS29 and binding affinity measured by ITC. *Top* shows the raw data and *Bottom* shows the integrated and normalized data fitted with a 1:1 binding model. VPS35L^16-38^ had a slightly higher affinity (1.87 μM ± 0.8 μM) than VPS35L^28-37^ (6.8 μM ± 1 μM) while the L27D/L35D mutant peptide showed no binding. *K*_d_ values are standard deviation (S.D.) with *n* = 3. **(B)** A 1.35 Å crystal structure of VPS29 bound to the VPS35L^16-38^ peptide confirms the binding of the core ^34^PL^35^ motif to VPS29 and extended interaction of adjacent residues predicted by AlphaFold2. The same site is also bound by PL motifs of known Retromer interactors such as TBC1D5 (**Fig. S2B**). **(C)** HEK293T cells were transfected to express GFP-tagged VPS35L wild type (WT) and mutants in the ^26^PL^27^ and ^34^PL^35^ sequences and subjected to GFP-nanotrap. 2-way ANOVA with Dunnett’s multiple comparisons test. Error bars represent the mean, S.D. *P < 0.05, **P < 0.01, ***P < 0.001, ****P < 0.0001. **(D-F)** Expression of VPS35L(R293E) in a VPS35L knock-out HeLa cell line fails to: **(D)** rescue the localisation of either VPS35L or the CCC complex to Retromer-decorated endosomes as observed with wild type VPS35L; **(E)** the expression and stability of VPS26C and the steady-state cell surface level of α5β1-integrin; **(F)** the trafficking of α5-integrin away from entry into LAMP1-positive late endosomes/lysosomes. (**C, E**) All blots are representative of three independent experiments (*n* = 3). **Fig. S15** shows quantified band intensities. (**D, F**) For each condition, Pearson’s coefficients were quantified from >30 cells per 3 independent experiments. Pearson’s coefficients for individual cells and means are presented by smaller and larger circles, respectively, coloured according to the independent experiment. Bars, mean of the 3 independent experimental means; error bars, S.D. of the 3 independent experimental means. The means were compared using a two-tailed unpaired t-test. ***= P ≤ 0.001. Error bars represent the mean, S.D. *P < 0.05, **P < 0.01, ***P < 0.001, ****P < 0.0001.

Although covering a more extensive binding surface, VPS35L N-terminal sequences closely mimic the association of VPS29 with PL motifs present in two Retromer accessory proteins, the RAB guanine-nucleotide exchange factor ANKRD27 and the RAB7 GTPase-activating protein (GAP) TBC1D5, as well as the Ile-Pro (IP) residues present in the Retromer hijacking effector RidL from the bacterium *Legionella pneumophila* (49, 50, 66) (**Fig. S2B**). This implies that Retriever will be excluded from interacting with these VPS29-binding Retromer accessory proteins. We have confirmed this using several complementary approaches. Firstly, we found that recombinant Retriever failed to bind recombinantly purified TBC1D5 (**Fig. S2C**). Secondly, GFP-TBC1D5 failed to bind endogenous Retriever in cells and GFP-VPS35L failed to isolate endogenous TBC1D5 (**Figs. S2D and S2E**). Third, unlike Retromer, we found that Retriever did not regulate the RAB7 GAP activity of TBC1D5 at the endosomal compartment, as CRISPR/Cas9 knock-out of VPS35L in HeLa cells did not phenocopy the defect in endosomal maturation and accumulation of hyperactivated RAB7-GTP observed in Retromer KO cells (67) (**Fig. S2F, S2G**). Lastly, Retriever was not bound by over-expressed mCherry-RidL (**Fig. S2H**). In all control experiments, the expected binding to Retromer of the TBC1D5 and RidL proteins was observed. These results show that while VPS29 assembled within the context of the Retromer complex provides a docking site for key Retromer accessory proteins that regulate RAB GTPases (49, 50, 66), in Retriever this accessory binding site is occluded and VPS29 is instead required to aid assembly and facilitate association with the CCC complex.

We finally assessed the importance of Retriever assembly to its function in endosomal trafficking using the model cargo α5β1-integrin. The interrelated effects of VPS29 association on coupling to the CCC complex precluded a clean mechanistic dissection of endosomal cargo recycling using VPS35L site-directed mutants targeting VPS29 binding. However, we confirmed the importance of VPS26C interaction using the VPS35L(R293E) mutant which blocks VPS26C binding but retains association with VPS29 and the CCC complex (**Fig. 1C**). In VPS35L KO HeLa cells the loss of VPS35L expression induced a reduction in VPS26C protein levels indicating their reciprocal requirement for Retriever stability. Knockout of VPS35L also resulted in (i) a loss of CCC complex association with endosomes, demonstrating a requirement of Retriever for CCC membrane recruitment, and (ii) an increased co-localisation of α5β1-integrin with the LAMP1-positive late endosome/lysosome (**Fig. 2D-F**) with a corresponding reduction in the steady-state cell surface expression of this adhesion molecule (6) (**Fig. 2E**). This indicates a defect in endosomal recycling, and these phenotypes were all rescued upon re-expression of wild type VPS35L but not by the VPS26C-binding mutant VPS35L(R293E) (**Fig. 2D-F**). VPS26C therefore plays a central role in the endosomal association and function of Commander through a mechanism that may include association with the cargo adaptor SNX17 (6), the WASH complex (30), and/or an inherent ability of VPS26C to associate with membranes like that observed for VPS26A in the SNX3-Retromer assembly (63).

### The COMMD proteins assemble into distinct heteromeric complexes

To begin to define the stoichiometry and structure of the CCC complex we co-expressed all ten human COMMD proteins in *E. coli* using polycistronic vectors. Four vectors were designed, each with the affinity purification tag on a different COMMD protein (COMMD1-His, COMMD2-His, COMMD5-His and COMMD10-His) (**Fig. S3A**). In parallel experiments, affinity purification of these four tagged-proteins followed by gel filtration and peptide mass spectrometry consistently resulted in the isolation of three homogenous and stable tetrameric subcomplexes; COMMD1-4-6-8 (isolated by COMMD1-His), COMMD2-3-4-8 (isolated by COMMD2-His) and COMMD5-7-9-10 (isolated by COMMD5 and COMMD10-His) which we referred to as subcomplex A, B and C respectively **(Fig. 3A; Fig. S3B-S3D**). These results could also be recapitulated using plasmids engineered to express only the four subunits of each stable tetrameric subcomplex (data not shown).

**Figure 3.**
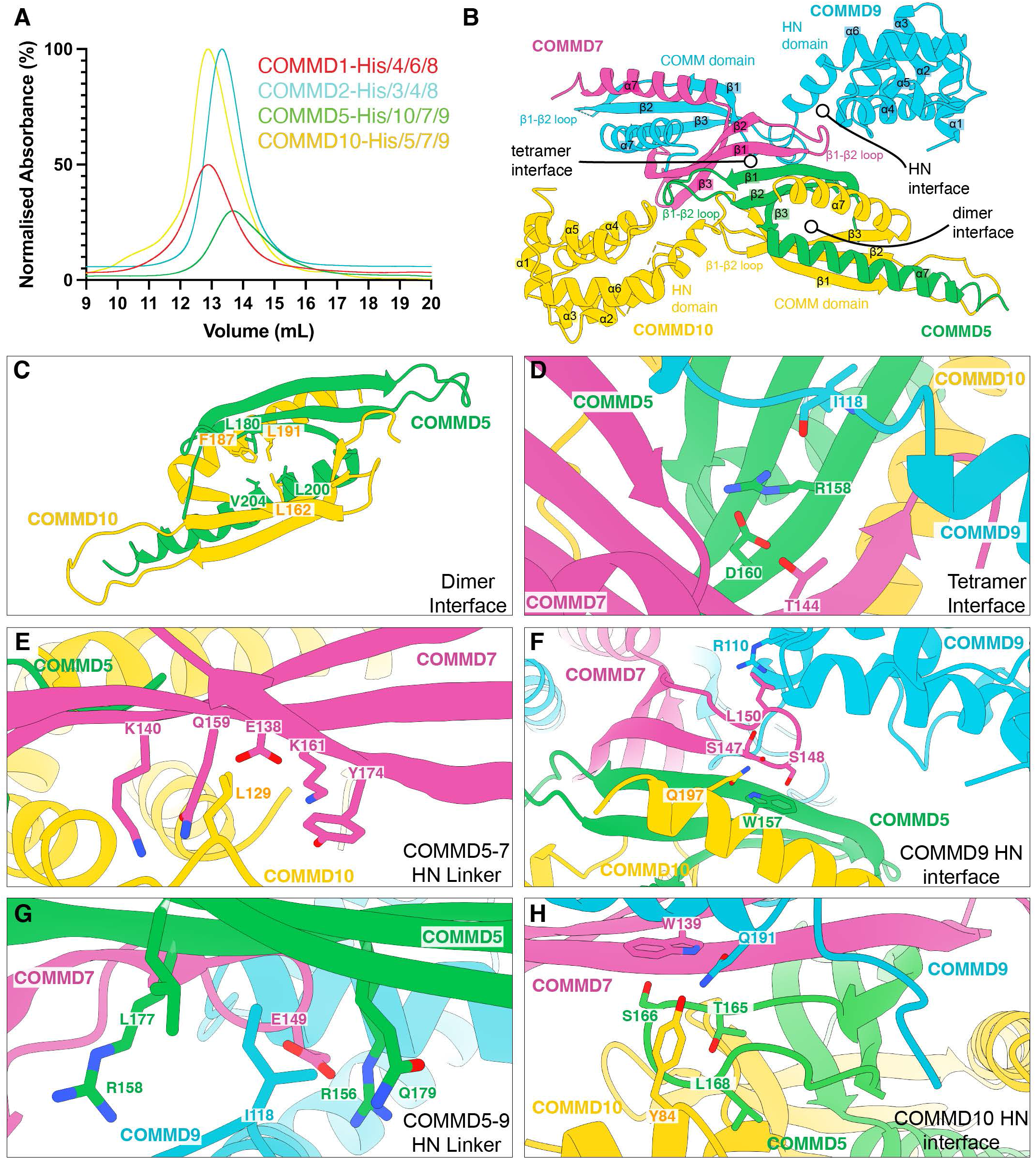
The COMMD proteins assemble into specific heteromeric complexes. (A) Purification of COMMD sub-complexes. The ten human COMMD proteins were co-expressed in *E. coli* and affinity purified via His-tags on different subunits (**Fig. S3A**) followed by gel filtration as indicated. Peptide mass spectrometry identified the subunits co-purified with each tagged protein (**Fig. S3B**) and reveals three distinct stable tetrameric complexes of COMMD1-6-4-8 (subcomplex A), COMMD2-3-4-8 (subcomplex B) and COMMD5-10-7-9 (subcomplex C). **(B)** The 3.3 Å crystal structure of the tetrameric assembly of subcomplex C (COMMD5-10-7-9). This tetramer is primarily built on three major binding interfaces that are shown in more detail below. **(C)** Highlights of key residues involved in the COMMD5-COMMD10 dimeric interface. **(D)** Key residues that form a β-sheet extension between the dimers of COMMD5-COMMD10 and COMMD7-COMMD9. **(E)** The COMMD10 key residue Leu129 binds in a hydrophobic pocket to stabilize the tetramer. **(F)** The unique COMMD9 HN domain interface in which residues from each of the four proteins in the tetramer are involved in forming a stable and specific tetrameric interaction. This image focuses on Trp157 of COMMD5. The Trp sidechain is the only strictly conserved residue across the entire COMMD family (analogous to Trp139 in COMMD7). (**G**) Details of the key interactions involving the COMMD9 linker between the HN and COMM domains centred around Ile118. (**H**) Similar to (**F**) showing details of the COMMD10 HN domain interactions with the other three subunits, centred on the COMMD7 Trp139 conserved sidechain.

While each purified COMMD sub-complex was relatively stable, only Subcomplex C produced diffraction quality crystals and its X-ray crystal structure was determined at a resolution of 3.3 Å. This revealed a 1:1:1:1 heterotetrameric assembly formed by two specific heterodimers of COMMD5-COMMD10 and COMMD7-COMMD9 (**Fig. 3B, Table S2**). The four COMM domains form an intimately assembled core structure with HN domains of COMMD9 and COMMD10 located peripherally. Electron densities for the HN domains of COMMD9 and COMMD10 were weak, and no clear densities were seen for the HN domains of COMMD5 or COMMD7 presumably due to flexibility. Notably, the structure closely matches the complex predicted by AlphaFold2 multimer (59, 68) for the four proteins (**Fig. S3E**). As expected, the four COMM domains are structurally similar to each other, composed of three anti-parallel β-sheets followed by a C-terminal α-helix (*30*).

The COMMD5-COMMD10 and COMMD7-COMMD9 dimers interact primarily via an extended β-sheet augmentation between COMMD5 and COMMD7 COMM domains to shape the tetramer (**Figs. 3B-D**). In addition, contacts between the HN and COMM domains of all four COMMD proteins contribute to the overall specificity of this assembly. Two critical contacts involve Leu129 in COMMD10 and the analogous residue Ile118 in COMMD9; these residues lie within the respective linkers between the HN and COMM domains (**Fig. 3E**). The linkers position these sidechains to reach across the tetramer interface and fit into complementary pockets on the distal COMMD7 and COMMD5 subunits respectively. In addition, the two HN domains themselves participate in a network of interactions with the other three respective subunits. For clarity we describe these two interfaces as centered around Trp157 of COMMD5 and Trp139 of COMMD7 (**Fig. 3F**), as these Trp residues are the *only* residues that are strictly conserved across all homologues and species of every COMMD protein (in COMMD9 and COMMD10 these are Trp128 and Trp139 respectively) (45). Each of these interfaces involves the N-terminal HN domain enfolding the loop between the β1-β2 strands of the COMM domain from their respective dimerisation partner. In this complex, the HN domain of COMMD10 enfolds the loop of the COMMD5 COMM domain while the HN domain of COMMD9 enfolds the corresponding loop of COMMD7. In both interfaces a serine in the β1-β2 loop forms a stacking interaction with the sidechain and a hydrogen bond with the backbone NH of the strictly conserved tryptophan residue of the neighbouring COMM domain; Ser148 of COMMD7 forms a stacking and backbone hydrogen bond with Trp157 of COMMD5, while in the second interface Ser166 of COMMD5 forms a stacking interaction and hydrogen bond with Trp139 of COMMD7. Other interactions also contribute to the specificity of each individual network (**Fig. 3G-H**), for example, Glu149 of COMMD7 makes a polar contact with Arg156 of COMMD5, and Tyr84 in the COMMD10 HN domain forms a hydrogen bond with Gln191 in COMMD9.

### Structure of the twelve-subunit core CCC complex

We next examined the role of CCDC22 and CCDC93 in COMMD interactions and assembly of the CCC complex by cloning all ten human COMMD proteins alongside CCDC22 and CCDC93 into a single biGBac construct for expression in insect cells (**Fig. S4A**). CCDC93 carried a carboxy-terminal 3xStrep-tag, and COMMD2, COMMD3, COMMD4, COMMD7, and COMMD9 contained amino-terminal Myc, SNAP, 6xHis, V5, and FLAG-tags respectively. StrepTactin-affinity isolation revealed that CCDC93 and all eleven other proteins were enriched in the desthiobiotin eluate, with size-exclusion chromatography of the peak fraction revealing a single homogeneous peak (**Figs. S4B, S4C**). Western analysis confirmed a complex of all twelve proteins, with native PAGE and mass spectrometry analysis being consistent with a 1:1 stoichiometry of each of the twelve subunits CCDC22, CCDC93 and COMMD1-10 (**Fig. S4C-S4E**). Estimates of the molecular mass of the entire Commander complex in cell lysates range from ∼500 kDa by size exclusion chromatography (17, 69) to ∼700 kDa by blue native PAGE (21) (data not shown), which is broadly consistent with the predicted molecular weight of 570 kDa if all sixteen subunits of Commander are present in a single copy.

Purified BS3-crosslinked CCC dodecameric complex was vitrified on graphene-oxide coated grids for single particle cryoEM (**Fig. S4F, S4G, Figs. S5-S7**). Data processing in CryoSPARC yielded a 3D reconstruction with an overall resolution of 3.1 Å (**Fig. S5, S6B, S7F, Table S1**). Initially a model of the dodecamer including all ten COMMD proteins and the N-terminal regions of CCDC22 and CCDC93 was constructed using AlphaFold2 multimer (57–59) (**Fig. S8A**). This predicts a structure with a highly specific arrangement of the COMMD proteins in a heterodecameric ring, with linker regions of CCDC22 and CCDC93 proteins between their N-terminal calponin-homology (CH) domains and their C-terminal coiled-coil domains tightly entwined through the COMMD assembly. The model was readily docked into the cryoEM map and with minimal adjustments was refined to produce an initial structure. The central ring of the COMM domains was much more clearly resolved than the peripheral HN domains, presumably due to their relative flexibility (**Fig. S7F; Movie S2**). To partially address this issue, we re-processed the data in RELION4.0. Several rounds of 3D classification of particles with and without alignment in combination with 3D refinement yielded a reconstruction with overall resolution of 3.5 Å (**Fig. S6, S7D, S7E, Table S1**). The HN domains of the COMMD proteins and the CH domain of CCDC93 were better resolved in this map, albeit still at lower resolution than the central core of COMM domains, facilitating further refinement of the model (**Fig. S6B, S7F**). Notably, an essentially identical *ab initio* structure was built using the machine-learning guided modelling software Modelangelo (70) although the structure was incomplete in many of the HN domains due to poorer density in these regions (data not shown). Although the complex studied by cryoEM includes the full-length CCDC proteins, no density is observed for the N-terminal CH domain of CCDC22 or the C-terminal coiled-coil domains of either protein, indicating significant flexibility in their relative orientation to the COMMD ring. Weak density is observed for the N-terminal CH domain of CCDC93 which is stabilized by its interaction with the HN domain of COMMD4.

The ten human COMMD proteins are found to assemble into a remarkable hetero-decameric closed ring structure (**Fig. 4; Fig. S8B)**. The arrangement of COMMD subunits around the ring follows a strict order of five heterodimers of (COMMD1-6)-(COMMD4-8)-(COMMD2-3)-(COMMD10-5)-(COMMD7-9) (**Fig. 4B-C**). This cryoEM structure is entirely consistent with *(i)* the COMMD10-5-COMMD7-9 crystal structure, *(ii)* with the tetrameric sub-assemblies observed in bacterial expression, *(iii)* and with models of the human complex predicted with AlphaFold2 across species as diverse as zebrafish and single-celled choanoflagellates (**Fig. S8C-S8E**). One surface of the ring is decorated by the N-terminal HN domains of COMMD1, 4, 2, 10, and 7, while the other consists of COMMD8, 3, 5 and 9 (human COMMD6 lacks the HN domain although it is present in other species) (**Fig. 4B**). As seen in the crystal structure of the COMMD5-7-9-10 heterotetramer, the interface between each COMMD heterodimer is mediated by specific contacts involving the four adjacent protomers (**Fig. S9A**). This leads to the precise organization of the heterodecameric ring.

**Figure 4.**
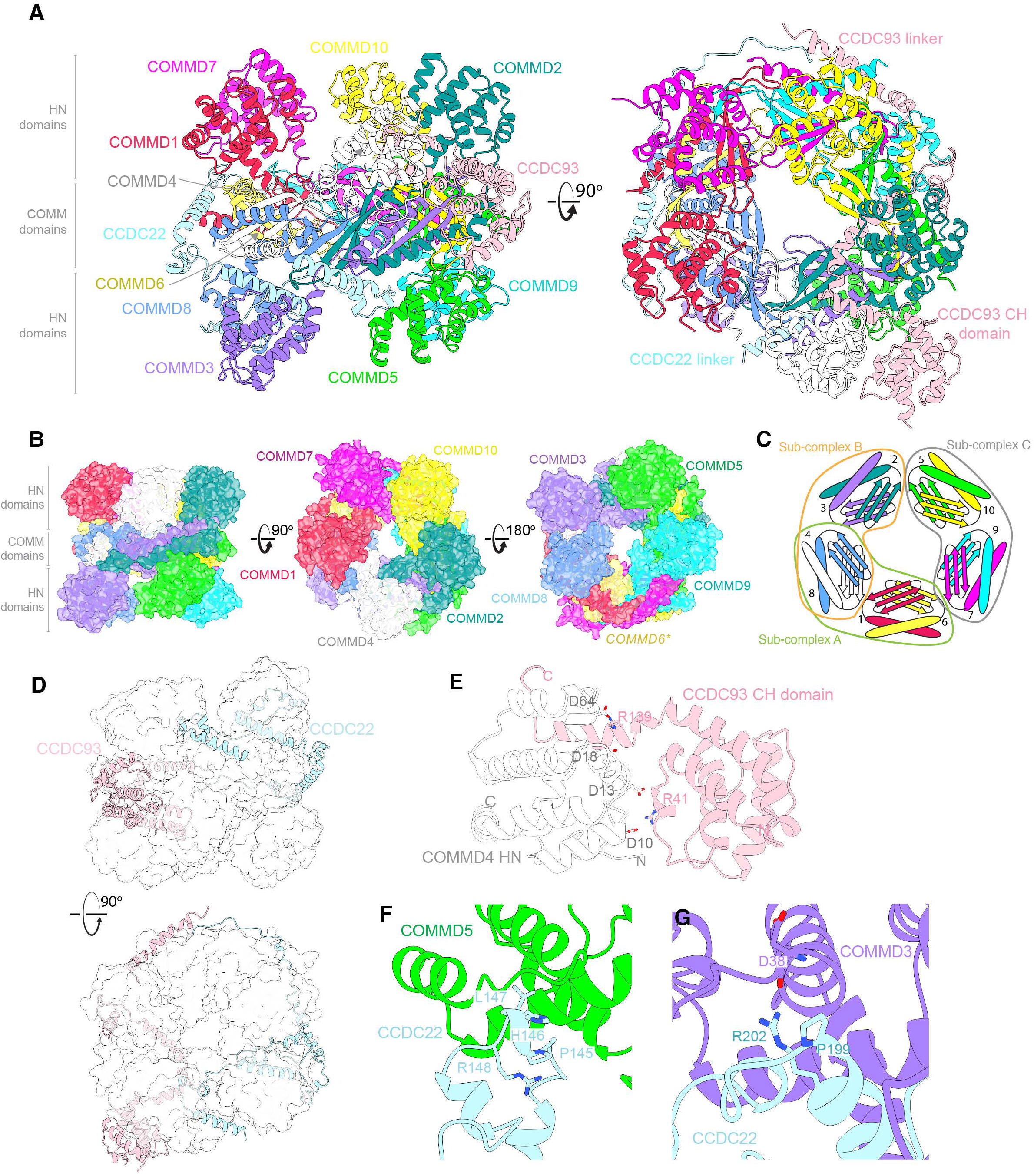
CryoEM structure of the human CCC complex. (A) CryoEM structure of the CCC complex revealing the ten COMMD proteins, the N-terminal CH domain of CCDC93 and linker regions of CCDC22 and CCDC93. Linker domains of CCDC22 and CCDC93 that are visible in our cryoEM map form intricate interactions with the decameric COMMD structure, leading to a highly intertwined structure. The CH domain of CCDC22 and extended coiled-coil regions of the CCDC proteins are not visible in current cryoEM maps due to their flexibility in relation to the stable COMMD decameric assembly. **(B)** Molecular surface highlighting the organization of the HN domains of COMMD1,4,2,10, and 7 on one side of the central COMM domain ring, and COMMD8,3,5 and 9 on the other side. Human COMMD6 lacks the HN domain although it is present in most other species. For clarity, CCDC22 and CCDC93 are omitted. **(C)** A schematic model showing the COMMD decamer and arrangement of the subcomplexes within this decamer. **(D)**. Interweaving of CCDC22 and CCDC93 within the COMMD ring. **(E)** Details of the interactions stabilizing the CCDC93 CH domain contact with the central COMMD ring, via the HN domain of COMMD4. **(F, G)** The two PxxR sequences in CCDC22 that form turn structures: **(F)** the ^145^PHLR^148^ motif binds the HN domain of COMMD5; and **(G)** the ^199^ PVGR^202^ motif binds the COMMD3 HN domain.

As well as the COMMD proteins themselves, the CCDC22 and CCDC93 linkers make extensive contacts with both the central COMM domain ring and the peripheral HN domains (**Fig. 4D; Fig. S9B-S9C**), while the CCDC93 CH domain is partly stabilised by direct interactions with the COMMD ring, in particular via the HN domain of COMMD4 (**Fig. 4E**). Two PxxR sequences in CCDC22 that form similar turn structures; ^145^PHLR^148^ (**Fig. 4F**) and ^199^PVGR^202^ (**Fig. 4G**) bind the HN domains of COMMD5 and COMMD3 respectively. The extensive interactions mediated by the linker regions of the CCDC proteins appears to enhance the stability of the COMMD ring, and this likely explains why only tetrameric sub-complexes are readily isolated when expressing the COMMD proteins in *E. coli* (**Fig. 3**). The structure is also consistent with previous truncation analyses that found N-terminal regions of CCDC22 and CCDC93 could interact with COMMD proteins but not subunits of Retriever (21, 30).

The conserved structure strongly implies that COMMD and CCDC22/CCDC93 proteins will function strictly as a dodecameric complex in the cell. The precise organization of the COMMD family members suggests the potential for different subunits to mediate distinct protein-protein interactions required for Commander activity. Several previous studies have linked individual COMMD proteins to different cellular functions. For example, COMMD1, 4, 2, 10, and 7 (the HN domains of which form one face of the decameric ring) have separately been linked to NF-κB signaling and protein ubiquitination (41, 44), while the COMMD5, 9, 3 and 8 proteins (the HN domains of which form the other face of the decameric ring, along with COMMD6) have been linked to Notch and G-protein coupled receptor signaling (71, 72). Phylogenetic analysis of the ten COMMD subunits as well as CCDC22 and CCDC93 demonstrated that all twelve proteins were present in the last eukaryotic common ancestor (LECA) (2) (**Fig. S10A, S10B**). No homologues were identified in Archaea or Bacteria. This suggests that the ten COMMD subunits likely arose by gene origination followed by gene duplication along the eukaryotic stem lineage, prior to LECA. The distribution of the ten subunits across extant eukaryotes involves parallel, lineage-specific loss in some species. These losses continued following the diversification of the major eukaryotic lineages: for example, within embryophytes (land plants), the model tracheophyte *Arabidopsis thaliana* appears to have lost Commander entirely, while the bryophyte *Physcomitrium patens* has retained four of the ancestral subunits (**Fig. S10B**).

To further assess the interdependence of all Commander subunits, the ten human COMMD proteins were each knocked out in eHAP cells, corresponding COMMD proteins were re-expressed with a C-terminal FLAG tag, immunoprecipitated and analyzed by peptide mass spectrometry (**Fig. S10C**). When used as bait, COMMD1, 3, 6, 7 and 9 all specifically isolated the entire Commander assembly, further confirming the overall inter-stability of the complex. In contrast we noted that COMMD2, 4, 5, 8 and 10 FLAG-tagged proteins were enriched only with specific subsets of COMMD proteins, and these subsets correlated with the sub-complexes observed in bacterial co-expression experiments in the absence of the CCDC proteins. Examination of the CCC structure suggests that the C-terminal FLAG-tags in these COMMD subunits may be affecting specific contacts of each of the proteins with CCDC22 and CCDC93. We speculate this leads to the loss of the CCDC interactions that stabilise the COMMD decameric ring, causing disruption of the CCC complex and loss of Retriever interaction, and further validates their importance for overall assembly. The interdependency of the COMMD and CCDC proteins for organisation of the Commander complex provides a molecular explanation for the high degree of subunit conservation across species.

### DENND10 is recruited by the coiled-coil domains of CCDC22 and CCDC93

As seen previously (6, 16, 20–22), our proteomic analyses with different COMMD subunits consistently identified DENND10 as being stably associated with the Commander assembly (**Fig. S10C**). Furthermore, DENND10 co-fractionates with all known Commander subunits by size-exclusion chromatography in both mouse and human cell lysates (69, 73), although it is not required for the stability of other Commander subunits and its deletion does not affect endosomal recycling of the canonical α5 integrin cargo (21). We used AlphaFold2 to perform a series of predictions of DENND10 to screen for and identify putative direct interactions with other Commander subunits. After exhaustive tests, a high confidence complex was predicted between DENND10 and a dimer composed of two central coiled-coil regions from CCDC22 and CCDC93 (CC1 and CC2) (**Fig. 5A; Fig. S11A**). In this predicted structure CCDC22 and CCDC93 form a V-shaped coiled-coil dimer that is bridged by conserved elements of the DENND10 DENN domain (**Fig. 5A; Fig. S11B**). To validate this model, we first tested their direct interaction *in vitro* using ITC (**Fig. 5B**). The CC1-CC2 coiled-coil regions of CCDC22 and CCDC93 were co-expressed and purified from bacteria and formed a stable dimer, which was subsequently able to bind to recombinant DENND10 with an affinity of 28 ± 6 nM. Mutations in CCDC22 and CCDC93 within the predicted binding interface either reduced the binding affinity or abolished the interaction to below detectable levels. The high affinity of the interaction was further confirmed by the formation of a stable trimer between DENND10 and the CCDC coiled-coil domains assessed by size-exclusion chromatography (**Fig. 5C; Fig. S11C**).

**Figure 5.**
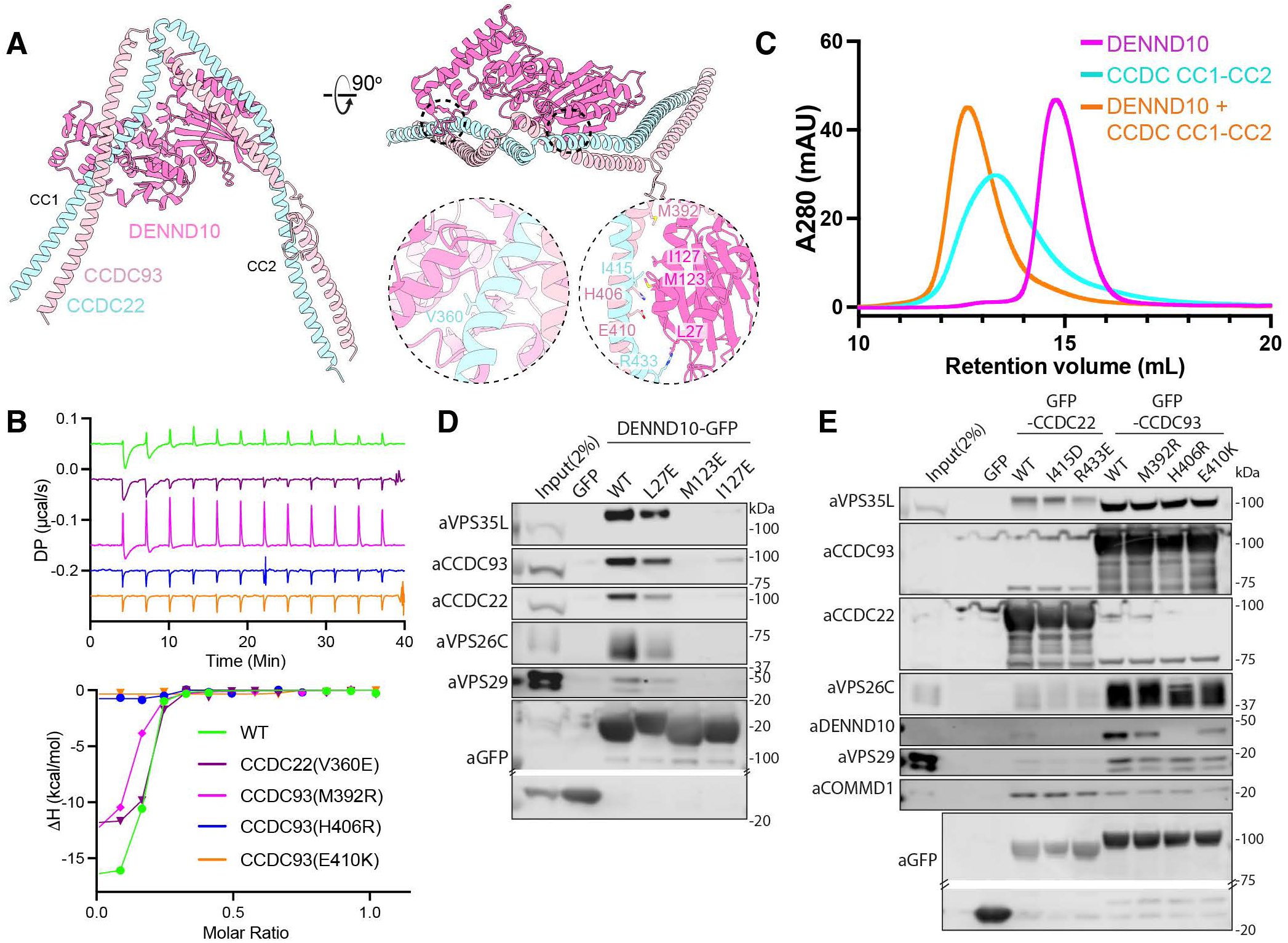
DENND10 associates with the central coiled-coil domains of CCDC22 and CCDC93. (A) Structure of the DENND10 complex with the dimeric CC1 and CC2 coiled-coil regions of CCDC22 and CCDC93 predicted by AlphaFold2. Model quality and predicted alignment errors are shown in **Fig. S11. (B)** DENND10 was titrated into purified wild-type and mutant CCDC22-CCDC93 CC1-CC2 complexes (CCDC22(325–485) + CCDC93(310–488)) and binding was measured by ITC. *Top* shows the raw data and *Bottom* shows the integrated and normalized data fitted with a 1:1 binding model. (**C**) Analytical size exclusion chromatography of DENND10 (magenta), CCDC22-CCDC93 CC1-CC2 complex (cyan) and DENND10 mixed with the CCDC22-CCDC93 forming a stable complex (orange). **(D,E)** HEK293T cells were transfected to express GFP-tagged DENND10 **(D)** or CCDC22 and CCDC93 wild type (WT) and mutants **(E)** and subjected to GFP-nanotrap. 2-way ANOVA with Dunnett’s multiple comparisons test. All blots are representative of three independent experiments (*n* = 3). **Fig. S15** shows quantified band intensities.

We further confirmed this complex in cells using additional structure-based mutations in all three proteins (**Fig. 5D and 5E**). GFP-tagged DENND10 was able to precipitate other Commander subunits including VPS35L, VPS26C, VPS29 and the CCDC22 and CCDC93 proteins, while mutations in the predicted interface DENND10(L27E), -(M123E) and –(I127E) either reduced or abolished these interactions. Reciprocal mutations in CCDC22 and CCDC93 also perturbed cellular interaction with DENND10, with the CCDC93(H406R) and CCDC93(E410K) mutations showing the strongest effect in line with the *in vitro* ITC measurements. Although DENN domains are generally thought to act as RAB GEFs, the only structure determined so far of a DENN domain-RAB complex is of DENND1B and RAB35 (74). The DENND10 sequence is highly divergent from DENND1B, and no putative RAB effector protein(s) have yet been identified, although there is evidence for an association with RAB27 (75). Furthermore, comparison with the DENND1B-RAB35 complex suggests that the CCDC proteins bind to DENND10 using overlapping surfaces. Thus, when associated with the Commander assembly DENND10 would be unable to engage a RAB GTPase in the same way as DENND1B association with RAB35 (**Fig. S11D**).

### Overall structure of the holo-Commander complex and disease mutations

Encouraged by the excellent agreement of our experimental crystal and cryoEM structures with AlphaFold2 modelled complexes we performed further predictions to assess how Retriever and the CCC complex assemble to form the larger Commander complex (see methods for details). Full-length CCDC22 and CCDC93 are predicted to form a heterodimer with four distinct coiled-coil regions (CC1-CC4) in two V-shaped structures, the first of which we showed interacts with DENND10 (**Fig. 5**). Our cryoEM structure shows that the N-terminal CH domain of CCDC93 is closely associated with the COMMD ring via the COMMD4 HN domain (**Fig. 4**), and this is also predicted by AlphaFold2. In contrast to the CCDC93 CH domain, the N-terminal CH domain of CCDC22, (which is not visible in the cryoEM map) is always predicted to extend back to form an intramolecular interaction with the V-shaped structure of the two C-terminal coiled-coil regions (CC3-CC4) (**Fig. S12A**). Interestingly, the two CCDC proteins share distant structural similarity with a number of IFT subunits of the intraflagellar transport (IFT) machinery, which also form comparable coiled-coil dimers with N-terminal CH domains (**Fig. S12B**) (76, 77).

After comprehensive testing of different potential assemblies, we identified an unambiguous interaction linking the CCC and Retriever complexes, between a conserved surface on VPS35L (opposite the VPS29-binding site) and the CCDC22-CCDC93 proteins (**Fig. S12A**). This is mediated primarily by the C-terminal CC3-CC4 coiled-coil structures with a minor interface involving the second CC2 region. By combining our experimental structures with overlapping AlphaFold2-derived models, we developed a complete structure of the Commander complex incorporating all sixteen subunits (**Fig. 6A; Fig. S13; Movie S3**). The decameric COMMD ring and trimeric Retriever are tethered together by the heterodimeric CCDC22 and CCDC93 proteins, with DENND10 associated with the apex of the structure. As mentioned, the interaction of the CCDC proteins with Retriever is mediated by the V-shaped CC3-CC4 segment at their C-terminus associating with VPS35L at a conserved surface distal from the VPS26C and VPS29 proteins. The overall shape of the complex is also restrained by the predicted intramolecular interaction between the CCDC22 N-terminal CH domain with this CCDC22/93 C-terminal structure. We validated the major interface between the CCDC proteins and VPS35L by mutagenesis of key residues, with VPS35L(R661A) or VPS35L(I710D), and CCDC22(V501D) or CCDC93(E503R) all specifically perturbing Retriever and CCC complex association without affecting assembly of either sub-complex (**Fig. 6B, 6C**).

**Figure 6.**
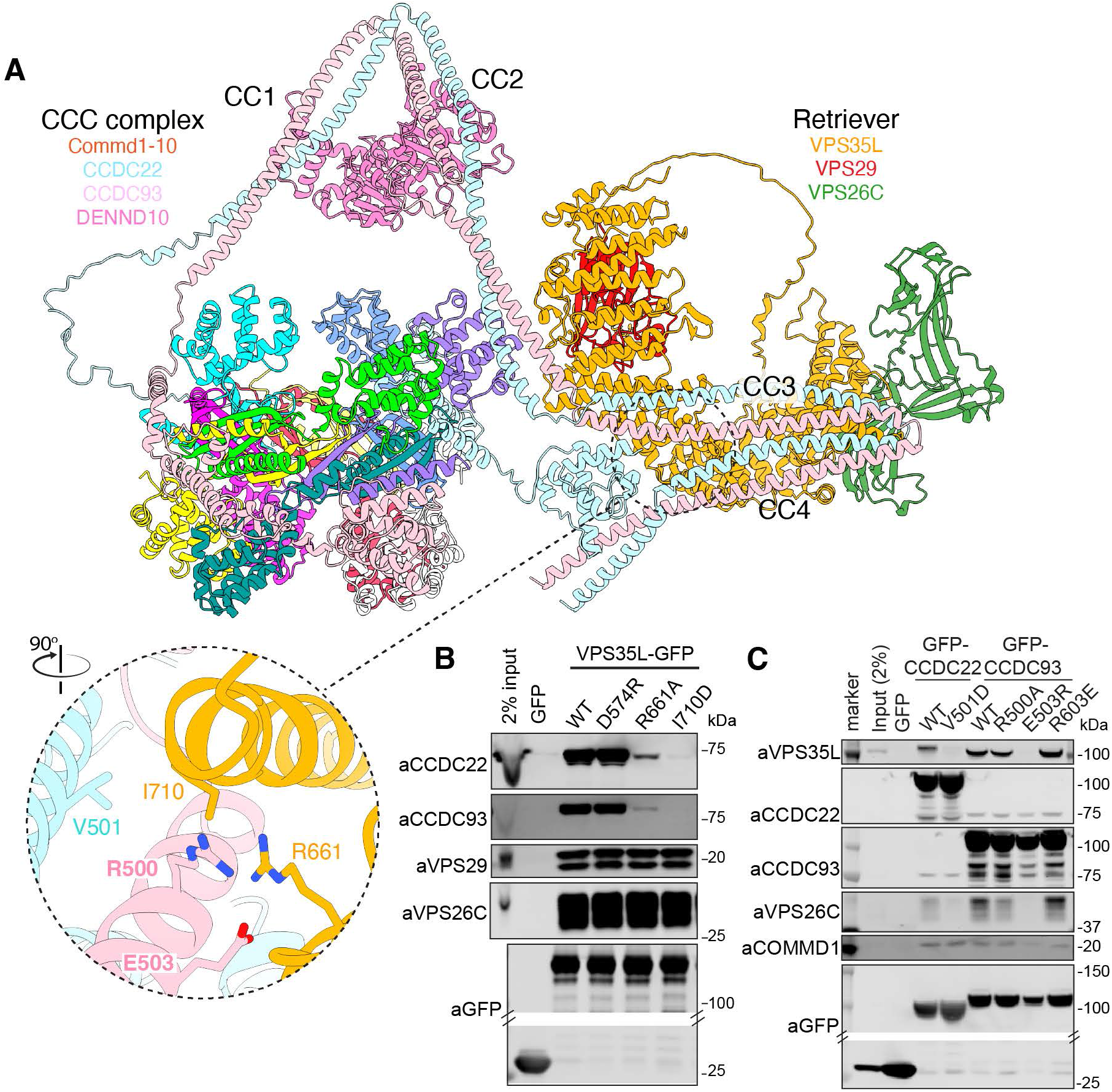
Assembly of the Commander holo-complex. (A) Overall model of the Commander complex combining cryoEM and crystal structures of the CCC and Retriever sub-assemblies and AlphaFold2 modelling of the coupling of CCC and Retriever via the C-terminal coiled-coil regions of CCDC22 and CCDC92 and the N-terminal CH domain of CCDC22. The general approach is shown in **Fig. S13**. **(B)** HEK293T cells were transfected to express GFP-tagged VPS35L wild type (WT) and mutants targeting the predicted interface with the CCC complex and subjected to GFP-nanotrap. **(C)** HEK293T cells were transfected to express GFP-tagged CCDC22 or CCDC93 wild type (WT) and mutants targeting the predicted interface with Retriever and subjected to GFP-nanotrap. All blots are representative of three independent experiments (*n* = 3). **Fig. S15** shows quantified band intensities.

In **Fig. S14** we plotted the electrostatic surface potential of the Commander complex, as well as surface conservation highlighting regions of likely functional importance. The electrostatic surface does not reveal any regions of clustered positive charge that would indicate a likely binding site for negatively charged phospholipid membranes, suggesting that membrane recruitment may be primarily dependent on protein-protein interactions, akin to Retromer (63, 78, 79). In contrast there are several regions that show a high degree of conservation. The first is a conserved patch on the surface of the CCDC22-CCDC93 coiled-coil structure, lying adjacent to the DENND10 protein (**Fig. S14B**). This aligns closely with a region previously mapped to be required for interacting with the Fam21 subunit of the actin-remodelling WASH complex (30). Another site is a deeply conserved pocket that lies directly within the interface between the VPS35L and VPS26C subunits (**Fig. S14C**). Previously the VPS26C subunit was shown to be required for coupling to the C-terminus of the SNX17 cargo adaptor protein (6), and we speculate this pocket may be involved in SNX17 recruitment. Lastly, the surface of the CCDC93 N-terminal CH domain is one of the most highly conserved regions of the Commander complex (**Fig. S14D**). Given the general actin-binding activity observed for other CH domains we speculate that it might be important for cytoskeletal interactions.

Finally, we mapped known mutations causative for XLID and RSS (7–12) onto the overall Commander model (**Fig. 7A**). This reveals a clustering of VPS35L and CCDC22 pathogenic mutations around the association interface between the Retriever and CCC complexes, providing insight into the destabilization of protein expression observed in patients harboring some of these mutations (8, 9, 11, 12). CCDC22(Y557C) is a highly conserved sidechain and lies directly within the interface with VPS35L. The N-terminal CH domain of CCDC22 is predicted to form a key interaction with the respective C-terminal coiled-coil domains of CCDC22 and CCDC93 resulting in an overall compact Commander structure (**Fig. 6A**), and a cluster of disease-causing mutations (T17A, T30A, V38M, R128Q) are predicted to destabilize this domain and its intramolecular interaction (**Fig. 7B**). VPS35L pathogenic mutations A830T, Del906 and P787L cluster towards the VPS29 interface and are anticipated to disrupt the C-terminal structure of the VPS35L solenoid. Unbiased interactome analysis comparing wildtype VPS35L with the three mutant proteins confirmed that all resulted in a pronounced loss in both VPS29 and CCC complex association (**Fig. 7C**). In contrast, the VPS35L(M931Wfs*2) and VPS35L(Del437-461) mutants (11) retained CCC complex association but had reduced binding to VPS29 (**Fig. 7C**). Taken alongside evidence that CCDC22(T17A), CCDC22(Y557C), and VPS35L(A830T) perturb endosomal retrieval and recycling of LRP1 and LDLR, and lead to the development of patient observed hypercholesterolemia (11, 30), these structural data begin to provide a molecular explanation for the perturbed stability and assembly of the Commander complex associated with XLID and RSS.

**Figure 7.**
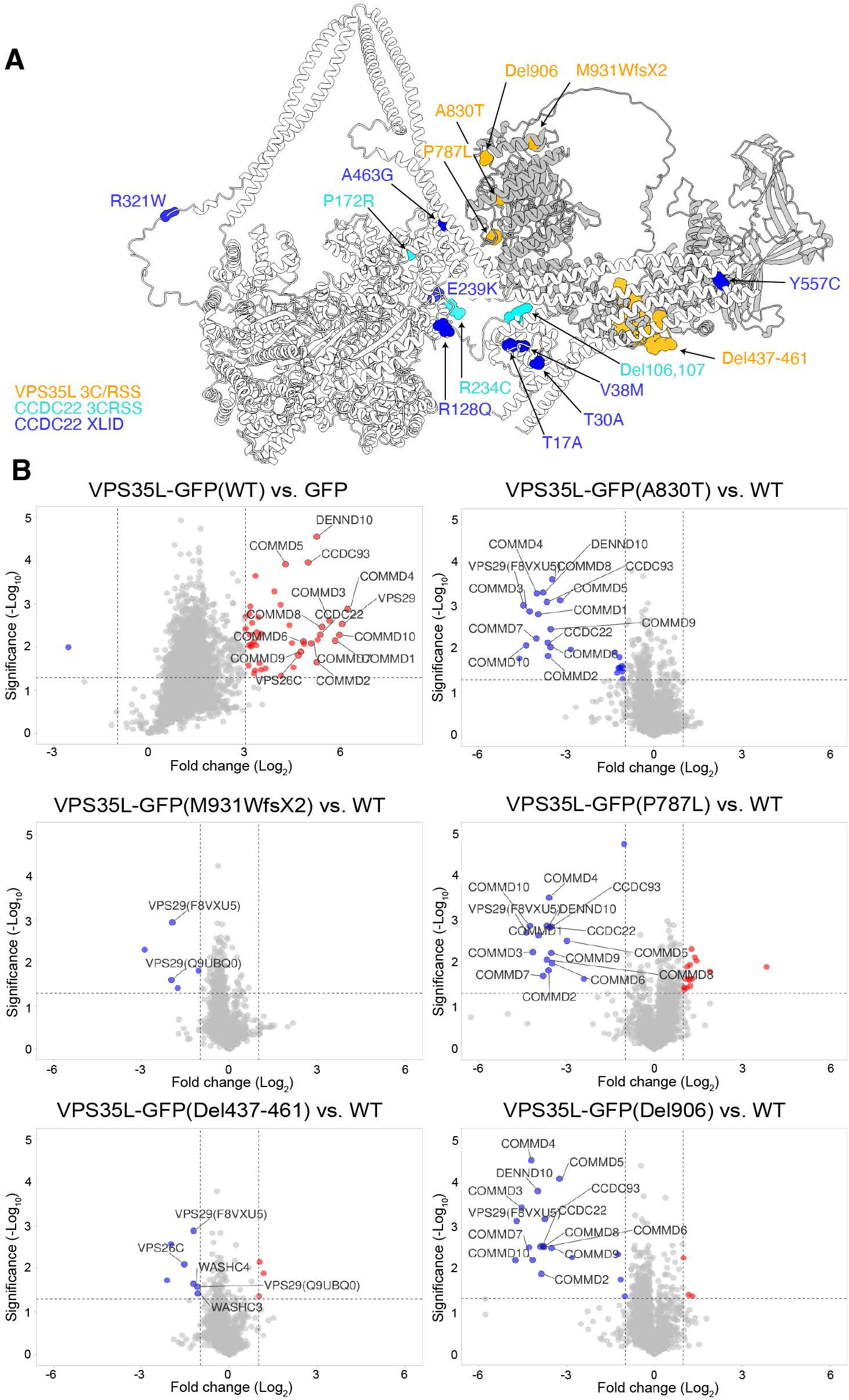
Structural and functional impacts of Commander mutations causing XLID and RSS. (A) *VPS35L* and *CCDC22* mutations associated with XLID and RSS mapped onto the Commander structure. **(B)** Volcano plots of identified interactors enriched (red circles) or depleted (blue circles) in GFP-nanotrap when comparing wild type GFP-tagged VPS35L and VPS35L mutants causative for RSS. In a single experiment, HEK293T cells were transfected to express GFP, GFP-tagged wild type (WT) and the five GFP-tagged VPS35L mutants and each were subjected to GFP-nanotrap prior to seven-plex TMT-based proteomic identification and quantification. Data is from 3 independent experimental repeats.

## DISCUSSION

Despite its fundamental role in cellular trafficking and importance in disease the molecular structure of Commander has been almost completely unexplored. Our studies provide a comprehensive understanding of how the Retriever and CCC complexes are assembled, and how they combine to form the overall Commander super-complex. The high degree of conservation of this large multi-subunit complex confirms its essential role in the regulation of membrane trafficking throughout evolution, and the structure now provides an atomic level description of its organization that explains a number of previous results including the co-dependence of each of the COMMD, CCDC, and VPS35L proteins on overall complex stability (13, 28). The conserved Commander structure and this co-dependency indicates that each of the subunits are likely to act in unison to mediate overall Commander activity, and implies that previous studies examining individual components of the complex, including our own (45), may need to be re-interpreted based on this new understanding. The Commander structure now paves the way towards a more complete molecular understanding of this essential complex and thereby of the core evolutionary conserved molecular mechanism of endosomal sorting.

### The structure and function of Retriever is distinct from the related Retromer complex

The Retromer complex composed of VPS35, VPS26A/B and VPS29 is a relatively well characterised complex that works in unison with different cargo adaptors including SNX3 and SNX27 for endosomal retrieval (80). Although it was known that Retriever shares similarities with Retromer, composed of the distantly related VPS35L and VPS26C subunits and the shared VPS29 protein and associates with at least one divergent adaptor SNX17 (2, 4, 6, 15), whether it was assembled or functioned in a similar way was unclear. Our studies show that it does indeed have an analogous architecture, with the central VPS35L protein providing an extended α-helical scaffold for the VPS26C and VPS29 subunits. However, Retriever has a much more compact structure, and most importantly has distinct conserved surfaces that mediate specific interactions with the CCC machinery and a sequence within its intrinsically unstructured N-terminal region that binds and regulates VPS29. This sequence mimics Pro-Leu-containing motifs found in previously characterised Retromer-interacting proteins, including the RAB7 GAP TBC1D5 (50), the RAB32/RAB38 GEF and SNARE trafficking protein VARP/ANKRD27 (49), and the secreted effector RidL from *Legionella pneumophila* (51, 53, 54). The sequence of VPS35L binds VPS29 with a high affinity that is enhanced by intramolecular tethering, and thus blocks Retriever-bound VPS29 from participating in these regulatory interactions. The incorporation of VPS29 into Retriever therefore is mutually exclusive of its ability to function in canonical Retromer-mediated transport. This also explains why synthetic macrocyclic peptides that target the conserved pocket on VPS29 can interact with Retromer but not Retriever (47). In human cells VPS29 is a highly abundant protein, typically present at up to twice the level of other Retromer subunits, and more than twenty times the concentration of other Commander subunits (81, 82). However, how the equilibrium between VPS29 association with either Retromer or Retriever is established and regulated remains an important question.

### The CCC complex is a unique assembly of enigmatic function

The structures presented here represent a significant advance in our understanding of the molecular interactions responsible for highly specific heterotypic interactions between the COMMD and CCDC subunits of the CCC and Commander complexes. The COMMD proteins have been shown to undergo homo- and heteromeric interactions using co-immunoprecipitation from cells (6, 16, 17, 21, 22, 28, 43–45, 71, 72), or following pairwise co-expression in bacterial systems (45, 71). The proteins also must form obligate dimers due to the distinct structure of their C-terminal COMM domains (45). However, in proteomics studies, including experiments that use COMMD family members as baits, the entire set of ten COMMD proteins are always identified in a complex together along with other Commander subunits (this study and (6, 16, 17, 20–22)). Altogether these studies strongly suggest that the COMMD subunits themselves likely exist in a co-assembled structural state. However, whether multiple distinct COMMD assemblies exist or whether they are all co-assembled in a single ten-subunit complex was unknown (15). Our biochemical, biophysical, and structural data confirms the latter, defining precisely how COMMD family members preferentially interact with each other in specific heterodimeric partnerships and provides a detailed understanding of their complete assembly into a ten-subunit heterodecameric ring. Furthermore, reconstitution of the whole CCC complex indicates that stable decameric ring formation is entirely dependent on interaction with the CCDC22 and CCDC93 subunits.

The structures of the COMMD and CCDC proteins demonstrate how COMMD proteins form a singular decameric complex in conjunction with stabilising sequences from the CCDC proteins and provide an unambiguous picture of how multiple inter-subunit contacts contribute to the specificity of the ring-shaped assembly. Although there are many individual requirements for precise COMMD interactions, some general findings are that the β1-β2 loop within the COMM domain of each COMMD subunit docks closely into a complementary pocket formed by the HN domain and linker of its cognate dimeric partner. These interfaces each involve a network of interactions that encompass a strictly conserved tryptophan within the C-terminal α7-helix of the neighbouring subunit in the ring. As one example, the HN domain of COMMD10 engages the β1-β2 loop of its dimeric partner COMMD5, together these interact with the Trp139 sidechain of COMMD7, and this network is stabilised by the presence of the C-terminal α7-helix of COMMD9. The one exception to this in the human complex is COMMD6 which lacks the HN domain, although the domain is found in orthologues of most other species.

One functional implication for the structural organisation of the COMMD assembly is that the HN domains are positioned peripherally and therefore appear to be primed for mediating potential specific intermolecular interactions. It has previously been proposed that the sequence divergence in the N-terminal HN domains could allow for distinct interactions amongst the family members, for example with the NF-κB complex (37, 44), and actin and microtubule cytoskeletal components (83). More generally, while many different protein families have evolved the ability to form multimeric ring structures, this most commonly involves homomeric symmetric interactions with one or a few repeated protein subunits (84–86). Heteromeric pseudo-symmetric structures composed of multiple paralogous proteins as seen for the COMMD decamer are less common, and typically arise from multiple gene duplication events. Some prominent examples include the eukaryotic 20S proteosome (87), the Sm/Lsm RNA-binding proteins (88–90), the CMG helicase (91–93) and origin recognition complex (94, 95) involved in initiating DNA replication. Identifying whether specific functional interactions are mediated by the different COMMD subunits remains an exciting avenue of investigation.

One important finding is that stable COMMD assembly requires the highly specific intercalation of CCDC22 and CCDC93 linker sequences between the N-terminal CH domains and C-terminal coiled-coil regions. These linkers make extensive contacts with different COMMD subunits to tie the assembly together. Two particularly interesting contacts are mediated by CCDC22 sequences with a particular P-x-x-R turn structure that interacts with the HN domains of COMMD5 and COMMD3. Although speculative, it raises the possibility that other HN domains could mediate binding of proteins with related sequences. In addition to stabilising the COMMD ring, we also find that central coiled-coil regions of the CCDC22-CCDC93 dimer mediate recruitment of the peripheral subunit DENND10. As the name suggests DENND10 (also called FAM45A) is a member of the DENN domain family which are generally thought to act as GEFs for RAB GTPases (96–99). It is localised to late endosomes and its knockdown perturbs aspects of endosomal morphology and function (21, 75), although it appears dispensable for assembly of other Commander subunits (21). To date only one crystal structure of a DENN domain in complex with a RAB GEF substrate has been determined, DENDD1B with RAB35 (74), while the only other structures known are Longin and DENN domain-containing dimers of C9orf72/SMCR8 and Folliculin/FNIP2 that actually act as RAB GAP complexes (100–104). DENND10 is distinct from all of these, as it lacks key residues found in DENND1B required for GEF activity, and the interface with CCDC22-CCDC93 would preclude RAB binding in the same region. Therefore, whether DENND10 has GEF (or GAP) activity, or plays a distinct role within the Commander complex remains to be determined.

### Assembly of the Commander holo-complex and its role in endosomal recycling

The experimentally validated model for the interaction between the C-terminal coiled-coil regions of CCDC22 and CCDC93 and VPS35L provides a molecular explanation for the coupling of the CCC and Retriever assemblies to generate the holo-Commander complex. Although this will require full structural confirmation, we show that mutations designed based on the AlphaFold2 modelling specifically block CCC and Retriever interaction. Our data thus supports the idea that while the CCC and Retriever assemblies can be considered as distinct stable structures, the function of these proteins likely depends on their incorporation into the Commander holo-assembly. The CCDC proteins share some structural similarity with other CH domain-containing and coiled-coil heterodimers (105), predominantly involved in regulating cytoskeletal interactions such as with ciliary microtubules (76, 77, 106), within dynein-adaptor assemblies (107–109), or at the kinetochore (110), and CH domains are also often involved in direct interactions with both actin and microtubule filaments (111–114). Endosomal recycling by Commander requires dynamic organisation of actin-rich retrieval domains on the endosomal surface (5). Endosome-associated branched actin is nucleated by Arp2/3 following activation by the Wiskott-Aldrich syndrome protein and SCAR homologue (WASH) complex, which interacts with Commander subunits (5, 6, 13, 21, 30, 83). Given this functional connection, it is tempting to speculate that CH domain interactions could be important for establishing the actin-rich microdomains required for endosomal sorting (5).

The structure of the Commander complex has allowed us to map the locations of causative mutations for XLID and RSS developmental disorders, leading to altered craniofacial features, cerebellar hypoplasia, hypercholesterolemia and cardiovascular development (7–14). Most missense mutations map to key structural elements or inter-subunit interfaces, and those we have tested all result in significant loss in overall Commander protein levels. In contrast we found that two deletion and frameshift mutations in VPS35L proximal to the VPS29 binding site had a specific impact on VPS29 interaction, without seriously affecting overall Commander assembly. This shows that XLID/RSS mutations can lead to either overall Commander instability or loss of specific interactions within the complex and reaffirms the important role that VPS29 plays in Commander function. These clinical phenotypes confirm the important connection between Commander and the endosomal WASH complex, given that the only other known mutations leading to RSS are found in WASH subunits WASHC4/KIAA1033 (115) and WASH5C/Strumpellin (10). Beyond XLID and RSS developmental disorders, altered Commander function leads to hypercholesterolemia linked to reduced trafficking of LDLRs (7-10, 12-14), defective copper homeostasis (116, 117), and protection against myocardial infarction by affecting LDLR transport (28, 29). Commander is also required for cellular infection by HPV (6) and SARS-CoV-2 (33–36), antibody-dependent cellular phagocytosis of cancer cells (118), and biogenesis of melanosomes in pigmented cells (119).

### Limitations of this study

Our work provides the overall structural framework for analysing Commander function in a wide array of cellular and disease-associated processes. However, while the core structures of the CCC complex and the interaction of VPS29 with sequences from VPS35L have been determined at high resolution using both X-ray crystallography and cryoEM, the coiled-coil regions of the CCDC proteins remain unresolved by experimental methods, and cryoEM maps of Retriever have revealed its overall architecture but not its entire atomic structure. In addition, while the interactions of the CCC complex with DENND10 and with Retriever have been mapped and experimentally validated both *in vitro* and *in situ*, it will still be important to obtain high resolution experimental structures of these complexes in the future. Ultimately, purification and structural studies of the full sixteen subunit Commander holo-complex will be needed to provide a fully complete picture of this complex, to identify potential conformational rearrangements in its overall organization, and eventually to determine how it interacts and assembles on the endosomal membrane and with integral membrane cargos and adaptors.

## ACKNOWLEDGEMENTS

We thank the Wolfson Bioimaging Facility, the Bristol Proteomics Facility, and the GW4 Facility for High-Resolution Electron Cryo-Microscopy funded by the Wellcome Trust (202904/Z/16/Z and 206181/Z/17/Z) and BBSRC (BB/R000484/1) at the University of Bristol for their support, and the University of Bristol Advanced Computing Research Centre for the provision of high-performance computing. We acknowledge the Medical Research Council (MRC) - LMB Electron Microscopy Facility for access and support of electron microscopy sample preparation and data collection and thank Jake Grimmett and Toby Darling (LMB scientific computation) for their support. We acknowledge the Bio21 Mass Spectrometry and Proteomics Facility (MMSPF) for the provision of instrumentation, training, and technical support. We thank Dr. Alun Jones (IMB, UQ) for assistance with peptide mass spectrometry and acknowledge use of the University of Queensland Remote Operation Crystallization and X-ray (UQ ROCX) Facility and the assistance of Dr. Kasun Athukorala Arachchige for protein crystallisation. X-ray data were collected on the MX2 microfocus beamline at the Australian Synchrotron. We thank Dr Andrew Carter (MRC-LMB) for suggesting the biGbac system and helping to establish this in Bristol. P.J.C. is supported by the Wellcome Trust (104568/Z/14/Z and 220260/Z/20/Z), the MRC (MR/L007363/1 and MR/P018807/1), the Lister Institute of Preventive Medicine, and the award of a Royal Society Noreen Murray Research Professorship (RSRP/R1/211004). B.M.C. is supported by an Investigator Grant, Senior Research Fellowship and Project Grant from the National Health and MRC (APP2016410, APP1136021 and APP1156493). D.A.S. is supported by an Investigator Fellowship and Project Grant from the National Health and MRC (APP2009732 and APP1156732). K.E.M is supported by the Wellcome Trust through a Sir Henry Wellcome Postdoctoral Fellowship (220480/Z/20/Z). M.D.H. was supported by the AINSE PGRA, and R.B. and S.S. were supported by the EndoConnect European Research Training Network (No. 953489). C.S. and I.B. are Investigators of the Wellcome Trust (210701/Z/18/Z; 106115/Z/14/Z). T.A.W is supported by a Royal Society University Research Fellowship (URF/R/201024) and a grant from the Moore Foundation. E.D is funded by the MRC (MC_UP_1201/13) and Human Frontier Science Program (Career Development Award CDA00034/2017). We would also like to acknowledge and thank Milot Mirdita, Sergey Ovchinnikov, Martin Steinegger and the ColabFold team for making their AlphaFold2 modelling pipeline available for public use.

## AUTHOR CONTRIBUTIONS

M.D.H., K.E.M., D.A.S., B.M.C. and P.J.C. designed research; M.D.H., K-E.C., R.J.H., and R.G. performed X-ray crystallography, ITC analysis and AlphaFold2 modelling; K.E.M., and M.C. performed insect cell purification, K.E.M., V.J.P-H., S.K.N.Y., J.R., U.B., performed Retriever cryo-EM and data analysis; K.E.M., V.J.P-H., T.H.D.N. performed CCC complex cryo-EM and data analysis; K.E.M., R.B., K.K., S.S., J.S., C.M., C.S.P., T.I.C., N.J.C., A.S.N., and K.J.H. performed cell-based analysis and proteomics; E.R.R.M. and T.A.W. performed evolutionary analysis; K.E.M., S. Satoh., E.D., T.A.W., I.B., C.S., D.A.S., E.D., B.M.C. and P.J.C. acquired funding; and M.D.H., K.E.M., R.B., B.M.C. and P.J.C. wrote the paper guided by all authors.

## INCLUSION AND DIVERSITY

We support inclusive, diverse, and equitable conduct of research.

## STAR★METHODS

### KEY RESOURCES TABLE

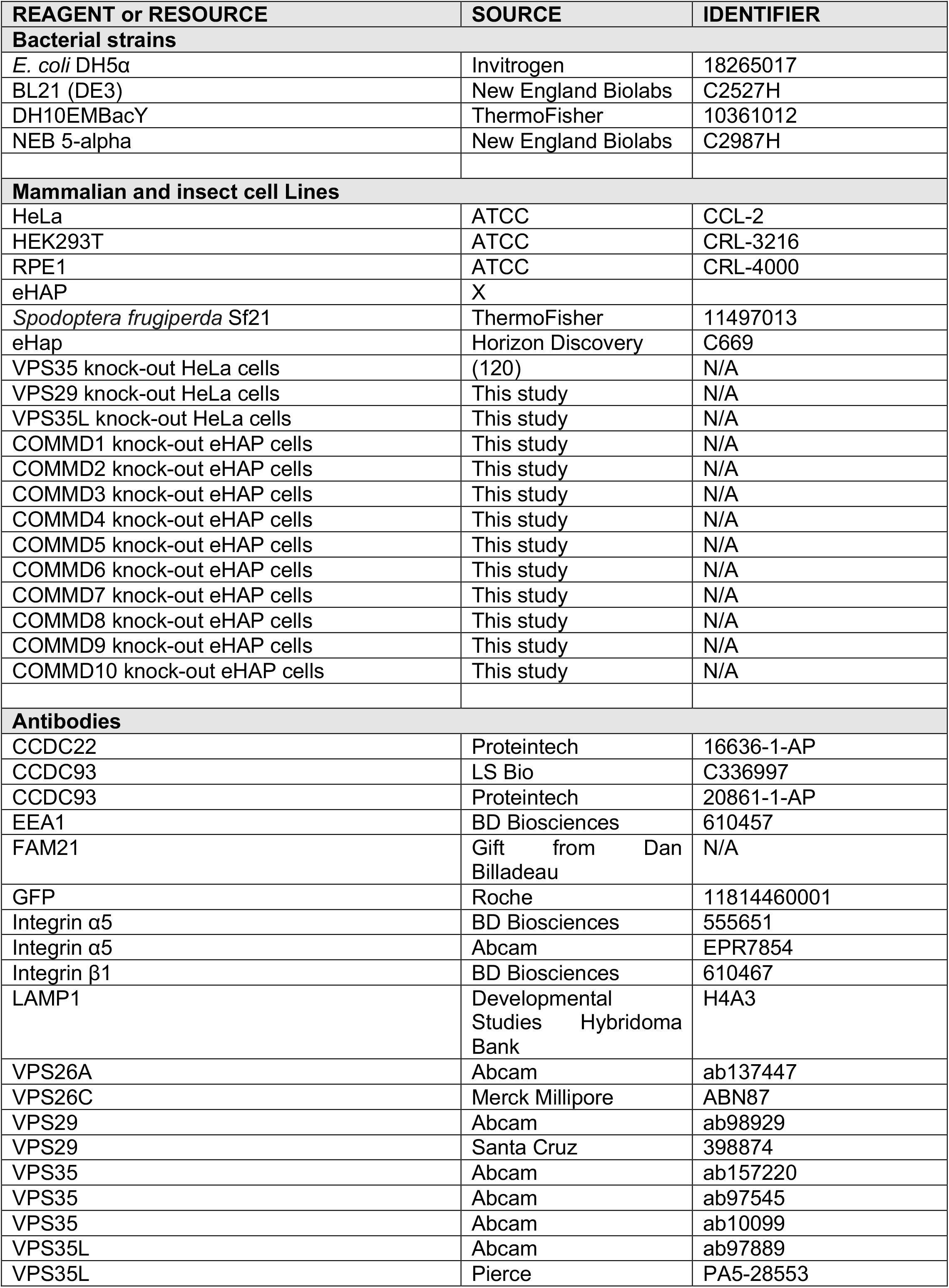
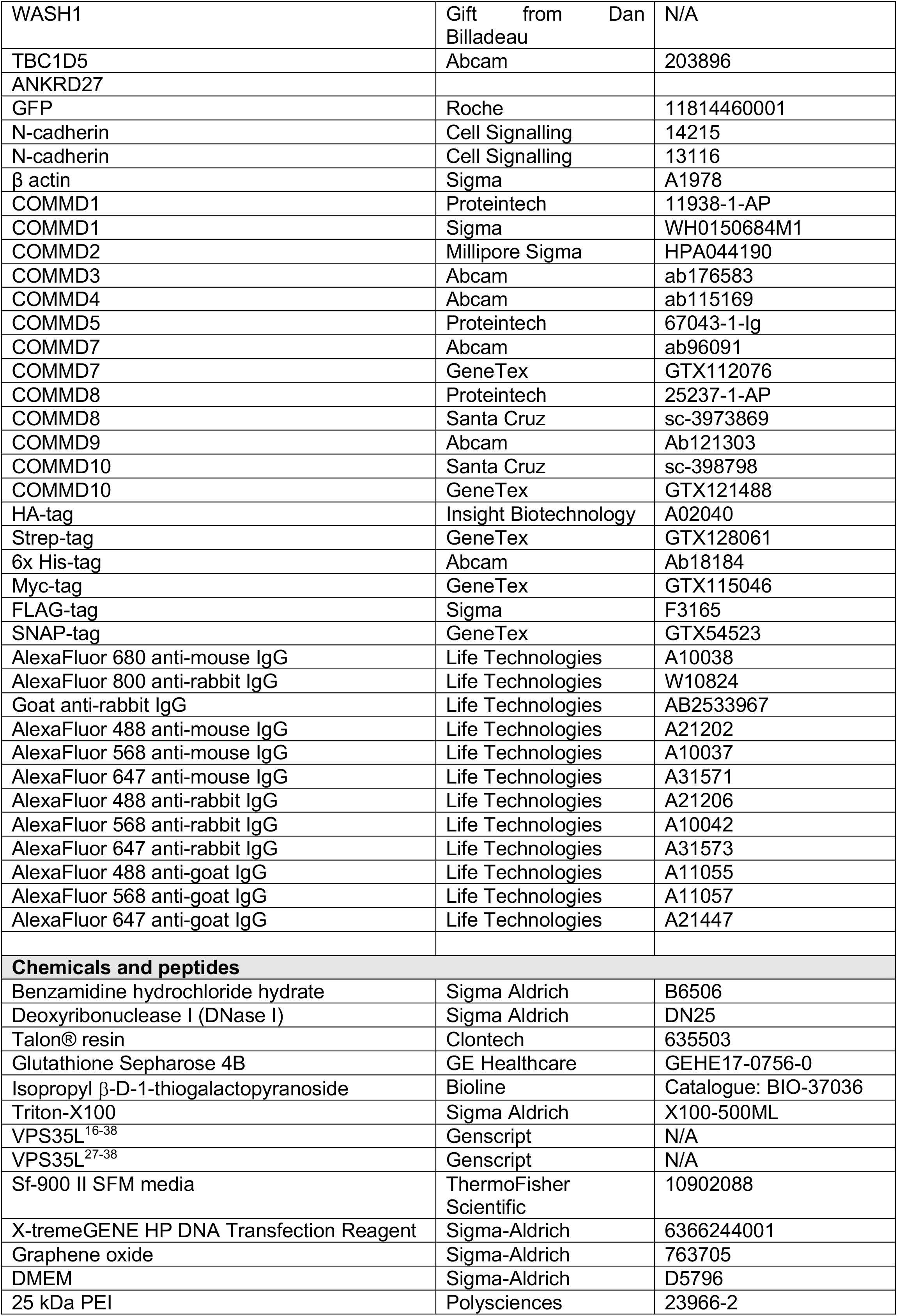
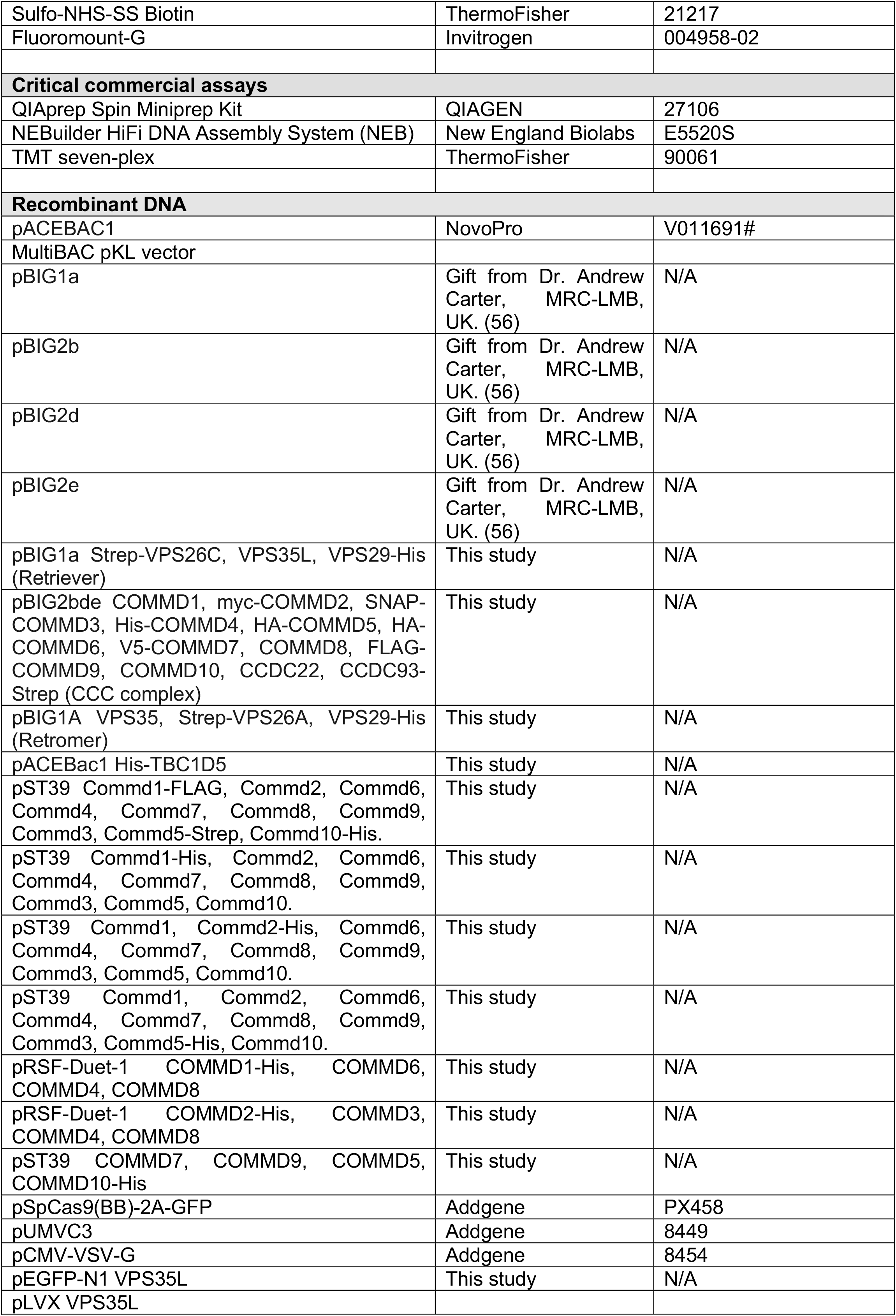
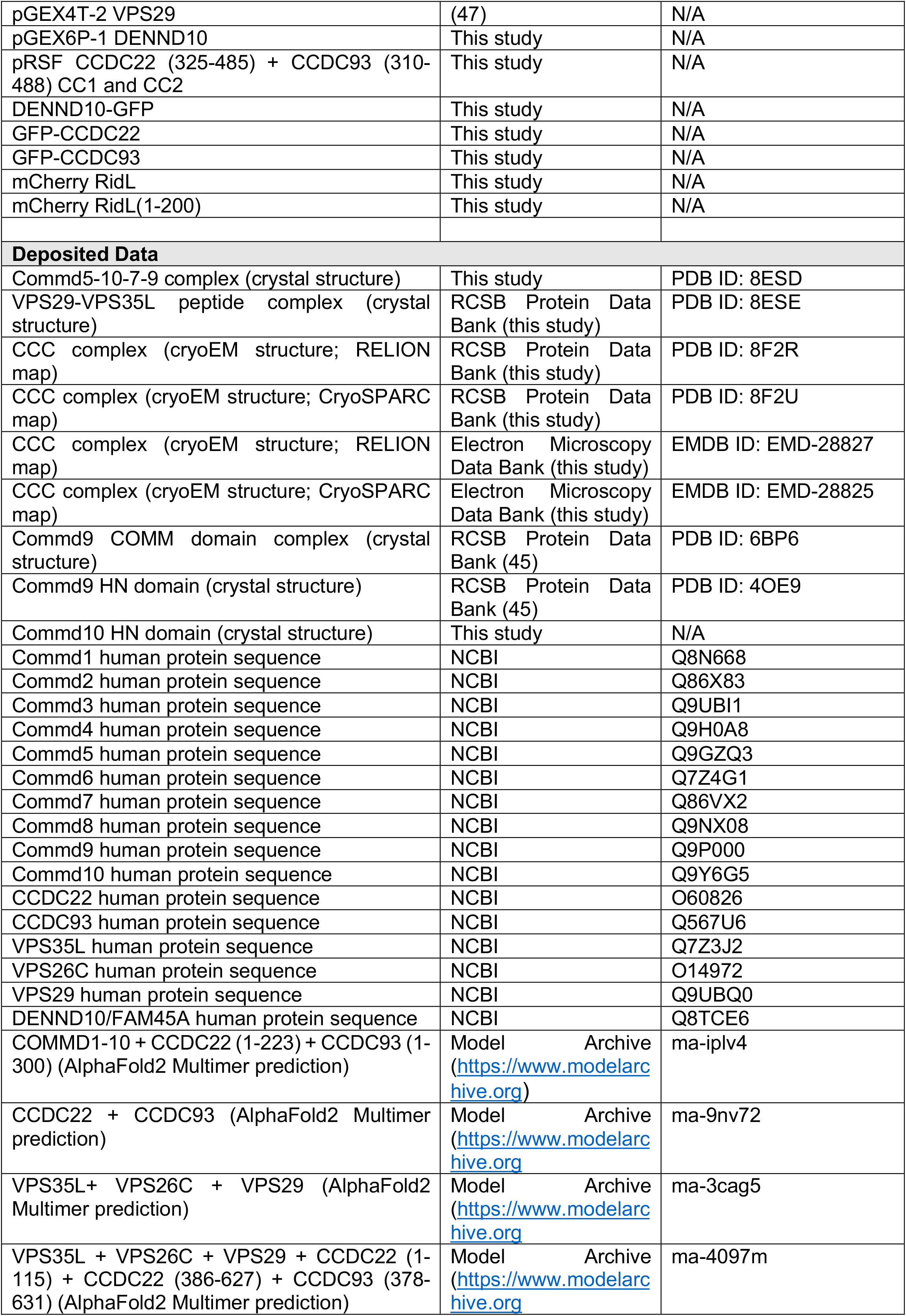
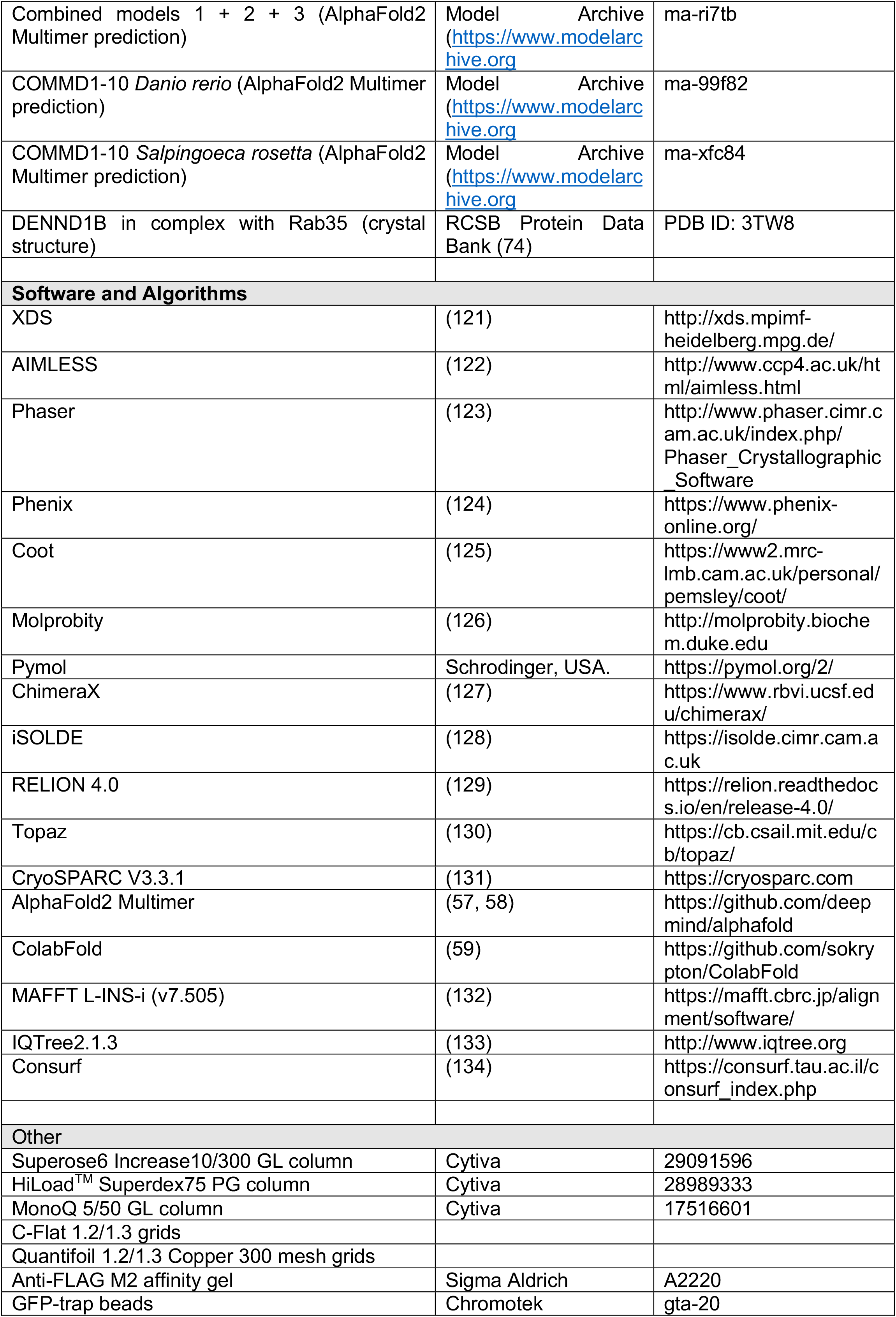
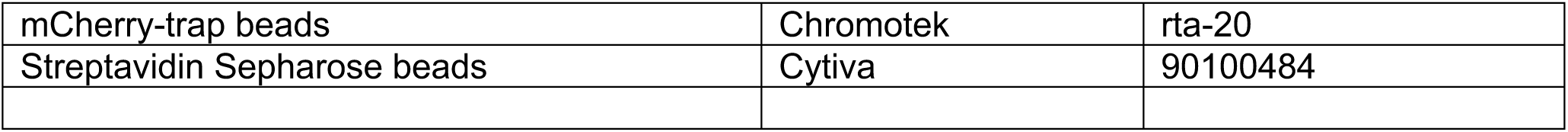

### RESOURCE AVAILABILITY

#### Lead Contact

Further information and requests for resources and reagents should be directed and will be fulfilled by the Lead Contacts, Prof. Brett Collins, Prof. Peter Cullen and Dr. Kerrie McNally.

#### Materials Availability

Plasmids generated in this study are available from the Lead Contacts with a completed Materials Transfer Agreement.

#### Data and Code Availability

- Coordinates for the COMMD1-10/CCDC22/CCDC93 complex have been deposited at the Protein Data Bank (PDB) with accession codes 8F2R (CryoSPARC) and 8F2U (Relion) with respective Electron Microscopy Data Bank under accession codes EMD-28825 (CryoSPARC) and EMD-28827 (Relion). Coordinates and structure factors for the crystal structure of the COMMD5-COMMD10-COMMD9-COMMD7 complex have been deposited at the PDB with accession code 8ESD. Coordinates and structure factors for the crystal structure of the VPS29-VPS35L peptide complex have been deposited at the PDB with accession code 8ESE.
- Predicted structures of the Commander complex and sub-assemblies using AlphaFold2 have been deposited in the ModelArchive (https://www.modelarchive.org) with accession numbers as outlined in **Table S8**.
- The data for the FLAG-tagged COMMD subunit mass spectrometry proteomics have been deposited in the ProteomeXchange Consortium via the PRIDE (135) partner repository with accession number XXXX.
- This paper does not report original code.
- Any additional information and all the relevant raw data required to reanalyze the data reported in this paper is available from the lead contact upon request.

### EXPERIMENTAL MODEL AND SUBJECT DETAILS

*Escherichia coli* BL21 (DE3) cells were used for the overexpression of native recombinant proteins. Cells were grown at 37°C and protein expression was induced with 0.8 mM ispropylthio-β-galactoside (IPTG) before the temperature was reduced to 21°C and cultures were allowed to grow for 18 h. HeLa and RPE1 cells were maintained in DMEM (D5796; Sigma-Aldrich) plus 10% fetal calf serum (F7524; Sigma-Aldrich) under standard conditions. These cell lines were obtained from America Type Culture Collection (ATCC). Parental and stable cells lines were negative for mycoplasma by DAPI staining. *Spodoptera frugiperda* Sf21 cells (Cat no. 11497013, ThermoFisher Scientific) for baculoviral expression of recombinant proteins in insect cells were grown at 26°C in Sf-900 II SFM media (Cat no. 10902088, ThermoFisher Scientific).

### METHOD DETAILS

#### Antibodies

The following primary antibodies used in this study are found in **Table S3**, along with their dilution and usage. Secondary antibodies used in this study are found in **Table S4**.

#### Cloning with biGBac plasmids

Retriever and Retromer: Genes for human Retriever and Retromer were codon optimized for *Spodoptera frugiperda* and synthesized by Twist Biosciences (San Francisco, CA). Codon optimised genes were cloned into pACEBAC1. Retromer and Retriever were assembled through Gibson assembly into pBIG1A (empty pBIG plasmids were gifts from Dr Andrew Carter, MRC-LMB, Cambridge, UK) (136). For TBC1D5 expression, the coding region of TBC1D5 was subcloned from pEGFP-C1 TBC1D5 into pACEBac1. CCC complex: Genes for human CCC complex expression were codon optimized for *Spodoptera frugiperda* while avoiding the introduction of *Pme*I, *Swa*I, *Bam*H1 and *Hind*III restriction sites and synthesized by Twist Biosciences (San Francisco, CA). Codon optimised sequences were cloned into the MultiBAC pKL transfer plasmid at the *Bam*HI and *Hind*III sites within the multiple cloning site. The CCC complex genes were assembled using the biGBac cloning strategy (55, 56). Plasmid DNA was prepared using a QIAprep Spin Miniprep Kit according to the manufacturer’s protocol (Cat no. 27106, Qiagen).

#### Bacmid purification

pACEBac1 or pBIG vectors were transformed into DH10EMBacY competent cells which contain a modified baculoviral genome (55). Transformations were left to recover overnight before being plated onto agar plates containing 50 µg/ml kanamycin, 10 µg/ml tetracycline, 7 µg/ml gentamycin, 40 µg/ml Isopropyl β-d-1-thiogalactopyranoside (IPTG) and 100 µg/ml Blue-Gal (Cat no. 15519028, ThermoFisher Scientific). The multigene transfer vector integrates with the baculoviral genome via Tn7 transposition. White colonies were grown overnight in 2 ml of LB supplemented with 50 µg/ml kanamycin, 10 µg/ml tetracycline, 7 µg/ml gentamycin. Bacmid DNA was prepared using buffers from a QIAprep Spin Miniprep Kit (Cat no. 27106, Quiagen) according to the MultiBac protocol.

#### Baculovirus generation

Sf21 cells were seeded at 1×10^6^ cells/well in a 6-well plate in a total volume of 3 ml of Sf-900 II SFM media (Cat no. 10902088, ThermoFisher Scientific). Bacmid DNA was transfected into Sf21 cells using X-tremeGENE HP DNA Transfection Reagent (Cat no. 6366244001, Sigma-Aldrich) according to the manufacturer’s protocol and incubated at 26°C for 72 h. The media from the transfected culture was used to infect a 25 ml suspension culture of Sf21 cells at 1×10^6^ cells/ml. At 48 h post proliferation arrest the V1 generation of virus was harvested by pelleting the cells at 2000 rpm for 10 min and collecting the supernatant. To amplify the infectivity of the virus, V1 was added to a culture of Sf21 cells and supernatant harvested - termed V2. All viruses were stored at 4°C in the dark.

#### Protein expression in insect cells

For protein expression, the V1 or V2 virus were used to infect suspension cultures of Sf21 insect cells in Sf-900 II SFM media (Cat no. 10902088, ThermoFisher Scientific). Cells were seeded at 0.6×10^6^ cells/ml in 2 L Erlenmeyer shaker flasks in a total volume of 600 ml. At a density of 1×10^6^ cells/ml, 6 ml of V1 or V2 was added to the culture. At 48 h post proliferation arrest cells were harvested by centrifugation at 2000 rpm for 10 min. Cell pellets were either immediately used for protein purification or stored at −20°C.

#### Retriever and TBC1D5 purification from insect cells

The insect cell pellets were resuspended in lysis buffer (25 mM HEPES pH 8.0, 300 mM NaCl, 2 mM β-mercaptoethanol, EDTA-free protease inhibitor tablets (A32965, Pierce)) and lysed on ice using a 130-Watt Ultrasonic Processor (UY-04714-51, Cole-Parmer) for a total of 2 min 30 s using a 10 s on 30 s off cycle. Lysates were cleared by centrifugation at 4°C for 30 minutes at 18,000 x g. His TALON resin was used to purify his-tagged proteins. Purification was performed at 4°C. TALON resin was equilibrated with lysis buffer (25 mM HEPES pH 8.0, 300 mM NaCl, 2 mM β-mercaptoethanol). Cleared cell lysate was then added to the column and allowed to flow through the TALON resin. Once the lysate had completely flowed through the column, the column was thoroughly washed in 10x CV lysis buffer, followed by 10x CV wash buffer (25 mM HEPES pH 8.0, 300 mM NaCl, 2 mM β-mercaptoethanol and 20 mM imidazole). His-tag proteins were eluted from the column by elution buffer (25 mM HEPES pH 8.0, 300 mM NaCl, 2 mM β-mercaptoethanol and 200 mM imidazole). Size exclusion chromatography (SEC) was performed at 4°C using an ÄKTA prime and purifier system (GE Healthcare). A Superdex200 size exclusion column 10/300 GL (GE healthcare, catalogue number 28990944) was equilibrated in SEC buffer (25 mM HEPES pH 8.0, 300 mM NaCl, 2 mM β-mercaptoethanol). Protein was injected onto the column and 0.5ml fractions were collected.

### Retromer purification from insect cells

Retromer was purified using the same method as Retriever/TBC1D5 but using different buffers – lysis buffer: 25 mM HEPES pH7.5, 150 mM NaCl, 2 mM β-mercaptoethanol, EDTA-free protease inhibitor tablets (A32965, Pierce); wash buffer: 25 mM HEPES pH 7.5, 150 mM NaCl, 2 mM β-mercaptoethanol and 20 mM imidazole; elution buffer: 25 mM HEPES pH 7.5, 150 mM NaCl, 2 mM β-mercaptoethanol and 200 mM imidazole; SEC buffer: 25 mM HEPES pH 7.5, 150 mM NaCl, 2 mM β-mercaptoethanol.

#### CCC complex purification from insect cells

Insect cell pellets were resuspended in 5x volume of lysis buffer (50 mM HEPES pH7.2, 150 mM NaCl, 2 mM β-mercaptoethanol, 0.1% Triton-X100 with EDTA-free protease inhibitor tablets (A32965, Pierce). Lysates were sonicated on ice using a 130-Watt Ultrasonic Processor (UY-04714-51, Cole-Parmer) for a total of 2 min 30 s using a 10 s on 30 s off cycle. Lysates were cleared by centrifugation at 20,000 rpm for 25 min at 4°C. Cleared lysates were loaded onto a Econo-Pac Chromatography Column (Cat no. 7321010, Bio-Rad) packed with 1 ml Steptactin resin (2-1201, IBA Lifesciences) pre-equilibrated in lysis buffer. The column was washed with 2 x 25 ml lysis buffer and bound protein eluted using 5 x 1 ml lysis buffer plus 2.5 mM desthiobiotin. A subset of rotein containing fractions were crosslinked with 1 mM BS3 (11841245, Thermo Scientific) for 2 h at 4°C. The reaction was quenched using 1 M Tris, pH 7.5 at a final concentration of 50 mM. Crosslinked and non-crosslinked protein containing fractions were gel filtered using a Superose 6 10/300 GL size exclusion column (Cat no. 29091596, GE Healthcare) attached to an ÄKTA pure chromatography system (GE Healthcare) pre-equilibrated in buffer containing 50 mM HEPES pH7.2, 150 mM NaCl, 2 mM β-mercaptoethanol, 0.01% (v/v) Triton-X100. Fractions of 500 µl were collected and analyzed. All purifications steps were performed at 4°C and samples kept on ice.

#### Native PAGE

Samples were prepared in a 1X dilution of Novex Tris-Glycine Native Sample Buffer (2X) (LC2673, ThermoFisher Scientific). Samples were separated by native polyacrylamide gel electrophoresis (native-PAGE) on a Novex WedgeWell 8 to 16% Tris-Glycine mini protein gel (XP08162BOX, Invitrogen). A 1X running buffer was prepared using (10X) Novex Tris-Glycine Native Running Buffer and used to fill the chamber of Invitrogen Mini Gel Tanks (A25977, Invitrogen). Typically, 20 µg of protein was loaded per well along with 4 µl of NativeMark™ Unstained Protein Standard (LC0725, ThermoFisher Scientific) in one lane as a molecular weight marker. Following PAGE the gel was washed in ddH_2_0 for 5 min. To visualise the proteins the gel was immersed in Coomassie stain, made with 0.1% Coomassie Brilliant Blue R-250 (B7920, Sigma-Aldrich), 40% methanol, 10% acetic acid and filtered through a Whatman No. 1 filter. The gel submerged in Coomassie stain was heated in a microwave for 30 s and allowed to incubate for 2 min with gentle agitation. The stain was removed and rinsed with ddH_2_0 before immersing in de-stain (20% methanol (v/v), 10% acetic acid (v/v) in ddH_2_0) and microwaving for 30 s. The gel was incubated in de-staining solution until bands could be distinguished from background stain. Gels were visualised with an Odyssey infrared imaging system (LI-COR Biosciences).

#### Negative stain EM

5 μg of Retriever or the CCC complex were placed onto carbon-coated pioloform copper-mesh grids and incubated for one minute. After the incubation, the excess protein solution was blotted and the grid was washed quickly in 4 μl 3% uranyl acetate, blotted again and then incubated with 4 μl of 3% uranyl acetate for one minute. After the uranyl acetate incubation, the grids were blotted, washed a third time in uranyl acetate before blotting dry and left to air dry. Images were recorded on a 200-kV Tecnai F20 microscope (FEI) equipped with a FEI Ceta 4k x 4k charge-coupled device camera at 68,000 magnification corresponding to a pixel size of 1.63 Å/pixel. A total of 14,000 particles from 280 images were picked and reference free two-dimensional classification was performed with RELION 3.2.

#### Graphene oxide coating of Quantifoil 1.2/1.3 Copper Mesh grids

Graphene oxide coated grids were prepared the day before use. Quantifoil 1.2/1.3 Copper 300 mesh grids were glow discharged for 1 minute using an Edwards S150B Sputter coater discharger at power level 7, 40 mA. Graphene oxide (Sigma, 763705, 2 mg/ml in H_2_0) was freshly diluted 1/10 to 0.2 mg/ml in MilliQ water. The diluted graphene oxide solution was span at 600 x g until the visible flakes pelleted. 3 µl of span graphene oxide solution, taken from the top, was applied to the glow-discharged grids and incubated for 1 minute. After incubation, grids were blotted with Whatman No.1 filter paper and washed/blotted twice with 20 µl of MilliQ H_2_O and a final wash with 20 µl of MilliQ H_2_0 was applied to the bottom of the grid. Grids were air-dried overnight.

#### Cryo-EM grid preparation

4 µl of ∼ 0.2 mg/ml purified Retriever was vitrified in ethane-propane on glow-discharged C-Flat 1.2/1.3 grids using a FEI Vitrobot. Grids were screened for suitable ice using a 200kV Talos Arctica equipped with energy filter and *a* Gatan K2 direct electron detector. 3 µl of ∼0.1 mg/ml purified, cross-linked human CCC complex was applied onto a graphene oxide coated Quantifoil 1.2/1.3 Copper 300 mesh grid. The grids were not glow discharged before sample application. Sample was incubated on the grid for 30 seconds prior to blotting (3-3.5 seconds at blot force −15) and vitrification in liquid ethane using a FEI Vitrobot Mark IV. Grids were screened for suitable ice and particle distribution using a 200 kV ThermoFischer Glacios equipped with Falcon III direct electron detector. **Cryo-EM data collection** Data collection of the Retriever complex was performed on at 200 kV Talos Arctica equipped with energy filter and Gatan K2 detector. EPU software (ThermoFischerScientific) was used for automated data acquisition. Data was collected in super-resolution mode with a virtual pixel size of 0.525 Å per pixel. 4,862 movies were collected, with a dose rate of 61.7 e-/Å^2^. Data collection of the CCC complex was performed on a 300 kV ThermoFischerScientific Titan Krios transmission electron microscope with a Falcon IV direct electron detector. EPU software (ThermoFischerScientific) was used for automated data acquisition. Data was collected from 2 independently prepared grids, in 2 separate sessions. The Falcon IV detector was used in counted mode with a pixel size of 1.084 Å per pixel for both data collections. The first dataset consisting of 2871 movies was collected with a total dose 43.55 e-/Å^2^ over a total exposure time of 9.01 seconds. The second dataset, consisting of 2779 movies was collected with a total dose of 42.73 e-/Å^2^ over a total exposure time of 9.01 seconds. A range of defocus values (−1.2, −1.4, −1.6, −1.8, −2.0 µm) were used for collection of both datasets. Movies on the Titan Krios were collected in EER format.

#### Cryo-EM data processing Retriever

Movies were imported into RELION 4.0 (129). Using the motion-correction program implemented within RELION, movies were dose-weighted, drift-corrected, gain-corrected and summed into single micrographs. A binning factor of 2 was used during motion correction, resulting in motion-corrected micrographs with a pixel size of 1.05 Å. CTFFIND-4.1, integrated within RELION, was used to estimate the contrast transfer function (CTF) parameters for the motion-corrected micrographs. Micrographs with high astigmatism values or crystalline ice were removed from the dataset.

Retriever particles were manually picked, with a particle diameter of 180 Å until ∼3000 particles had been picked. The manually picked particles were used to train Topaz (130) which is implemented within RELION. Particles were then autopicked using the Topaz trained network. Particles with a figure-of-merit (FOM) value above −3 were extracted using a box size of 240 pixels. We then performed 2 rounds of reference-free 2D classification. Following 2D classification, we only discarded classes that did not look like they contained protein particles (*e.g.* ice crystals). We kept all classes that looked to contain protein particles as we did not want to lose rare orientations of the particles during the 2D classification steps. 252,548 particle stacks were imported into CryoSPARC V3.3.1 (131). Initial 3D reconstructions were generated using an *ab initio* job with 4 classes. The resulting four 3D initial reconstructions were then used as templates in a heterologous refinement with 4 classes. The map quality of resulting classes from the heterologous refinement were assessed in Chimera. One class, with 119,564 particles, was selected for further processing. This class was refined using homogenous refinement followed by NU (Non-uniform) refinement. Gold-standard Fourier Shell Correlations (FSCs) and directional FSC plots and measurements of sphericity (calculated using the online 3DFSC tool) indicate that the final map has an overall resolution of ∼4.2 Å. However, this is an overestimation of the resolution of the map, as it was very clear from the reference-free 2D classification classes and 3D refinement jobs that Retriever particles displayed preferred orientation. The final CryoSPARC refined map does not contain particles observed from all angles, has a large deviation from the directional FSC and displays low sphericity (0.715 out of 1) (**Fig. S1I, S1J**). The map is therefore insufficient for model building and refinement.

We tried unsuccessfully to overcome these preferential orientation issues by changing the sample prep, including but not limited to; changing buffer composition and pH; addition of detergents; protein concentration; grid type (Quantifoil copper/gold, lacey, hole size); grid support (carbon, graphene oxide); freezing conditions (blotting time, humidity, plasma cleaning time, incubation time, blotting equipment (FEI Vitrobot, Leica EM GP)). We also collected a tilted dataset but data processing with this dataset was unsuccessful (data not shown). We also tried several methods of data processing in RELION/CryoSPARC to recover less frequent orientations. However, these were unsuccessful at improving the distribution of views and the overall final map (data not shown).

#### Cryo-EM data processing the CCC complex

Data processing of the CCC complex micrographs was first performed in CryoSPARC V3.3.1. The two movie datasets from the two independent grids were processed independently until indicated. Movies were imported into CryoSPARC and were processed using CryoSPARC’s patch motion correction. CTF estimation was achieved using CryoSPARC’s patch CTF. Particles were initially picked using the blob picker tool with a minimum particle diameter of 100 Å and a maximum particle diameter of 180 Å. Picked particles were extracted and underwent several rounds of reference-free 2D classification. Good 2D classes were selected and used as templates for a further round of particle picking using CryoSPARC’s termplate picker tool. These newly picked particles were extracted and underwent several rounds of 2D classification and class selection to discard any ‘bad particles’. The best 2D classes were then used to generate *ab initio* reconstructions of the complex. For the first dataset, consisting of 2,779 micrographs, 198,705 particles were used to generate 2 *ab inito* reconstructions. For the second dataset, consisting of 2,872 micrographs, 151,195 particles were used to generate 4 *ab initio* models. The *ab initio* models from each respective dataset were then used as templates in a hetero refinement job using the same particles as the *ab inito* job. Following hetero refinement, maps were opened in chimera and the best 3D classes were selected for further refinement using homo refinement and in the case of the first dataset, a further step of NU refinement. The resulting 2 refined reconstructions from the 2 independent datasets (consisting of 138,424 and 55,894 particles respectively) were then used to create templates for a further round of particle picking, using the create template tool in CryoSPARC. Particles were picked using the templates created from their respective dataset and extracted. Particles picked and extracted from the second dataset (374, 899) were classified using 2D classification and good classes (209,507 particles) were used to train Topaz. This Topaz trained model was then used to pick particles in RELION4.0 (see below). The extracted particles from each independent dataset were combined to give a total of 2,063,629 particles. Good classes of particles were selected following several rounds of 2D classification. 4 *ab initio* models were generated from 333,394 particles and these models were used as templates for a subsequent hetero refinement. The classes were opened in Chimera and the best class consisting of 153,104 particles was chosen for homo refinement followed by NU refinement to give a final reconstruction with an overall resolution of 3.1 Å. Some regions of the map were less well resolved than the central core domain (which was very well resolved in the CryoSPARC reconstruction) probably due to flexibility of these domains relative to the core. To better resolve the flexible regions of the map, we processed the data in RELION 4.0. The two independent datasets were processed analogously in RELION but kept independent until stated. Micrographs were imported into RELION, dose-weighted, drift-corrected, gain-corrected and summed into single micrographs using the motion-correction program implemented within RELION. CTF estimation was done using CTFFIND-4.1 within RELION. Particles were autopicked using the Topaz implementation within RELION, using the Topaz trained model from the CryoSPARC processing (see above). Particles with a figure of merit above −1 were extracted with a box size of 264 and then binned by a factor of 4. Particles underwent one round of 2D classification, from which the best classes were selected and used in a 3D classification job with 6 classes. The final CryoSPARC map of the CCC complex (described earlier) was binned, imported into RELION and used as an initial model in these 3D classification jobs. A T parameter of 4, 40 iterations and angular sampling of 7.5 degrees were used for the 3D classifications. 3D classes were opened in Chimera and the best 3D class was selected. Particles in this class were re-extracted without binning and then the selected particles from each independent dataset were combined to give a total of 166,533 particles which were then refined to 4.2 Å. The particles then were subjected to one consecutive rounds of CTF refinement (beam tilt, anisotropic magnification and defocus), followed by Bayesian polishing (137) and 3D refinement. This further improved the global resolution of the map to 4.1 Å. To improve the resolution of the densities of the flexible HN domains, the angular assignments from the latest refinement were used for a round of alignment-free 3D classification, with a T parameter of 16. The best 3D class (20,034 particles), which contained densities for all the HN domains as well as the CH domain of CCDC93, was selected for a further round of 3D refinement, resulting in a 3.8 Å reconstruction, which was further improved to 3.5 Å following a final round of CTF refinement and 3D refinement.

#### Model building and refinement of the CCC complex

The AlphaFold2 model of the COMMD1-10, CCDC22 (1–260), CCDC93 (1–306) complex (see below) was docked into the sharpened RELION 4.0 (129) map using Chimera X (127). The docked model was then passed through PHENIX (138) real-space refinement to correct Ramachandran outliers and bond angles. To improve the stereochemistry and clash score this model was refined with the Chimera X plugin iSOLDE (128) and then subjected to several rounds of further refinement and rebuilding using a combination of PHENIX (138), COOT (139) and iSOLDE (128) resulting in a final model with excellent refinement statistics and stereochemistry based on Molprobity scores. All images were rendered using ChimeraX (127).

#### Molecular biology and cloning for *E. coli* expression

A series of gene cassettes codon optimised for bacterial expression were synthesised by the Gene Universal Corporation (USA). These were then progressively cloned into the pST39 vector to allow co-expression of all ten human COMMD family members. Briefly, gene cassettes were as follows cassette 1 (*Xba*I): COMMD1, COMMD2, COMMD6; Cassette 2 (*Hind*III): COMMD4; Cassette 3 (*Eco*RV): COMMD7, COMMD8, COMMD9; Cassette 4 (*Nru*I): COMMD3, COMMD5-Strep, COMMD10-His. Each protein contained a 5’ ribosomal binding site and a 3’ stop codon and were expressed off a single T7 promoter allowing the simultaneous expression of each of the ten COMMD proteins individually. The order of COMMD proteins was determined due to the internal restriction site and the required restriction sites for vector linearisation and cloning. A series of four vectors were then developed from this template to only have a singular His tag on COMMD1, 2 and 5. Subsequent pST39 co-expression vectors encoding four proteins from subcomplex B (COMMD5-SBP-7-9-10-His) were synthetically generated by Gene Universal using the same codon optimised gene sequences. DENND10 was synthesized and codon optimized for *E.coli* expression by geneuniversal and subcloned into a pGEX6P-1 vector (BamHI). CCDC22 (325–485) and CCDC93 (310–488) were synthesized by geneuniversal and subcloned into a pRSFduet vector at BamHI and NdeI, respectively.

#### Protein expression and purification from *E. coli*

The bacterial expression plasmids were transformed into *Escherichia coli* BL21 DE3 competent cells (New England Biolabs) and plated on agar plates containing ampicillin. Clones from this agar plate were collected and grown overnight in 10 mL LB broth, overnight “starter” cultures were expanded into 1 L cultures. Cultures were grown until reaching OD_600_ reached 0.8 and induced with 0.8 mM isopropylthio-β-galactoside (IPTG). Cultures were then cooled to 20°C and allowed to grow for ∼16 h. Cells were harvested by centrifugation at 6000 x g for 5 min at 4°C and the harvested cell pellet was resuspended in lysis buffer (50 mM HEPES pH 7.4, 500 mM NaCl, 5 mM Imidazole, 10% glycerol, 1 mM n-Dodecyl-β-D-Maltopyranoside (DDM), 50 μg/mL benzamidine, 100 units of DNaseI, and 2 mM β-mercaptoethanol). If cells were to be stored for later purification, they were flash frozen in liquid nitrogen and stored at −80°C. Fresh or thawed cells were lysed using sonication and the lysate was clarified by centrifugation at 50,000 x g for 30 min at 4°C. Complexes containing His-tagged COMMD2 or COMMD10 were purified on a Talon resin (CloneTech) gravity column and eluted using 500 mM imidazole in a buffer containing 500 mM NaCl, 10% glycerol, and 2 mM β-mercaptoethanol. Complexes containing FLAG-tagged COMMD1 were purified using Anti-FLAG M2 affinity gel (Sigma Aldrich) and eluted using a genetically engineered TEV cleavage site. Eluted proteins were subsequently passed through a Superdex 200 10/300 column attached to an AKTA Pure system (GE healthcare) in 50 mM HEPES pH 7.4, 500 mM NaCl for crystallisation, and isothermal titration experiments; 50 mM Tris pH 7.4, 500 mM NaCl for mass photometry. For further analysis by MS COMMD complexes isolated using His or FLAG tags were passed through SEC in 50 mM Tris pH 7.4 and 30 mM NaCl and further purified using a monoQ anion exchange chromatography column with a gradient running from 30 mM NaCl to 500 mM NaCl. Wild type and mutant complexes of CCDC22 (325–485) and CCDC93 (310–488) contained a aminoterminal decaHis tag on CCDC93 and was co-expressed off the pRSF-duet vector, expression and purification were as above. DENND10 was expressed as above and purified in the same manner except talon resin was substituted for glutathione resin (Clonetech).

#### uHPLC, Mass spectrometry and protein identification

COMMD complexes purified by size exclusion chromatography and anion exchange chromatography were trypsinised at a ratio of 1:100 (trypsin:protein) overnight at 37°C and analysed by uHPLC-MS/MS on an Eksigent, Ekspert nano LC400 uHPLC (SCIEX, Canada) coupled to a Triple TOF 6600 mass spectrometer (SCIEX, Canada) equipped with a duo microelectrospray ion source. 5 µl of each extract was injected onto a 300 µm x 150 mm ChomXP C18 CL 3 μm column (SCIEX, Canada) at 5 µl/min. Linear gradients of 2-25% solvent B over 35 min at 5 μL/minute flow rate, followed by a steeper gradient from 25% to 60% solvent B in 15 min were used for peptide elution. The gradient was then extended from 60% solvent B to 80% solvent B in 2 min. Solvent B was held at 80% for 3 min for washing the column and returned to 2% solvent B for equilibration prior to the next sample injection. Solvent A consisted of 0.1% formic acid in water and solvent B contained 0.1% formic acid in acetonitrile. The ionspray voltage was set to 5500V, declustering potential (DP) 90V, curtain gas flow 25, nebuliser gas 1 (GS1) 13, GS2 to 15, interface heater at 150°C and the turbo heater to 150°C. The mass spectrometer acquired 250ms full scan TOF-MS data followed by up to 30x 50ms full scan product ion data in an Information Dependant Acquisition, IDA, mode. Full scan TOFMS data was acquired over the mass range 350-2000 Da and for product ion MS/MS 100-1600 Da. Ions observed in the TOF-MS scan exceeding a threshold of 100 counts and a charge state of +2 to +5 were set to trigger the acquisition of product ion, MS/MS spectra of the resultant 30 most intense ions. The data was acquired and processed using Analyst TF 1.7 software (ABSCIEX, Canada). Protein identification was carried out using Protein Pilot 5.0 for database searching.

#### Mass Photometry

Microscope coverslips were washed and inserted into a Refeyn mass photometry instrument (Refyn Ltd., UK) in the Centre for Microscopy and Microanalysis (CMM). All protein complexes were in a buffer containing 50 mM Tris pH 7.4 and 500 mM NaCl, and all buffers were filtered through a 0.22 μM filter. Calibration was preformed using a mass calibrant purchased from Sigma-Aldrich that contained bovine serum albumin, alcohol dehydrogenase and β-amylase. 6000 frames were collected for each protein and analysed using the Refeyn provided software. Briefly, movies record light scattering events as proteins interaction with the coverslips and the amount of light scattered is quantified and a histogram. Gaussian distributions were then fitted to each peak to determine the molecular weights.

#### Isothermal titration calorimetry (ITC)

The affinities of VPS29 interaction with the synthetic VPS35L peptides and DENND10 with the CCDC22 (325–485) and CCDC93 (310–488) complex and associated mutants was determined using a Microcal PEAQ instrument (Malvern, UK). Experiments we performed in 100 mM Tris (pH 7.4) and 300 mM NaCl. Native and mutant VPS35L peptides at 600 μM were titrated into 20 μM of VPS29, while 50 μM of DENND10 was titrated into 10 μM of wild type and mutant CCDC22 and CCDC93 complexes. In both cases in 13 x 3.22 μL aliquots were used at a temperature of 25°C. The dissociation constants (*K_d_*), enthalpy of binding (Δ*H*) and stoichiometries (N) were obtained after fitting the integrated and normalised data to a single site binding model. The apparent binding free energy (Δ*G*) and entropy (Δ*S*) were calculated from the relationships Δ*G* = RTln(*K*_d_) and Δ*G* = ΔH - TΔ*S*. All experiments were performed at least in triplicate to check for reproducibility of the data.

#### SPARSE matrix crystal screening

COMMD subcomplex C was purified and concentrated to ∼8 mg/ml for crystallization screening. Three commercially available SPARSE matrix hanging-drop crystal screens (LMB, PEGRX, JCSG+) were setup using a Mosquito liquid handling robot (TTP LabTech) at 20°C. Numerous crystal conditions were obtained for COMMD subcomplex C and an initial optimisation screen was performed to determine the best crystallisation conditions. The largest crystals were obtained when the protein solution was supplemented with 2 μM crown ether and 10% glycerol and grown in 22% ethanol and 5 mM EDTA. This condition was optimised in a 24 well vapour diffusion plate on glass cover slips by hanging drop, mixing 5 μl protein with 1 μl reservoir solution. Crystals were relatively small with a diamond-shaped morphology (maximum dimensions ∼50 μm). For data collection crystals were cryo-protected in reservoir solution containing 25% glycerol for 10 s prior to flash-cooling in liquid nitrogen. Likewise, VPS29 was concentrated to 16 mg/ml and incubated with 10 mM of the VPS35L peptide (^16^EFASCRLEAVPLEFGDYHPLKPI^38^; Genscript, USA). Using the same SPARSE matrix screens, and crystals were obtained in JCSG+ H3 (0.1 M Bis-tris pH 5.5 and 25% (w/v) PEG3350). 24 well trays using the same solution resulted in large rod-shaped crystals. For data collection crystals were cryo-protected in reservoir solution containing 20% glycerol for 10 s prior to flash-cooling in liquid nitrogen.

#### Crystallographic data collection and structure determination

Data was collected at the Australian synchrotron on the MX2 beamline. The data was integrated with XDS (140) and scaled with AIMLESS (122) in the CCP4 suite (141). Initially the structure was solved by molecular replacement using PHASER (142) within the PHENIX suite (138). For subcomplex C the templates used for molecular replacement searches were the available COMM domain dimer of COMMD9 (PDB: 6BP6) (45), the HN domain of COMMD9 (PDB: 4OE9) (45) and the HN domain of COMMD10 (*unpublished*). From analysis of the unit cell volume and Matthews Coefficient using XTRIAGE (138), it was estimated that a single copy of the COMMD5-7-9-10 tetramer was present in the asymmetric unit. PHASER was able to successfully place four copies of the COMMD9 COMM domain, one copy of the COMMD9 HN domain and one copy of the COMMD10 HN domain. The resulting model and electron density was sufficient to unambiguously determine the identities of each individual COMM domain. These COMM domains were rebuilt in COOT (139) allowing clear definition of the core heterotetramer of the COMMD5-7-9-10 COMM domains in the structure. Electron density for the two HN domains positioned by PHASER was relatively poor but they could be identified confidently as belonging to COMMD9 and COMMD10 based on their connectivity to the core COMM domains of these two subunits. Further refinement and rebuilding using a combination of PHENIX, COOT and the ChimeraX plugin, ISOLDE resulted in a final model with excellent refinement statistics and stereochemistry based on Molprobity scores. Despite the quality of the final structure and resulting maps, no electron density was observed for the N-terminal HN domains of either COMMD5 or COMMD7. This is likely due to flexibility in the orientation of these domains. VPS29 bound to the VPS35L peptide was solved using the same method as above, however we used the AlphaFold2 predicted structure of the complex as the input template for molecular replacement.

#### AlphaFold2 Modelling of the Retriever, the CCC complex and assembly of the Commander complex

All protein models were generated using AlphaFold2 Multimer (57, 58) implemented in the Colabfold interface available on the Google Colab platform (59). A final model was compiled by combining 3 models each of ∼2000 aa (the current limit of this platform). The models were as follows: COMMD1-10 + CCDC22(1–223), COMMD1-10 + CCDC93(1–300); CCDC22 + CCDC93 + DENND10; VPS29 + VPS26C + VPS35L; VPS35L + CCDC22(1-115; 368-627) + CCDC93(378–630) (see **Fig. S13**). Typically, three independent models were generated for each complex and the quality of the predicted complexes was assessed by examining multiple outputs including the iPTM scores (confidence scores for interfacial residues), predicted alignment error (PAE) plots, and finally a visual inspection of how well the resulting structures aligned with each other in PyMol. Notably, the various complexes invariably displayed highly consistent interfaces across multiple predictions. To generate the final assembled Commander complex, we merged the various predicted structures into a single PDB file, and then models for which we had experimental structures (COMMD1-10+CCDC22+CCDC93, VPS29+VPS35L peptide, and VPS29+VPS26C+VPS35L) were substituted where appropriate. This complete model was then refined using Phenix to fix various stereochemistry parameters including bond length and Ramachandra outliers to produce a final model. Similarly, we generated analogous complexes using AlphaFold2 implemented in ColabFold to model structures of the *Danio rerio* (zebrafish) and *Salpingoeca rosetta* (single cell choanoflagellate). These were entirely consistent with the predicted human complex.

#### Cell lines

Human cell lines (HeLa, HEK293T, and RPE-1) were cultured in humidified incubators at 37°C, 5% CO_2_ in DMEM (Sigma, Catalogue number D5796) supplemented with 10% (v/v) foetal bovine serum (Sigma, catalogue number F7524) and penicillin/streptomycin (Gibco).

#### Generation of HeLa Retriever KO cell lines

VPS35 knock-out HeLa cells were previously generated (120). To generate VPS29 or VPS35L KO HeLa cells, gRNAs targeting genes of interest were designed using the Broad Institute GPP sgRNA Designer (Supplementary Table 6) and cloned into pSpCas9(BB)-2A-GFP (PX458). HeLa cells were transfected with 2 µg pX458 using FuGene, according to manufacturer’s instructions. Cells were incubated for 24 hours before cells were sorted for GFP expression by FACS. Single cells were deposited into 96 well plates containing Iscov’s modified Dulbecco’s medium (Sigma-Aldrich) supplemented with 10% (vol/vol) FBS (Sigma-Aldrich) and Penicillin/Streptomycin. Single cell clones were expanded and screened for gene KO by lysis and Western blotting.

#### Generation of eHAP COMMD KO cell lines

Human eHap cells were obtained from Horizon Discovery. Cells were cultured in Iscove′s Modified Dulbecco′s Medium (IMDM) supplemented with 10% (v/v) fetal calf serum (FCS; CellSera), and penicillin/streptomycin (Gibco) at 37°C under an atmosphere of 5% CO_2_. Constructs for CRISPR-Cas9 genome editing were designed using the CHOPCHOP website (143) and oligonucleotides encoding gRNA sequences cloned into the pSpCas9(BB)-2A-GFP (PX458) plasmid (a gift from F. Zhang (144); Addgene, plasmid 48138) as previously described (145). The gRNA sequences and targeting loci are described in Supplementary Table 7. Constructs were transfected using Lipofectamine 3000 (ThermoFisher Scientific) according to manufacturer’s instructions, and single GFP positive cells sorted into 96 well plates. Clonal populations were expanded and screen by a combination of SDS-PAGE and immunoblotting and Sanger sequencing of genomic PCR products cloned into pGEM4Z (146). Genomic mutations detected by Sanger sequencing are described in Supplementary Table 7. For generation of FLAG-tagged cell lines, inserts containing cDNA sequences were commercially synthesized (IDT technologies) to contain a C-terminal FLAG tag and compatible overhangs for Gibson assembly. Inserts were combined with pBABE-puro plasmid (Addgene, 1764) cut with BamHI-HF and HindIII-HF restriction enzymes (NEB) and Gibson assembled using the NEBuilder HiFi DNA Assembly System (NEB) as per manufacturer’s instructions. Retroviral particles were made in HEK293T cells using pUMVC3 and pCMV-VSV-G (Addgene, 8449 and 8454) packaging plasmids as previously described (*50*). Viral supernatant was collected at 48 h post-transfection, filtered with 0.45 μm PVDF membrane (Milipore) and combined with 8 µg ml^−1^ polybrene for transduction. Infected cells were selected using 2 μg ml^−1^ puromycin, and transduction verified by SDS-PAGE and immunoblotting.

#### Molecular cloning and site-directed mutagenesis

To generate VPS35L-GFP, VPS35L was subcloned into the EGFP-N1 or lentiviral pLVX vector. VPS35L was amplified using Q5 High-Fidelity 2X Master Mix (NEB, M0492) following the manufacturer’s protocol. Following PCR, bands were resolved on agarose gel and purified with GFX PCR DNA and Gel Band purification kit (GE Healthcare, 28-9034-70). Amplified gene or 1 µg of plasmid backbone were then digested using appropriate restriction enzymes (1.5 µl), and the plasmid backbone was additionally treated with 1.5 µl of quick-CIP (NEB, M0525) to prevent self-ligation. Digestion reaction was carried out in 1x CutSmart buffer and nuclease-free water in a final volume of 40 µl at 37°C for 1 hour. Digestion products were purified as previous and 50 µg of backbone and 6-times excess of insert were then used for ligation using T4 DNA ligase (Invitrogen, 15224017). Primers for site-directed mutagenesis were designed using Agilent QuikChange® Primer design tool. PCR reactions were carried out using QuikChange II Site-Directed Mutagenesis Kit (Agilent, 200523-5) following the manufacturer’s protocol. After PCR, non-mutated template vector was removed from the PCR mixture through digestion for 1 hour at 37°C by the enzyme Dpn1. Following digestion by Dpn1, 4 μl of the mixture was transformed into XL10 Gold (Agilent, 200315) chemically competent cells and plated on suitable antibiotic-containing agar plates. Sequencing of purified plasmid DNA established whether the desired mutation had been introduced.

#### Gibson assembly

NEBuilder Assembly Tool was used to design primers for Gibson Assembly reactions. The fragments were amplified using overlapping primers. 0.02 pmol of pLVX_Puro digested with EcoRI and BamHI and −0.04-0.06 pmol PCR-amplified fragments were mixed with Gibson Assembly 2x Master Mix (NEB, E2611) according to manufacturer’s instructions and incubated for 1h at 50°C. 2 µl of the reaction was transformed into NEB 5-alpha Competent *E. coli* (NEB, C2987H) cells.

#### Transfection and transduction of cell lines

PEI (polyethylenimine) was used to transfect HEK 293 cells with constructs for GFP/mCherry traps or to produce lentivirus. An aqueous 10 µg/ml stock of linear 25 kDa PEI (Polysciences, catalogue number 23966-2) was used for transfections. For 10 cm or 15 cm, 2.5 ml or 5 ml of Opti-MEM^β^ was added to 2 separate sterile tubes respectively. In the first Opti-MEM^β^ containing tube, 10 µg or 15 µg DNA was added for 10 cm or 15 cm dishes respectively. To the second tube, 3:1 PEI:DNA ratio was added and the contents of the tube mixed by vortexing. The Opti-MEM^β^ /PEI mixture was then filter sterilised by filtering through a 0.2µm filter. The sterilised PEI/Opti-MEM^β^ was then added to the Opti-MEM^β^ /DNA mixture and the tube was mixed by vortexing. The mixture was left to incubate at room temperature for 20 min. Following incubation, HEK 293 cells were washed in PBS, then PBS was removed and the transfection mixtures were carefully added to the cell dishes. HEK 293 cells were incubated with the transfection mixture, under normal growth conditions, for 4 h. The transfection media was removed at the end of the incubation period and replaced with normal growth media. Cells were further incubated for another 24/48 hours prior to experimental use. To generate lentivirus, a 15 cm dish of HEK 293 cells were transfected with 15 µg of PAX2, 5 µg pMD2.G and 20 µg of lentiviral expression vector using PEI as described above. After the 48 h incubation, the growth media containing the lentivirus was harvested and filtered through a 0.45 µm filter. Cells to be transduced were seeded at 50,000 cells per well of a 6-well plate and left to settle prior to addition of lentivirus.

#### GFP/mCherry nanotraps

Dishes containing cells expressing GFP/mCherry or GFP/mCherry tagged proteins (either transiently or stably) were placed on ice. The cell media was removed and the cells were washed three times with ice cold PBS (Sigma). Cells were lysed with lysis buffer (20 mM Hepes pH 7.2, 50 mM potassium acetate, 1 mM EDTA, 200 mM D-sorbitol, 0.1% Triton X-100, 1x protease cocktail inhibitor, pH7.5 or 50mM Tris pH7.5 with 0.5% NP-40 in PBS with protease inhibitors). 500 µl or 1 ml of lysis buffer was used per 10 cm or 15 cm dish respectively. Lysis was aided through the use of a cell scraper. The lysates were then cleared by centrifugation at 13,200 rpm for ten minutes at 4°C. 15 µl of GFP-trap beads (Chromotek, catalogue number gta-20) or mCherry-trap beads (Chromotek, catalogue number rta-20) were pre-equilibrated in lysis buffer, through three rounds of washing in lysis buffer, prior to adding cleared cell lysate. 10% of cell lysate was retained for input analysis. Trap beads and lysates were incubated together on a rocker at 4°C for 1 h. Following incubation, Trap beads were pelleted by centrifugation at 2000 rpm, for 30 seconds at 4°C. Supernatant was then removed, and beads were either washed a further three times in 20 mM Hepes pH 7.2, 50 mM potassium acetate, 1 mM EDTA, 200 mM D-sorbitol, 0.1% Triton X-100, 1x protease cocktail inhibitor, pH7.5 or twice in 50 mM Tris pH7.5 with 0.25% NP-40 in PBS with protease inhibitors and once with 50mM Tris pH7.5 in PBS with protease inhibitors through rounds of re-suspension and pelleting. After the final wash, all lysis buffer was removed from the Trap beads. Beads were then either stored at −20°C or processed for SDS-PAGE analysis.

#### Quantitative western blot analysis

BCA assay kit (Thermo Fisher Scientific, USA) was used to determine protein concentration with equal amounts being resolved on 4%-12% NuPAGE precast gels (Invitrogen, USA). Polyvinylidene fluoride membranes (Immobilon-FL; EMD Millipore, USA) were used for transfer with protein detection quantified using the Odyssey infrared scanning system (LI-COR Biosciences, USA) and fluorescently labelled secondary antibodies. We routinely performed western blot quantification where a single blot is simultaneously probed with distinct antibody species targeting proteins of interest followed by visualisation of secondary antibodies conjugated with distinct spectral dyes. All quantified western blots are the mean of at least 3 independent experimental repeats, with statistical analysis performed using Prism 7 (GraphPad Software, USA). All quantitation of western blots is shown in **Fig. S15.**

#### Biotinylation of cell surface proteins

Fresh Sulfo-NHS-SS Biotin (Thermo Fischer Scientific, no. 21217) was dissolved in 4°C PBS (pH 7.4) at 0.2 mg/ml prior to incubating with prewashed (twice with ice-cold PBS) cells placed on ice to reduce the rate of endocytosis and endocytic pathway flux. Cells were incubated for 30 minutes at 4°C, followed by incubation in TBS for 10 minutes to quench the biotinylation reaction. Cells were then lysed in lysis buffer and subjected to Streptavidin bead-based affinity isolation (GE Healthcare, USA).

#### Immunofluorescence staining

Cells were seeded onto sterile 13mm glass coverslips. Once cells were ready to be fixed, growth media was aspirated off and cells were washed three times in PBS prior to fixation in 4% PFA (w/v) (paraformaldehyde, Pierce™ 16% Formaldehyde (w/v), Methanol-free, catalogue number 28906, diluted to 4% (w/v) in PBS). Cells were incubated in 4% PFA for 20 minutes at room temperature. Coverslips were then washed a further 3 times in PBS. For permeabilization, coverslips were incubated in 0.1% (v/v) Triton^β^ X-100 in PBS for 5 minutes at room temperature. Alternatively, if the cells were going to be stained for LAMP1, cells were permeabilised in 0.1% (w/v) saponin in PBS for 5 minutes. After permeabilization, coverslips were then washed a further 3 times in PBS. Coverslips were blocked for 15 minutes in 1% (w/v) BSA in PBS at room temperature. Primary antibodies were diluted in 1% (w/v) BSA in PBS (for Triton^β^ X-100 permeabilised cells) or 1% (w/v) BSA, 0.01% (w/v) saponin in PBS. 60µl of diluted antibody solution was pipetted onto a strip of Parafilm as a dot. Coverslips were inverted and placed onto the dots so that the cells were immersed into the antibody solution. Coverslips were incubated with primary antibody for 1 hour at room temperature. Coverslips were then washed three times in PBS before placing onto 60 µl dots containing Alexafluor-conjugated secondary antibodies and DAPI (if required, 0.5 µg/ml) for 1 hour at room temperature, then washed 3 times in PBS and once in water. Coverslips were mounted onto glass microscope slides in Fluoromount-G (Invitrogen, 004958-02).

#### Confocal Microscopy

Fixed cells were imaged at room temperature using a Leica SP5, Leica SP5-II or Leica SP8 multi-laser confocal microscope. A 63x NA 1.4 UV oil-immersion lens was used to take all images. Leica LCS or LAS X software was used for the acquisition of images. Colocalisation analysis was performed in Volocity 6.3.1 software (PerkinElmer) with automatic Costes background thresholding.

#### Affinity enrichment mass-spectrometry of FLAG-tagged COMMD subunits

Cell pellets (triplicate sub-cultures representing each cell line) were harvested by scraping and washed in PBS (137 mM NaCl, 2.7 mM KCl, 10 mM Na_2_HPO_4_, 1.8 mM KH_2_PO_4,_ pH 7.4). Protein concentration was determined using the Pierce Protein Assay Kit (ThermoFisher Scientific), following which 1 mg of material was solubilized for affinity enrichment mass-spectrometry (AEMS) as previously described (*50*). Briefly, cell pellets were solubilized in 20mM Tris-Cl pH 7.4, 50mM NaCl, 10% (v/v) glycerol, 0.1mM EDTA, 1% (w/v) digitonin and 125 units of benzonase (Merck) and soluble material loaded onto Pierce™ Spin Columns (ThermoFisher Scientific) containing anti-FLAG M2 affinity gel (Sigma) pre-equilibrated with 20mM Tris-Cl pH 7.4, 60mM NaCl, 10% v/v glycerol, 0.5mM EDTA, 0.1% w/v digitonin. Following a 2 h incubation at 4°C, columns were washed with the same buffer and enriched protein complexes eluted with the addition of 100 μg ml^−1^ FLAG peptide (Sigma). Eluates were acetone precipitated and precipitates resuspended in 8 M urea in 50 mM ammonium bicarbonate (ABC). Proteins were reduced and alkylated by incubation at 37°C for 30 mins with 10 mM tris(2-carboxyethyl)phosphine hydrochloride (TCEP; Bondbreaker, ThermoFisher Scientific) and 50 mM chloroacetamide (Sigma Aldrich), following which samples were diluted to 2 M urea using 50 mM ABC prior to digestion with 1 μg of trypsin (ThermoFisher Scientific) at 37°C overnight. Peptides were acidified to 1% Trifluoroacetic acid (TFA) and desalted using stagetips containing 2x 14G plugs of 3M™ Empore™ SDB-XC Extraction Disks (Sigma) as described (145). Peptides dried using CentriVap concentrator (Labconco) and samples reconstituted in 0.1% TFA, 2% CAN for analysis by mass-spectrometry.

For COMMD1^FLAG^, COMMD6^FLAG^, and COMMD9^FLAG^ and parental eHap1 cell lines, eluates prepared as above were analysed on an LTQ Orbitrap Elite (Thermo Scientific) in conjunction with an Ultimate 3000 RSLC nanoHPLC (Dionex Ultimate 3000) using the liquid chromatography (LC) and mass-spectrometry instrument parameters previously described (147). The basic LC setup consisted of a trap column (Dionex-C18 trap column 75 μm × 2 cm, 3 μm, particle size, 100 Å pore size; ThermoFisher Scientific) run at 5 μl/min before switching the pre-column in line with the analytical column (Dionex-C18 analytical column 75 μm × 50 cm, 2 μm particle size, 100 Å pore size; ThermoFisher Scientific). The separation of peptides was performed at 300 nL/min using a 95 min non-linear ACN gradient of buffer A [0.1% formic acid, 2% ACN, 5% DMSO] and buffer B [0.1% formic acid in ACN, 5% DMSO]. Mass-spectrometry data were collected in Data Dependent Acquisition (DDA) mode using m/z 300–1650 as MS scan range, rCID for MS/MS of the 20 most intense ions. Lockmass of 401.92272 from DMSO was used. Other instrument parameters were: MS scan at 100,000 resolution, maximum injection time 150 ms, AGC target 1_E_6, CID at 30% energy for a maximum injection time of 150 ms with AGC target of 5000. Dynamic exclusion with of 30 seconds was applied for repeated precursors. For COMMD2^FLAG^, COMMD4^FLAG^, COMMD5^FLAG^, COMMD7^FLAG^, COMMD8^FLAG^, COMMD10^FLAG^ and parental eHap1 cell lines, eluates were analysed on an Orbitrap Exploris 480 Thermo Scientific) in conjunction with an Ultimate 3000 RSLC nanoHPLC (Dionex Ultimate 3000) using liquid chromatography and mass-spectrometry instrument parameters previously described (147). The LC setup was identical to that described above, except that for COMMD2^FLAG^, COMMD4^FLAG^, COMMD5^FLAG^, COMMD7^FLAG^, COMMD8^FLAG^ and parental control, the non-linear ACN gradient used for the separation of peptides was 65 mins in length. Mass spectrometry was conducted in data-dependent acquisition mode, whereby full MS1 spectra were acquired in a positive mode at 120000 resolution using a scan range of 300 – 1600 m/z. The ‘top speed’ acquisition mode with 3 s cycle time on the most intense precursor ion was used, whereby ions with charge states of 2 to 6 were selected. MS/MS analyses were performed by 1.2 m/z isolation with the quadrupole, fragmented by HCD with collision energy of 30%. MS2 resolution was at 15000. AGC target was set to *standard* with auto maximum injection mode. Dynamic exclusion was activated for 20 s.

Affinity enrichment mass-spectrometry data were analyzed using the MaxQuant (148) and Perseus (149) platforms as previously described for similar data in (150). In brief, raw mass-spectrometry data from each batch of AEMS experiments were separately analysed in MaxQuant with the data combined during workup in Perseus. Default MaxQuant search parameters were used with “Label free quantitation” set to “LFQ” and “Match between runs” enabled. Trypsin/P cleavage specificity (cleaves after lysine or arginine, even when proline is present) was used with a maximum of 2 missed cleavages. Oxidation of methionine and N-terminal acetylation were specified as variable modifications. Carbamidomethylation of cysteines was set as a fixed modification. A search tolerance of 4.5 ppm was used for MS1 and 20 ppm for MS2 matching. False discovery rates (FDR) were determined through the target-decoy approach set to 1% for both peptides and proteins. The MaxQuant ProteinGroups.txt output tables were imported into Perseus and LFQ intensities were log2 transformed. Values listed as being “Only identified by site,” “Reverse,” or “Contaminants” were removed from the data set. Experimental groups were assigned to each set of triplicates and the number of valid values for each row group calculated. For each experiment (containing a control and an enrichment group), single replicates with significant variation as evident through a principal component analysis (PCA) were removed, along with rows having less than 2 valid values in the enrichment group. Missing values in the relevant control group were imputed to values consistent with the limit of detection. A two-sided, two-sample Student’s t-test was performed between control and each enrichment group, with the resulting data plotted on volcano plot. The threshold of significant enrichment was set to 2-fold (log2 fold change = 1) based on the distribution of unenriched proteins quantified.

#### TMT labelling and high pH reversed-phase chromatography

The samples were reduced (10 mM TCEP, 55°C for 1 h), alkylated (18.75 mM iodoacetamide, room temperature for 30 min) and then digested from the beads with trypsin (2.5 µg trypsin; 37°C, overnight). The resulting peptides were then labeled with TMT seven-plex reagents according to the manufacturer’s protocol (Thermo Fisher Scientific, Loughborough, LE11 5RG, UK) and the labelled samples pooled and desalted using a SepPak cartridge according to the manufacturer’s instructions (Waters, Milford, Massachusetts, USA). Eluate from the SepPak cartridge was evaporated to dryness and resuspended in buffer A (20 mM ammonium hydroxide, pH 10) prior to fractionation by high pH reversed-phase chromatography using an Ultimate 3000 liquid chromatography system (Thermo Scientific). In brief, the sample was loaded onto an XBridge BEH C18 Column (130 Å, 3.5 µm, 2.1 mm X 150 mm, Waters, UK) in buffer A and peptides eluted with an increasing gradient of buffer B (20 mM Ammonium Hydroxide in acetonitrile, pH 10) from 0-95% over 60 minutes. The resulting fractions (5 in total) were evaporated to dryness and resuspended in 1% formic acid prior to analysis by nano-LC MSMS using an Orbitrap Fusion Tribrid mass spectrometer (Thermo Scientific).

#### Nano-LC Mass Spectrometry

High pH RP fractions were further fractionated using an Ultimate 3000 nano-LC system in line with an Orbitrap Fusion Tribrid mass spectrometer (Thermo Scientific). In brief, peptides in 1% (vol/vol) formic acid were injected onto an Acclaim PepMap C18 nano-trap column (Thermo Scientific). After washing with 0.5% (vol/vol) acetonitrile 0.1% (vol/vol) formic acid peptides were resolved on a 250 mm × 75 μm Acclaim PepMap C18 reverse phase analytical column (Thermo Scientific) over a 150 min organic gradient, using 7 gradient segments (1-6% solvent B over 1 min, 6-15% B over 58 min, 15-32% B over 58 min, 32-40% B over 5 min, 40-90% B over 1 min, held at 90% B for 6 min and then reduced to 1% B over 1min.) with a flow rate of 300 nl min^−1^. Solvent A was 0.1% formic acid and Solvent B was aqueous 80% acetonitrile in 0.1% formic acid. Peptides were ionized by nano-electrospray ionization at 2.0 kV using a stainless-steel emitter with an internal diameter of 30 μm (Thermo Scientific) and a capillary temperature of 275°C. All spectra were acquired using an Orbitrap Fusion Tribrid mass spectrometer controlled by Xcalibur 2.1 software (Thermo Scientific) and operated in data-dependent acquisition mode using an SPS-MS3 workflow. FTMS1 spectra were collected at a resolution of 120,000, with an automatic gain control (AGC) target of 200,000 and a max injection time of 50 ms. Precursors were filtered with an intensity threshold of 5000, according to charge state (to include charge states 2-7) and with monoisotopic peak determination set to peptide. Previously interrogated precursors were excluded using a dynamic window (60s +/-10 ppm). The MS2 precursors were isolated with a quadrupole isolation window of 1.2m/z. ITMS2 spectra were collected with an AGC target of 10,000, max injection time of 70ms and CID collision energy of 35%. For FTMS3 analysis, the Orbitrap was operated at 50,000 resolution with an AGC target of 50,000 and a max injection time of 105 ms. Precursors were fragmented by high energy collision dissociation (HCD) at a normalised collision energy of 60% to ensure maximal TMT reporter ion yield. Synchronous Precursor Selection (SPS) was enabled to include up to 10 MS2 fragment ions in the FTMS3 scan.

#### Proteomic Data Analysis

The raw data files were processed and quantified using Proteome Discoverer software v2.1 (Thermo Scientific) and searched against the UniProt Human database (downloaded January 2021; 169297 sequences) using the SEQUEST HT algorithm. Peptide precursor mass tolerance was set at 10 ppm, and MS/MS tolerance was set at 0.6 Da. Search criteria included oxidation of methionine (+15.995Da), acetylation of the protein N-terminus (+42.011Da) and Methionine loss plus acetylation of the protein N-terminus (−89.03Da) as variable modifications and carbamidomethylation of cysteine (+57.021Da) and the addition of the TMT mass tag (+229.163Da) to peptide N-termini and lysine as fixed modifications. Searches were performed with full tryptic digestion and a maximum of 2 missed cleavages were allowed. The reverse database search option was enabled and all data was filtered to satisfy false discovery rate (FDR) of 5%.

#### Phylogenetic analyses

Representative sequences of CCDC22, CCDC93, COMMD1, COMMD2, COMMD3, COMMD4, COMMD5, COMMD6, COMMD7, COMMD8, COMMD9 and COMMD10 were used to construct HMM profilers (HMMER 3.3.2) which were then searched against 30 proteomes from a representative selection of organisms (from RefSeq (151) and Genbank (152)) with an E-value threshold of 1×10^-5^. Duplicate COMMD sequences were removed, and representative query sequences were added (for identification of each different COMMD protein) before sequences were aligned using MAFFT L-INS-i (v7.505) (132), with separate alignments for CCD22, CCD93 and one alignment for all 10 COMMD proteins. Maximum likelihood trees were then inferred using IQTree (details below) and manually inspected, and outgroups to the COMMD10, CCD22 and CCD93 clades were removed from the unaligned sequence sets before alignment and subsequent tree inference. All maximum likelihood trees were inferred under the best-fitting model according to the Bayesian Information Criterion implemented in ModelFinder (part of IQTree2.1.3 (133)), including complex models allowing for across-site compositional heterogeneity (-m MFP -madd LG+C60+F+G,LG+C50+F+G,LG+C40+F+G,LG+C30+F+G,LG+C20+F+G,LG+C10+F+G,LG+F+G, LG+R+F --score-diff ALL). Each tree was inferred with 10,000 ultrafast bootstrap replicates (*61*). The best-fitting models were Q.yeast+R5 for the COMMD1-10 tree, LG+C20+F+G for CCD22, and Q.insect+F+I+G4 for CCD93.

### QUANTIFICATION AND STATISTICAL ANALYSIS

For data analysis of FLAG-tagged COMMD subunit mass spectrometry experiments, raw files were analysed using the MaxQuant platform (153), version 1.6.10.43 against canonical, reviewed and isoform variants of human protein sequences in FASTA format (Uniprot, January 2019). The default settings: “LFQ” and “Match between runs” were enabled. N-terminal acetylation and methionine oxidation were set as variable modifications while cysteine carbamidomethylation was specified as a fixed modification. Computation of protein enrichment was performed in Perseus (version 1.6.10.43) (149). Peptides labelled by MaxQuant as ‘only identified by site’, ‘reverse’ or ‘potential contaminant’ were removed and only those proteins quantified based on >1 unique peptide were considered for further analysis. LFQ intensities were log2 transformed and rows having less than 3 valid values in the enrichment group were removed and the missing values in the control group were imputed to values consistent with the limit of detection. The mean log_2_ LFQ intensities for proteins detected in each experimental group, along with p-values, were calculated using a two-sided two-tailed t-test. Significance was determined by permutation-based FDR statistics (149) where the s0 factor was iteratively modified to exclude all identifications enriched in the control experiment, yielding an s0 of 1 at 1% FDR.

For quantitation of Western blots protein detection was performed using the Odyssey infrared scanning system (LI-COR Biosciences, USA) and fluorescently labelled secondary antibodies. We routinely performed western blot quantification where a single blot is simultaneously probed with distinct antibody species targeting proteins of interest followed by visualisation of secondary antibodies conjugated with distinct spectral dyes. All quantified western blots are the mean of at least 3 independent experimental repeats, with statistical analysis performed using Prism 7 (GraphPad Software, USA). Colocalisation analysis of fluorescently labelled proteins was performed in Volocity 6.3.1 software (PerkinElmer) with automatic Costes background thresholding.

## SUPPLEMENTARY INFORMATION

### SUPPLEMENTARY MOVIES

**Supplementary Movie S1. Structure of Retriever determined by cryoEM and AlphaFold2 modelling.**

**Supplementary Movie S2. Structure of the CCC complex determined by cryoEM.**

**Supplementary Movie S3. Overall model of the Commander complex derived from cryoEM, X-ray crystallography and AlphaFold2 modelling.**

**Supplementary Figure S1.**
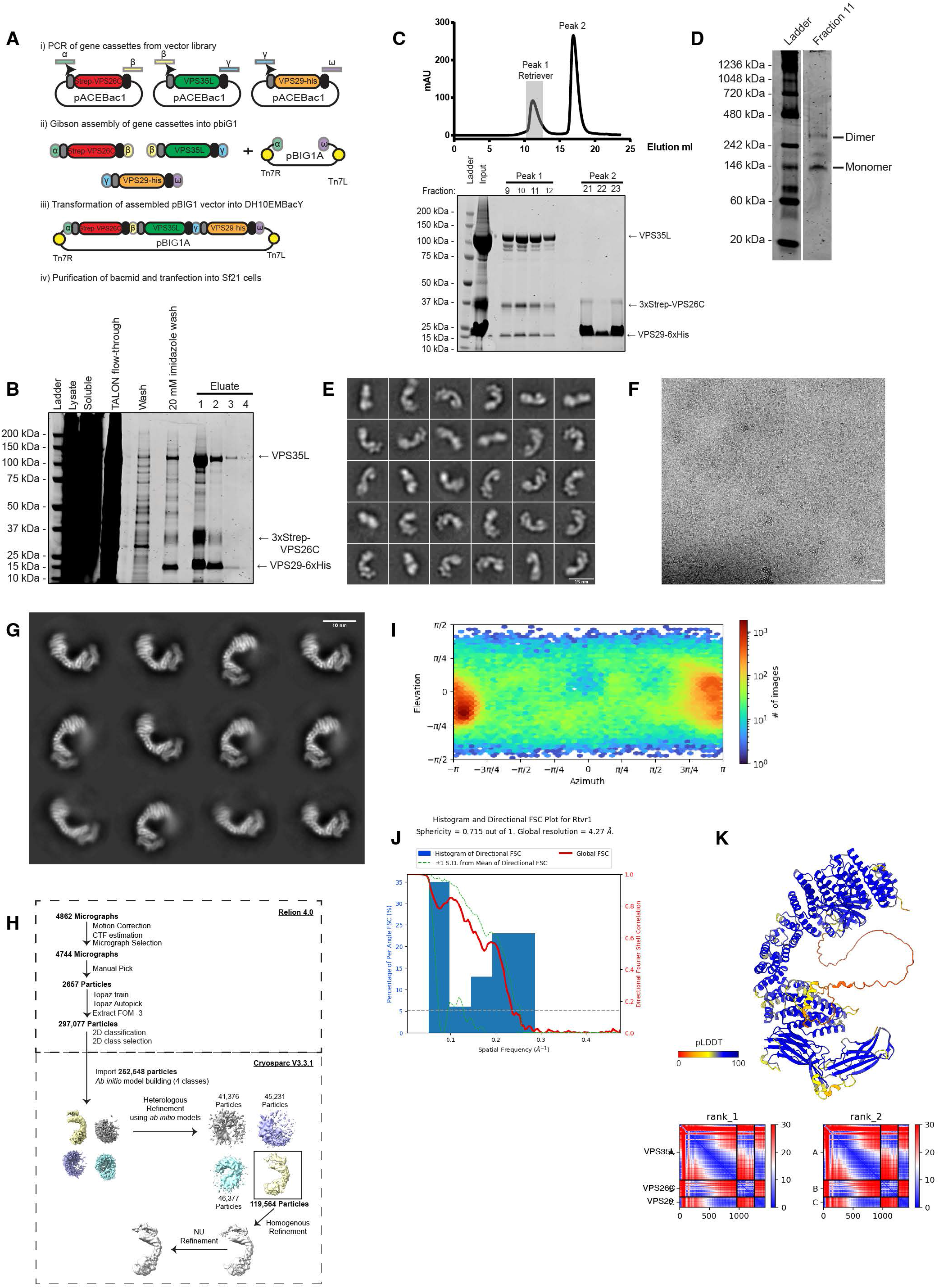
Expression, purification, cryoEM analysis and Alphafold modelling of the human Retriever complex. (A) Schematic for the biGBac cloning approach used to co-express Retriever in insect cells. **(B)** Representative Coomassie stained SDS-PAGE gel of the isolation of recombinant Retriever from Sf21 insect through affinity purification of VPS29-6xhis. Eluate fractions were combined and subjected to gel filtration. **(C)** Gel filtration profile of his-purified Retriever. Retriever was gel filtrated using a Superdex200 column. Fractions corresponding to A280 peaks were analysed by running SDS-PAGE gel and stained with Coomassie. Full Retriever complex corresponds to Peak 1. **(D)** Native PAGE of purified Retriever. **(E)** Negative stain electron microscopy analysis of Retriever revealed the elongated ‘footprint’-like morphology. Representative 2D classification classes of negatively stained Retriever. Scale bar represents 15 nm. **(F)** Representative motion-corrected micrograph. **(G)** Representative single particle cryoEM 2D class averages of Retriever. Note that only a ‘front’ view of the complex is visible in 2D classes. Scale bar represents 10 nm. **(H)** Data processing used to obtain a low resolution cryo-EM reconstruction of Retriever. **(I)** Angular projections of Retriever particles used for the final 3D reconstruction, indicating preferential orientation of the particle. Heat map calculated in CryoSPARC displaying the number of particles per viewing orientation. **(J)** Directional FSC plots and sphericity values for the Retriever reconstruction. These data were generated using a 3D-FSC server (https://3dfsc.salk.edu/). **(K)** AlphaFold2 prediction of Retriever with associated PAE plots for the top 2 ranked models.

**Supplementary Fig. S2.**
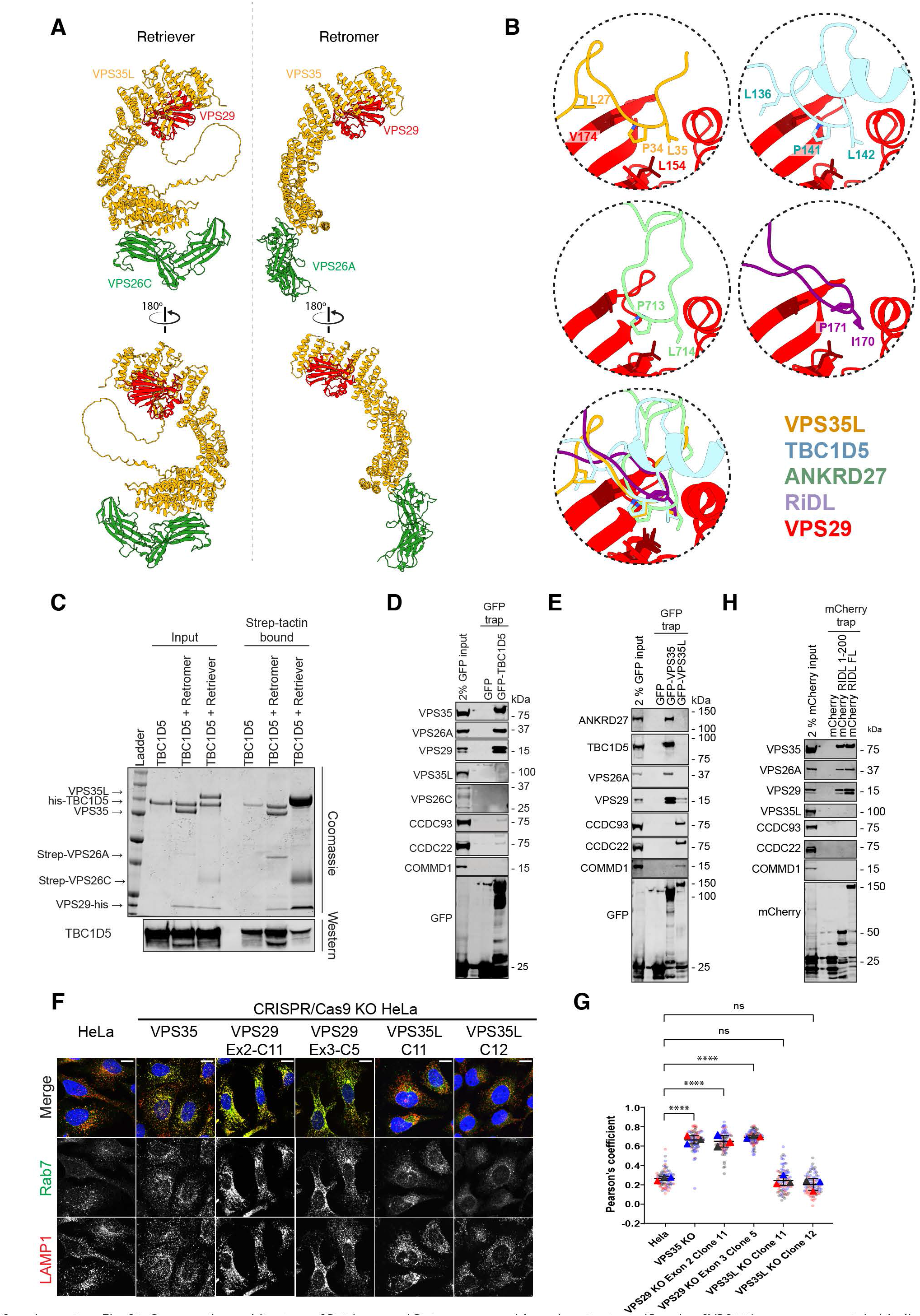
Comparative architecture of Retriever and Retromer assembly and context specific role of VPS29 in accessory protein binding. (A) Comparison between Retriever and Retromer assemblies. **(B).** VPS35L PL motif binding to VPS29 mimics the association of the Retromer accessory proteins, TBC1D5 (5GTU) and ANKRD27 (6TL0), and the *Legionella* effector RidL (5WYH) to VPS29. **(C)** Recombinant Strep-tagged VPS26A-Retromer and Strep-tagged VPS26C-Retriever were individually incubated with recombinant his-tagged TBC1D5 and subjected to Strep-tactin affinity isolation. Coomassie staining and Western analysis of TBC1D5 reveals robust association with Retromer but limited association with Retriever. Gel and blot are representative of two independent experiments. **(D, E, H)** HEK293T cells were transfected with GFP and **(D)** GFP-tagged TBC1D5, **(E)** GFP-VPS35 and GFP-VPS35L, and **(H)** mCherry, mCherry tagged-amino terminal region of RidL (1–200) or full length (FL) RidL and subjected to GFP-nanotrap or mCherry nanotrap. Blots are representative of three independent experiments in each case. **(F)** VPS35L KO cells do not have elevated lysosomal Rab7 levels. HeLa WT or HeLa KO cells were grown on glass coverslips and fixed with 4% PFA in PBS, stained with the indicated antibodies and DAPI and imaged by confocal microscopy. Scale bars represent 10 µm. Representative images from 3 independent experiments. **(G)** Quantification of Pearson’s coefficients between Rab7 and LAMP1 from (F). For each condition, 30 cells were quantified per 3 independent fixation and staining experiments (90 cells total). Pearson’s coefficients for individual cells are represented by transparent circles, coloured according to the independent experiment the cell belonged. The mean of the 30 cells per independent experiment are represented by solid triangles, coloured according to the independent experiment. Bars, mean of the 3 independent experimental means; error bars, s.d. of the 3 independent experimental means. Normality of data was checked prior to one-way ANOVA followed by Dunnett test for multiple comparisons. **** = p <0.0001, ns = not significant.

**Supplementary Fig. S3.**
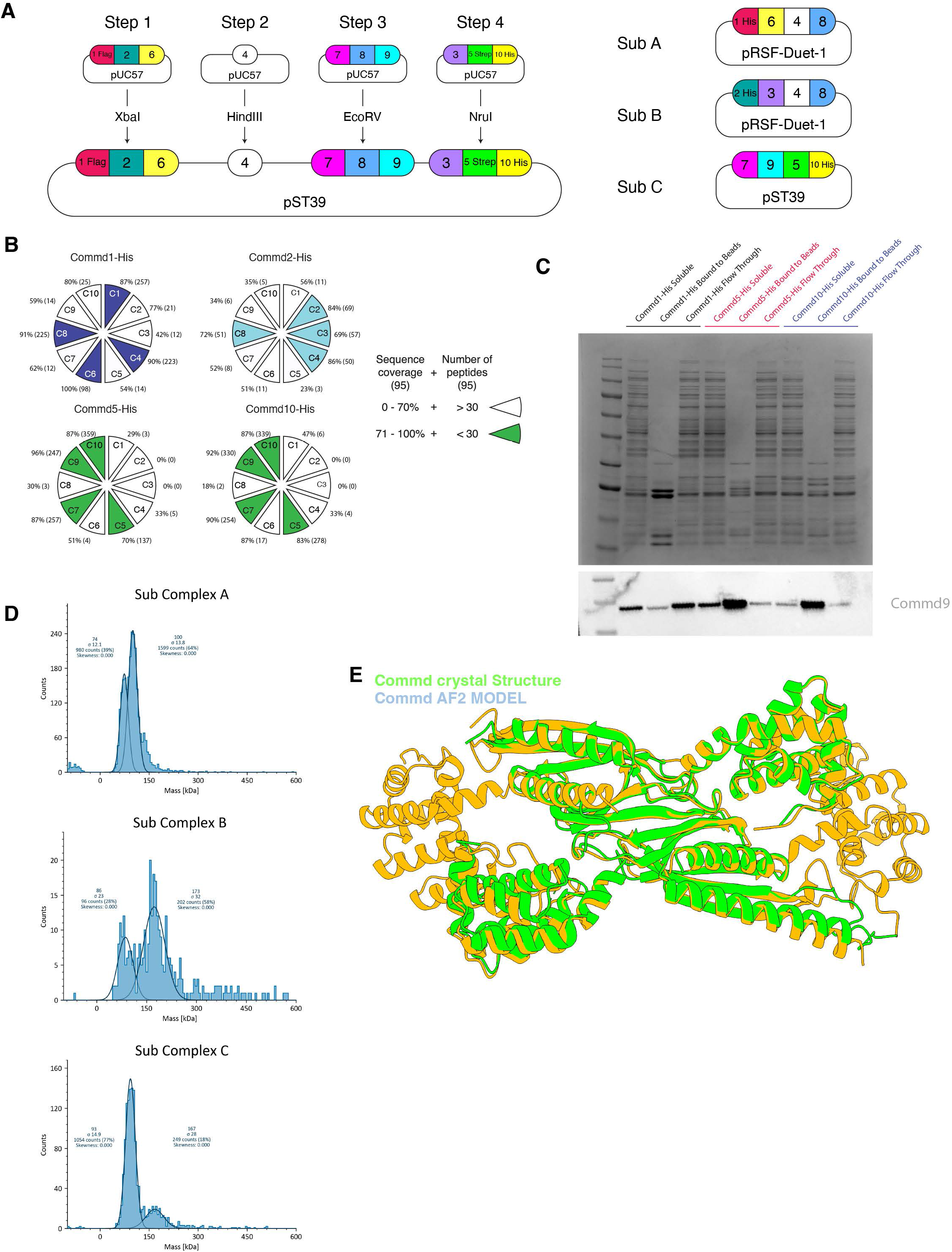
Expression, purification, and characterisation of recombinant COMMD complexes in *E. coli.* (A) Design of co-expression vector for isolation of COMMD complexes from *E. coli*. *Left* gene cassettes were ordered from Gene Universal in pUC57 vectors and sequentially cloned in a pST39 vector for simultaneous expression. This vector included 3 proteins with affinity tags COMMD1-FLAG, COMMD5-StrepII, and COMMD10-His. From this initial vector, 4 vectors were generated that contained only a single His tag on COMMD1, COMMD2, COMMD5 and COMMD10. *Right* shows pRSF-Duet-1 vectors that express the individual COMMD subcomplexes. **(B)** Purification of protein using different His tag locations lead to the proteomic identification of subcomplex A, subcomplex B, and subcomplex C. **(C)** This result was also confirmed by western blot with only COMMD5 and COMMD10 His able to enrich for COMMD9. (**D**) Mass photometry confirmed that the masses of subcomplexes generally agreed with the expected values (A, 74 kDa (74); B, 86 kDa (88); C, 93 kDa (92)). (**E**) Comparison of the AlphaFold 2 (AF2) prediction shows good agreement with the 3.3 Å crystal structure of subcomplex C.

**Supplementary Fig. S4.**
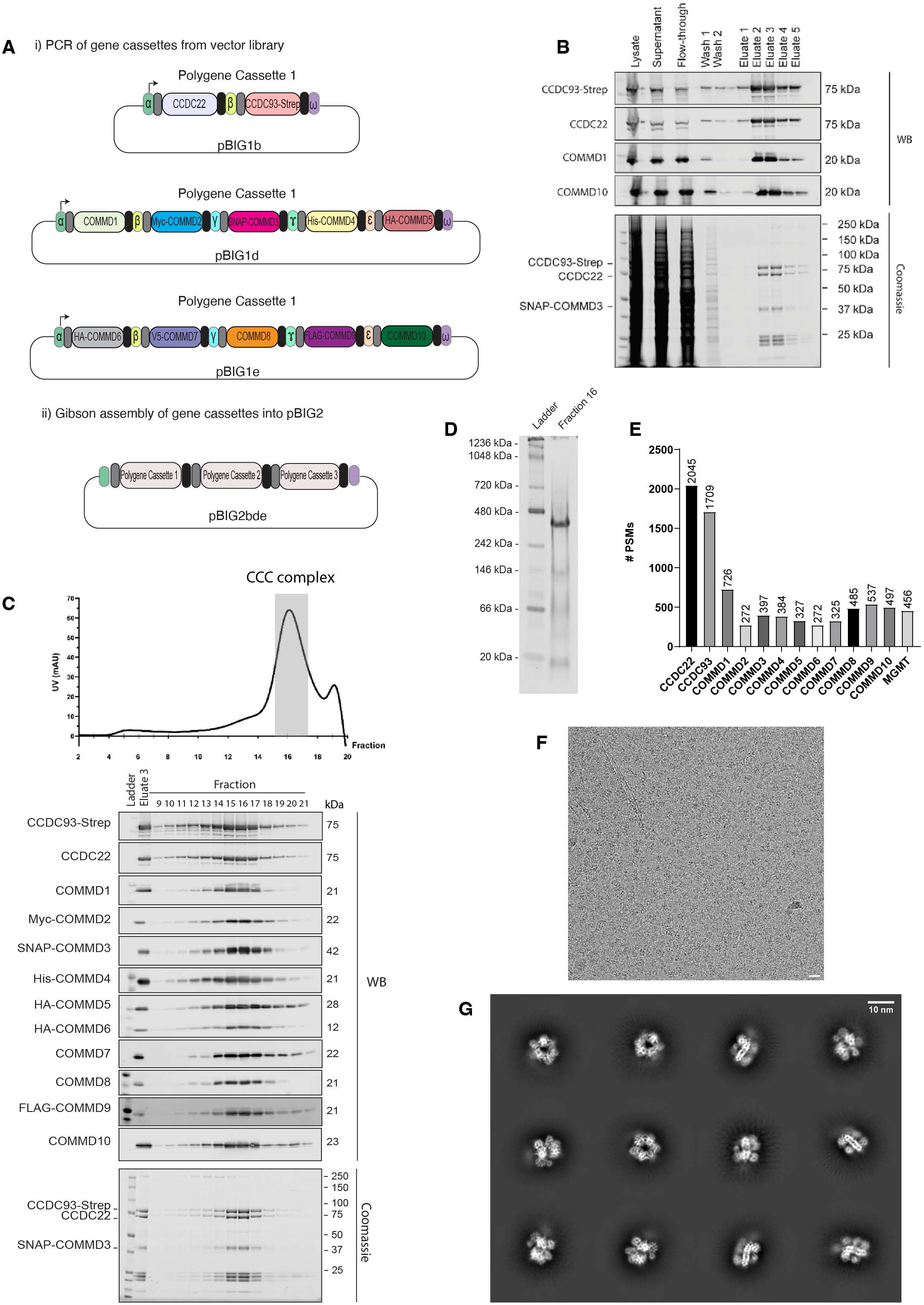
Expression, purification, and characterisation of recombinant CCC complexes in insect cells. (A) Schematic of the engineering of pBIGbde-CCC from Gibson assembly of individual polygene cassettes. **(B)** Coomassie and western analysis of strep-actin affinity purified human CCC complex from baculovirus infected insect cells. **(C)** Gel filtration of the affinity purified CCC complex and western analysis of the purified human CCC complex. **(D)** Native PAGE of purified CCC complex. **(E).** Mass spec analysis of purified CCC complex establishing the presence of all twelve proteins. Data is from one of two independent experiments. **(F)** Representative motion-corrected 10 Å lowpass filtered micrograph of vitrified, cross-linked CCC complex on graphene oxide coated 1.2/1.3 Quantifoil™ grids. Scale bar represents 20 nm. **(G)** Single-particle 2D cryo-EM classes, generated in cryoSPARC, revealing the globular, ring-like structure of the CCC complex. Scale bar represents 10 nm.

**Supplementary Fig. S5.**
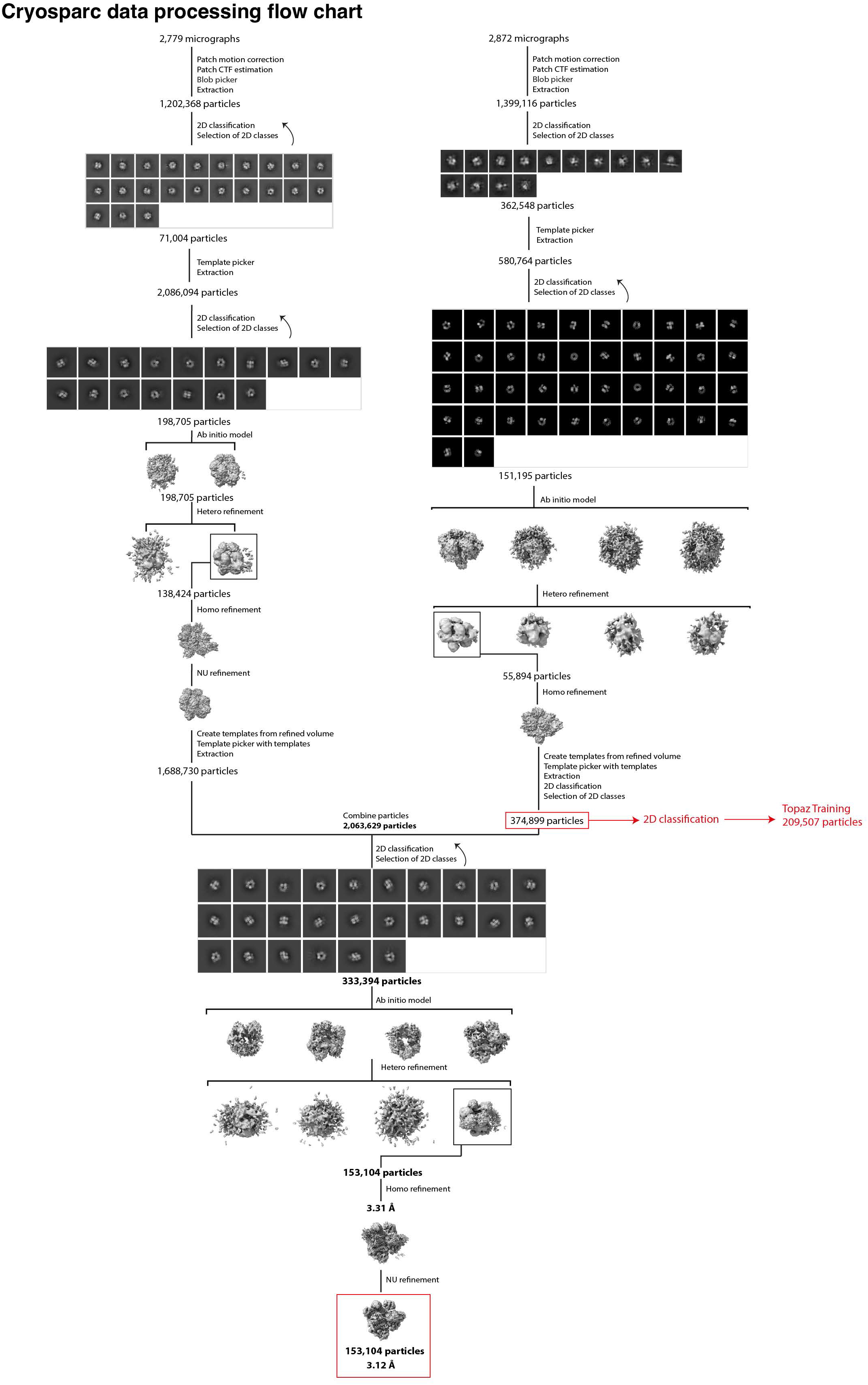
Cryo-EM data analysis of the CCC complex in cryoSPARC. Flow chart of the data processing strategy using CryoSPARC to obtain a 3.12 Å reconstruction of the CCC complex. Curved arrows represent iterative rounds of classification and class selection. Red boxes and text indicate inputs into the subsequent RELION4.0 processing strategy.

**Supplementary Fig. S6.**
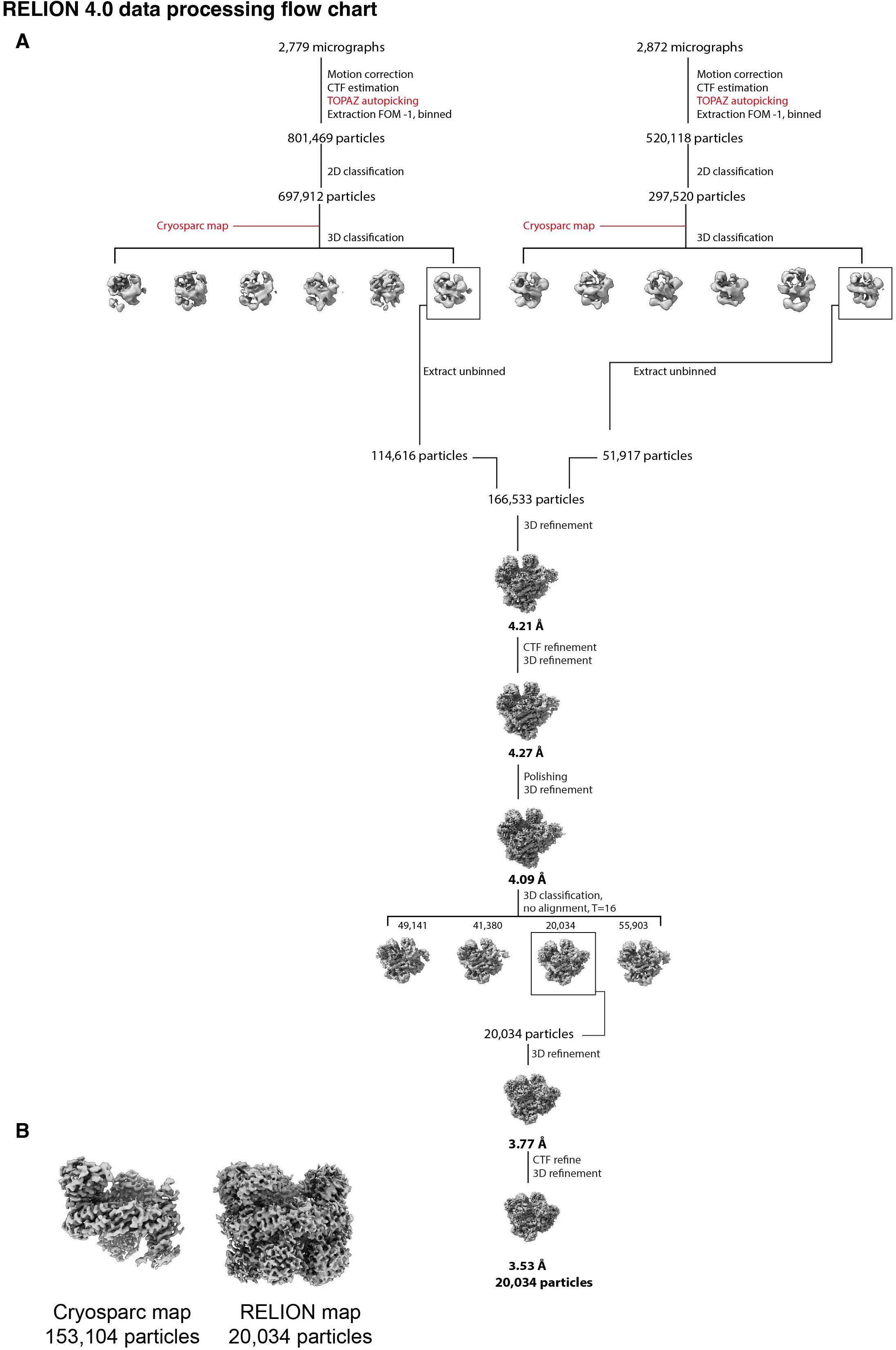
Cryo-EM data analysis of the CCC complex in RELION4.0. (A) Flow chart of the data processing strategy using RELION4.0 to obtain a 3.53 Å reconstruction of the CCC complex. Red text indicates inputs from processing the data in CryoSPARC. **(B)** Representative snapshots of cryoSPARC and RELION reconstructions from ChimeraX. Whilst the cryoSPARC reconstruction has a core of higher resolution, the RELION reconstruction contains better resolved densities for flexible regions.

**Supplementary Fig. S7.**
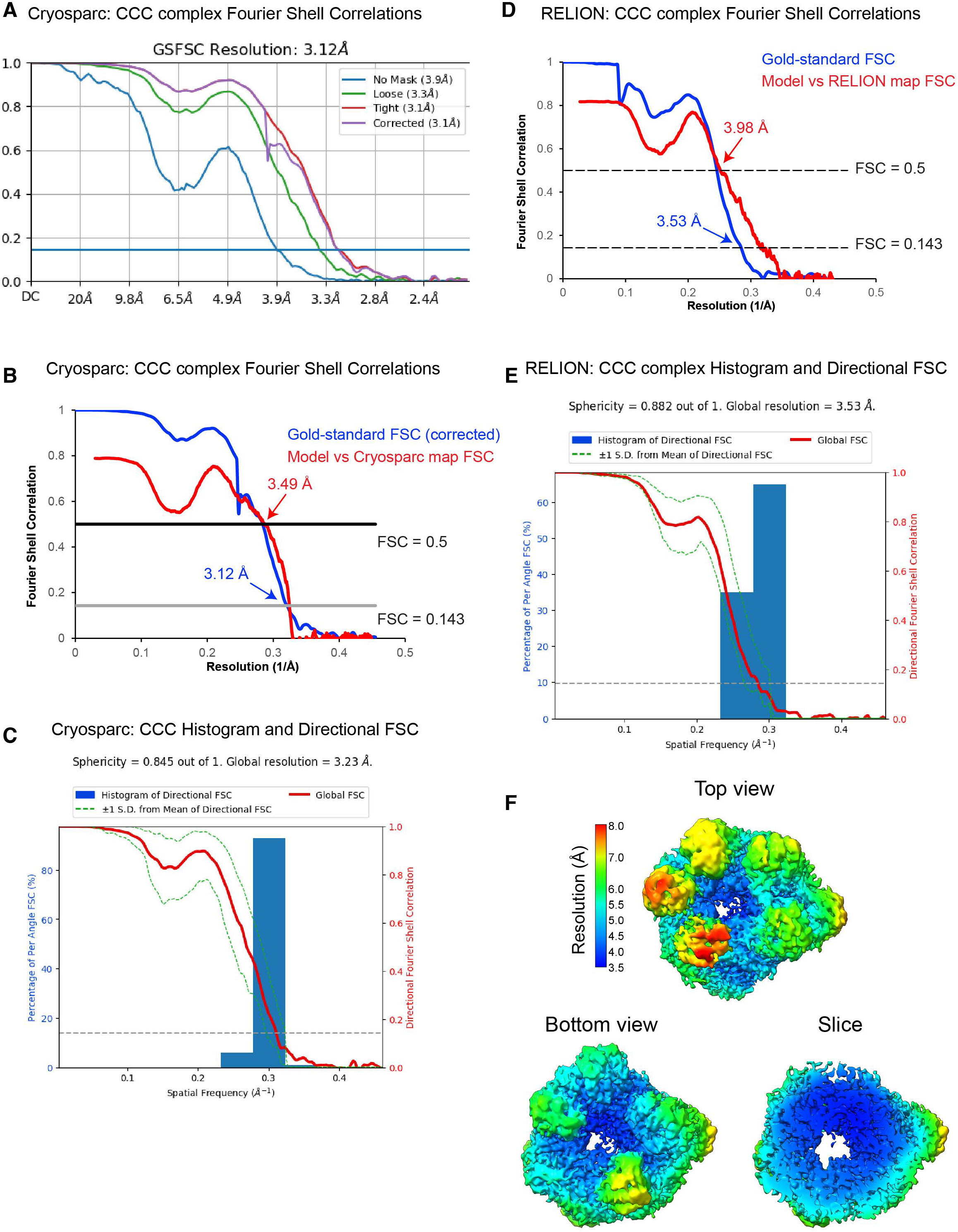
Overall, 3D and local resolution estimations. (A) Gold standard FSC (Fourier Shell Correlation) plots for the CryoSPARC reconstruction. Resolution was estimated at FSC = 0.143. Plot was generated in CryoSPARC. (**B)** Gold-standard (blue) and map vs model (red) FSC plots. Resolution of gold-standard estimated at FSC = 0.143, model-vs-map estimated at FSC = 0.5. **(C)** Directional FSC plots and sphericity values for the CryoSPARC reconstruction. These were calculated using a 3D-FSC server (https://3dfsc.salk.edu/). **(D)** Gold standard (blue) and model-vs-map (red) FSC plots for the RELION4.0 reconstruction. Gold standard resolution was estimated at FSC = 0.143, model-vs-map resolution was estimated at FSC = 0.5. **(E)** Directional FSC plots and sphericity values for the RELION4.0 reconstruction. These were calculated using a 3D-FSC server (https://3dfsc.salk.edu/). **(F)** Local resolution estimates for the RELION4.0 reconstruction. The reconstruction was coloured according to local resolution estimation in RELION.

**Supplementary Fig. S8.**
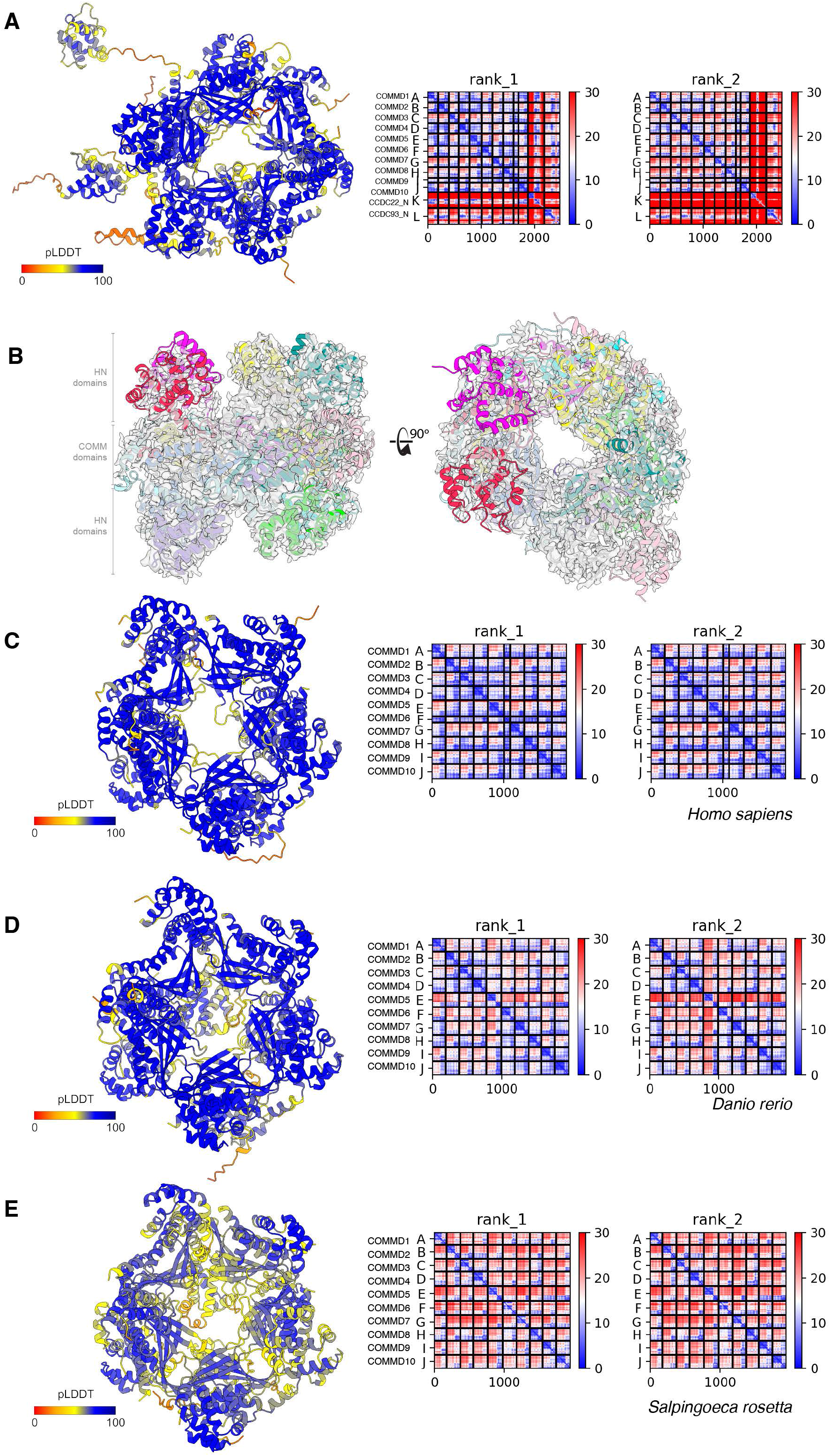
AlphaFold2 modelling of CCC complex across evolution. (A) Alphafold2 model coloured by the confidence metric (pLDDT) of human COMMD1-10 and the N-terminal domains of CCDC22 and CCDC93 with PAE plots of the top 2 ranked models. **(B)** Same view as in Fig. 4A showing the fit of the core CCC subunits to the cryoEM density. Further modelling of the COMMD decamer was conducted using sequences from **(C)** *Homo sapiens*, **(D)** *Danio rerio*, and **(E)** *Salpingoeca rosetta*. Each model displayed highly connected structural correlations between subunits based on PAE plots and consistent and confident decamer assembly.

**Supplementary Fig. S9.**
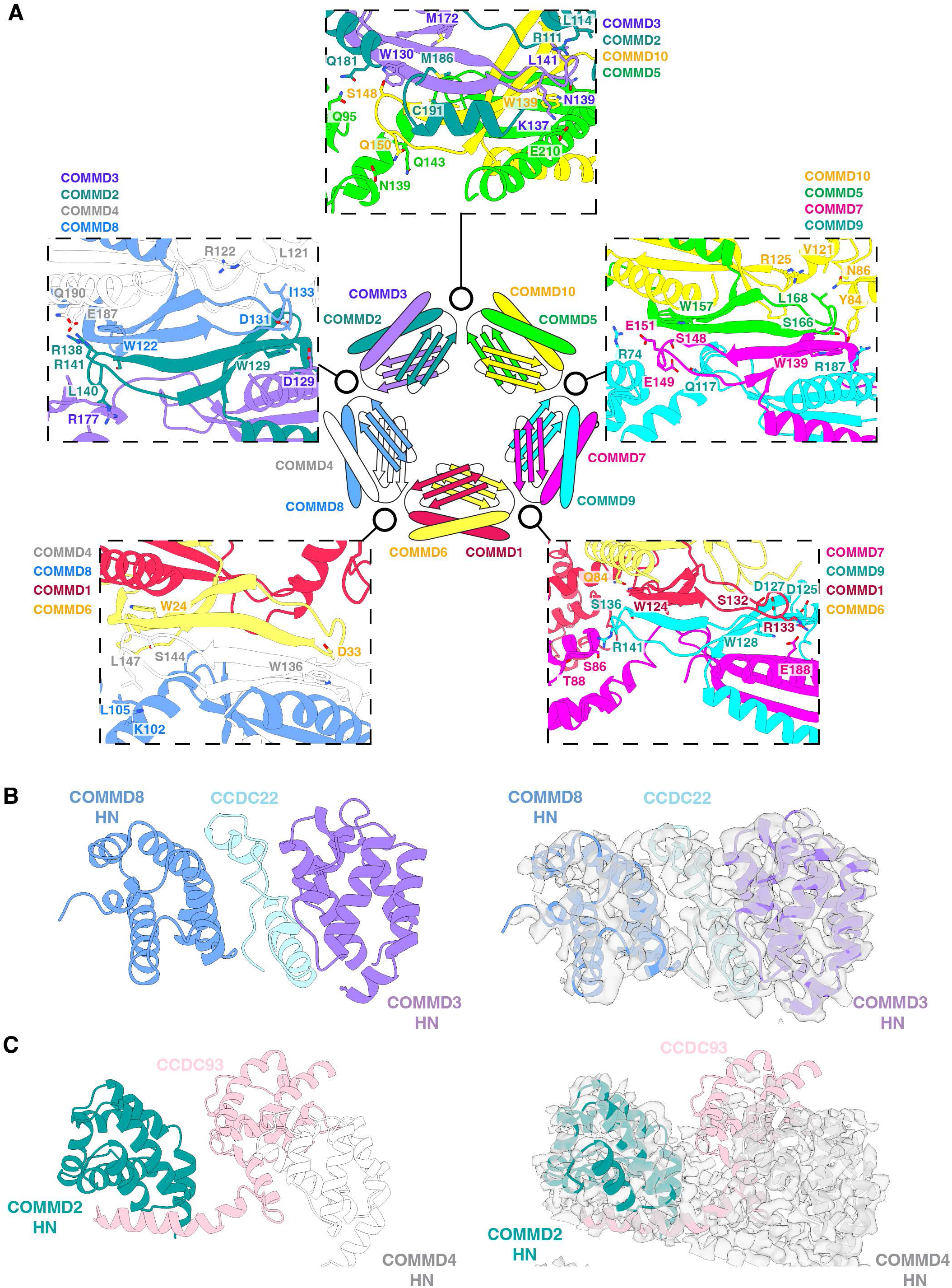
CCDC22 and CCDC93 linkers make extensive contacts with both the central COMM domain ring and the peripheral HN domains. (A) Details of each of the five interfaces between the COMMD heterodimers of the heterodecameric ring. The central schematic is as shown in **Fig. 4D** to provide a reference for each interface. The structural panels show adjacent heterotetramers all shown in the same orientation, placing the strictly conserved Trp sidechain of each subunit as the focal point. Many specific interactions between adjacent subunits determine the precise COMMD organization. **(B).** Interface between CCDC22 and the HN of COMMD3 and COMMD8. **(C)** Interface between CCDC93 and HN domains of COMMD2 and COMMD4.

**Supplementary Fig. S10.**
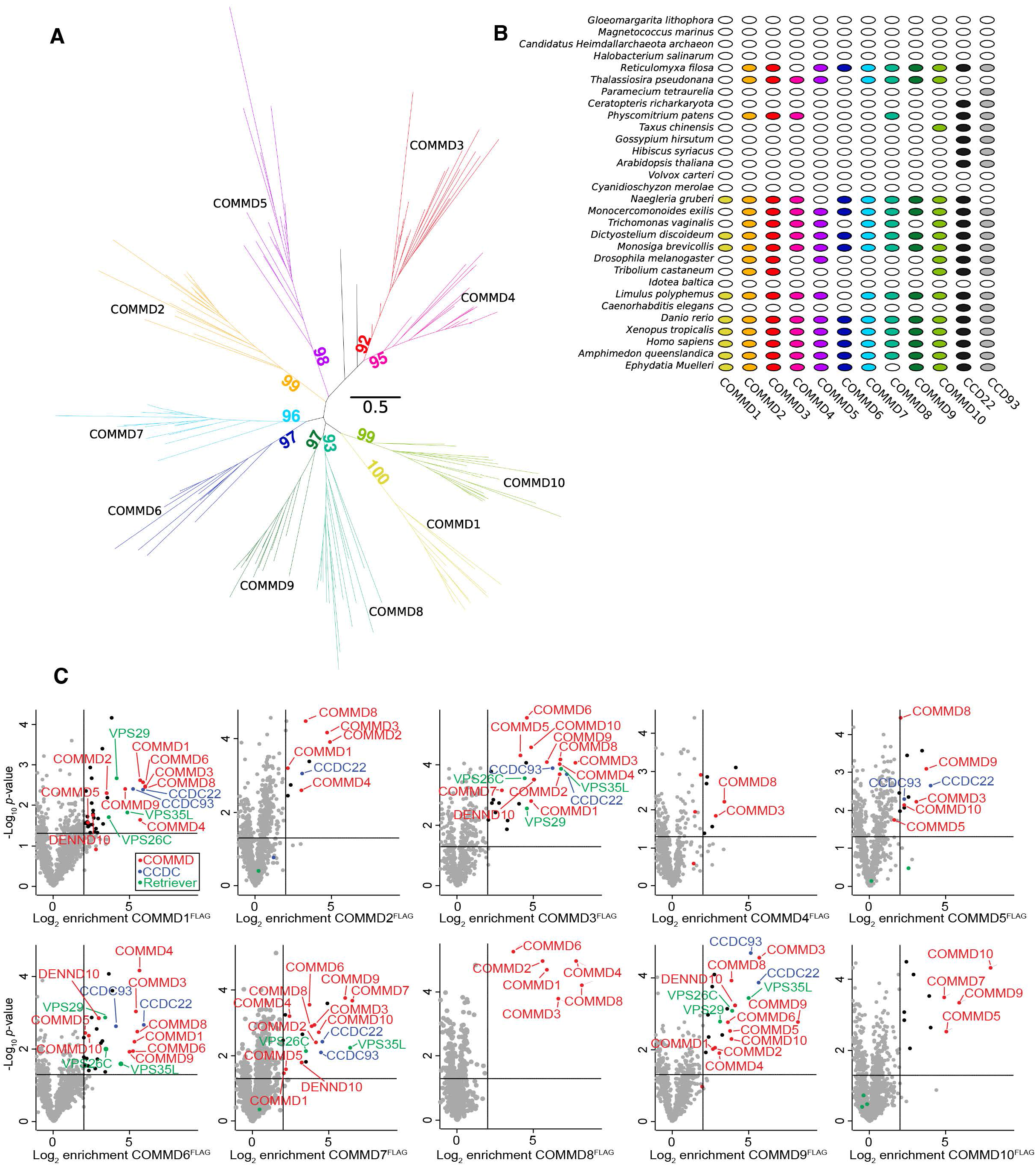
Evolutionary origins of COMMD proteins and the CCC complex. (A). A maximum-likelihood phylogeny of COMMD1-10 proteins from 23 representative eukaryotic taxa inferred under the best-fitting Q.yeast+R5 substitution model. Each COMMD subunit forms a strongly supported (>90% bootstrap) clan in the unrooted phylogeny, and each clan contains representatives from all of the major lineages (supergroups) of eukaryotes; this implies that all ten COMMD subunits were already present in the last eukaryotic common ancestor (that is, LECA appears 10 times in the tree). Based on the absence of COMMD homologues in Bacteria and Archaea, this protein family likely originated on the eukaryotic stem and proliferated via a series of gene duplications prior to the radiation of the modern eukaryotic groups. **(B).** Presence-absence patterns of COMMD and CCDC22/CCDC93 in a set of representative modern eukaryotes. The presence-absence pattern of COMMD genes in modern eukaryotes, taken together with the phylogeny in (A), indicates that these genes have been lost multiple times independently in different eukaryotic lineages. **(C).** Digitonin-solubilized COMMD knockout eHap cell lines expressing the indicated COMMD^FLAG^ construct were subjected to affinity enrichment using Flag agarose beads followed by label free quantitative proteomics. The threshold of significant enrichment was determined to be 2-fold (log2 fold change = 1) based on the distribution of unenriched proteins. Black and colored dots indicate significantly enriched proteins. Red, COMMD subunits; Blue, CCDC complex subunits; Green, Retriever subunits.

**Supplementary Figure S11.**
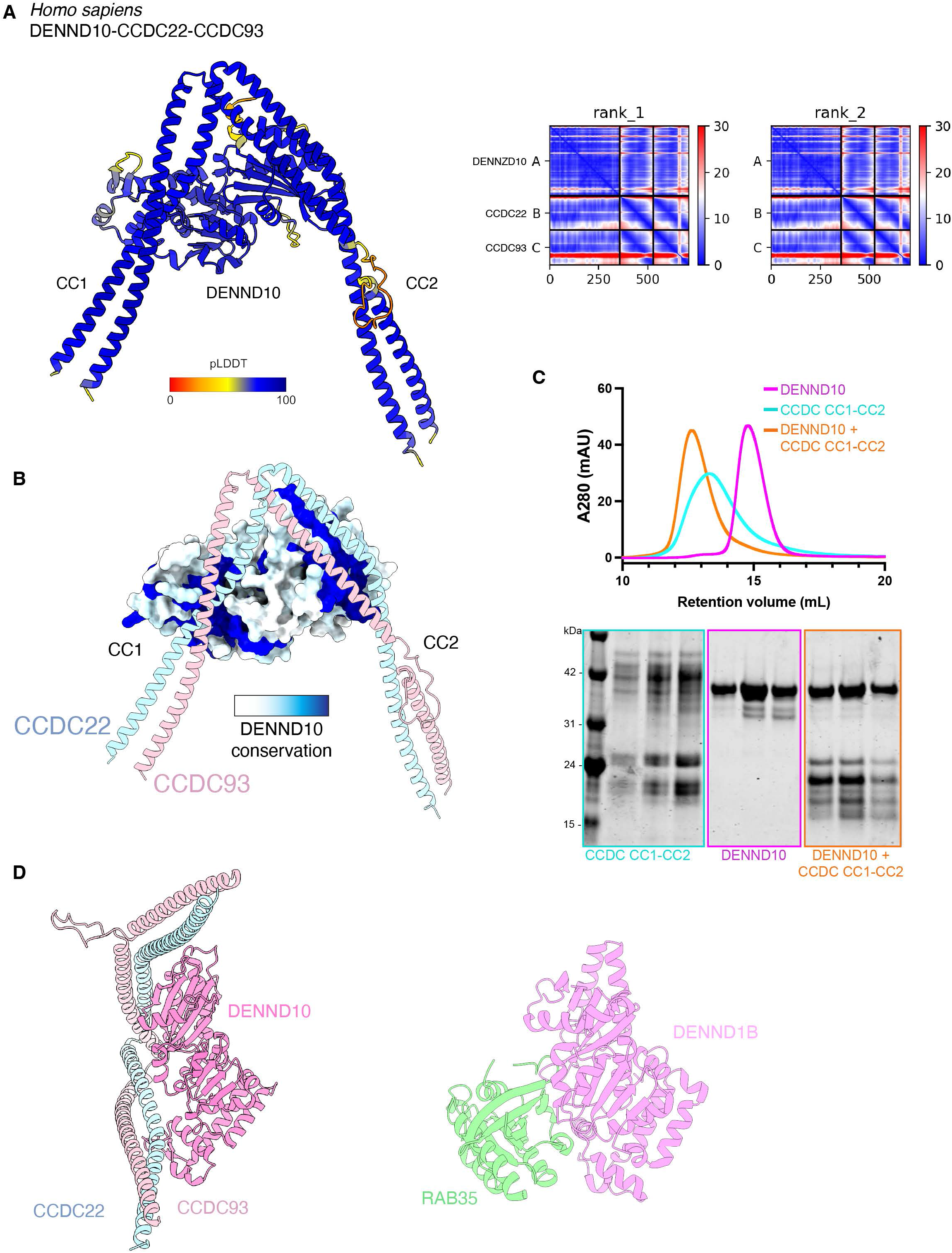
AlphaFold2 modelling of the DENND10-CCDC22-CCDC93 complex. (A) Alphafold2 model of human DENND10 and the central CC1-CC2 coiled-coil domains of CCDC22 and CCDC93 coloured by the confidence metric (pLDDT). The PAE plots of the top 2 ranked models are shown on the right. **(B)** Same model as in (A) but with DENND10 shown in surface representation coloured according to conservation with CONSURF (134). **(C)** Top panel shows analytical size exclusion chromatography of DENND10 (magenta), CCDC22-CCDC93 CC1-CC2 complex (cyan) and DENND10 mixed with the CCDC22-CCDC93 forming a stable complex (orange). The bottom panel shows Coomassie stained gel of the peak fractions for each sample. (**D**) The predicted structure of DENND10 bound to CCDC22-CCDC93 is shown in the same orientation alongside the crystal structure of DENND1B in complex with RAB35 (PDB ID: 3TW8)(74).

**Supplementary Figure S12.**
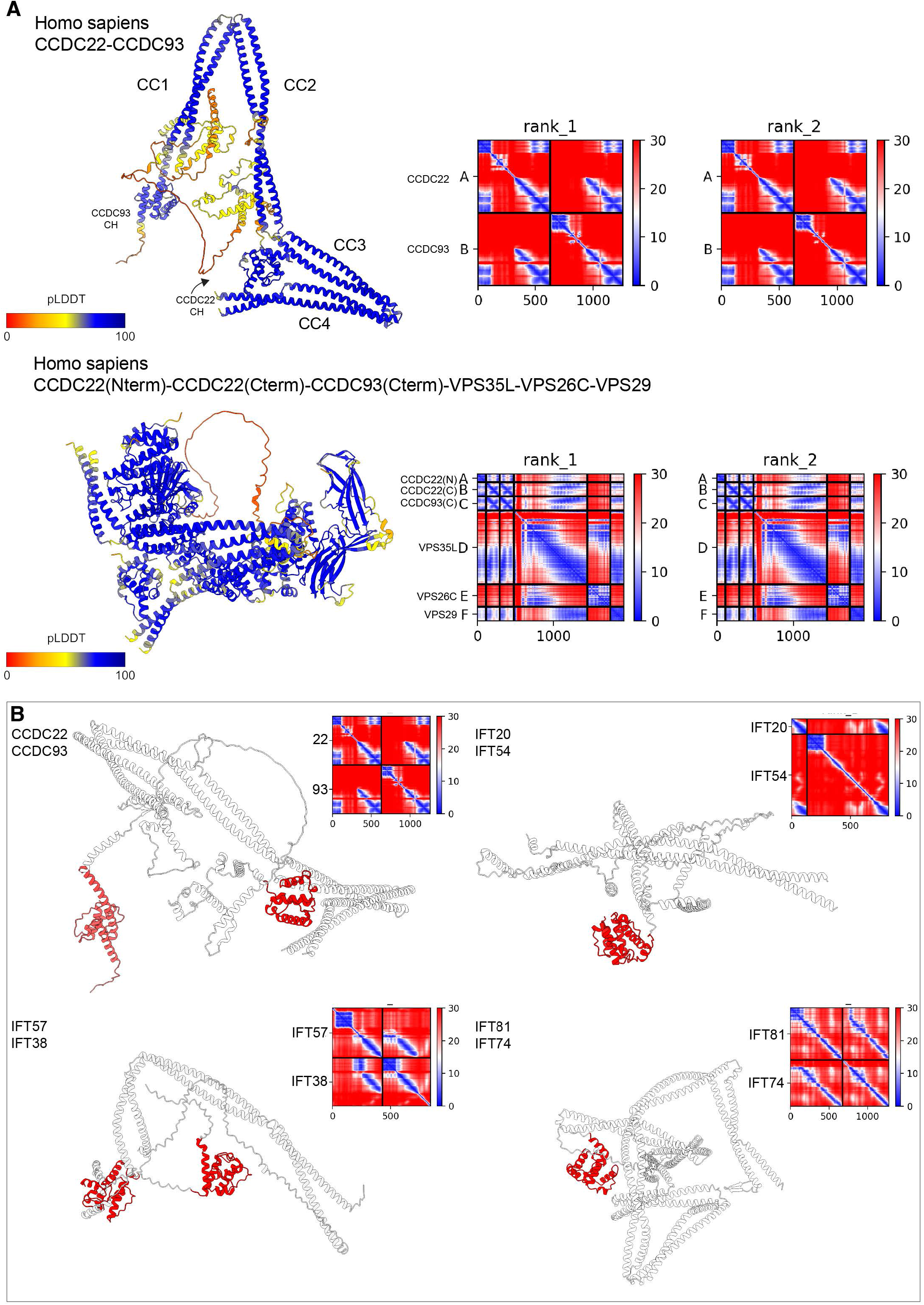
CCDC22 and CCDC93 heterodimeric coiled-coil share similarity with various IFT subunits of the intraflagellar transport machinery. AlphaFold2 models of full length CCDC22 and CCDC93 (*top*) as well as truncated CCDC22 and CCDC93 in complex with Retriever (*bottom*) show the formation of a highly conserved coiled-coil structure able to interact with Retriever. **(B)** Highlights the similarity between CCDC22 and CCDC93 and IFT proteins. Each of these proteins contain a calponin homology (CH) domain (red) and an extended coiled-coil region. Previously CH domains have been shown to bind both actin and microtubules which could suggest a similar function for the CH domains of CCDC22 and CCDC93.

**Supplementary Figure S13.**
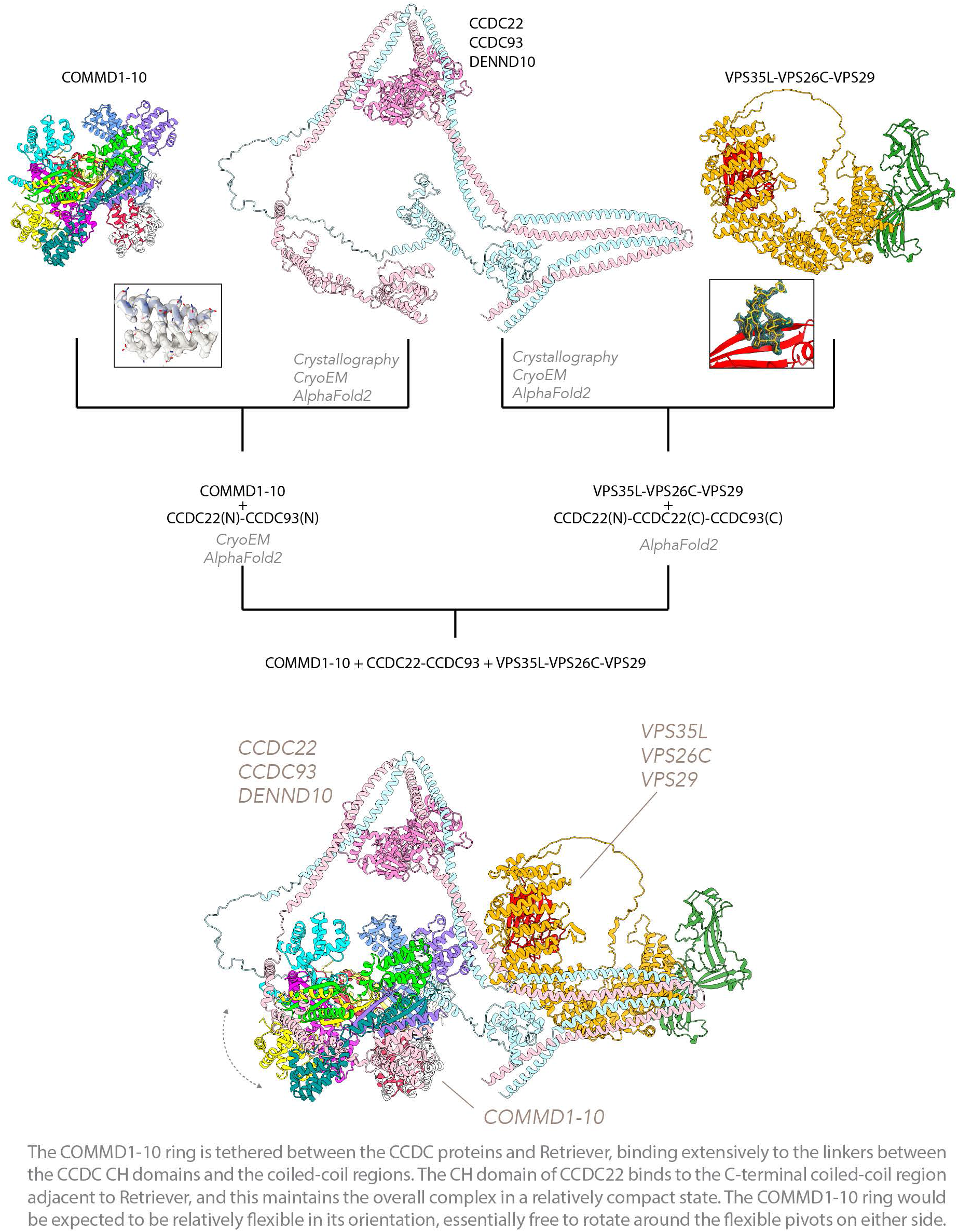
Workflow for assembling the complete Commander complex combining experimental structures and AlphaFold2 modelling. A flow chart showing the various methodologies used to assemble the Commander complex. We used a combination of AlphaFold2 modelling,X-ray crystallographic structures and cryoEM to establish models of Retriever and the CCC complex. Using AlphaFold2 it was possible to model CCDC22 and CCDC93, which act as a bridge between the two complexes. The final model reveals that the COMMD1-10 ring is tethered between the CCDC proteins and Retriever, binding extensively to the linkers between the CCDC CH domains and the coiled-coil regions. The CH domain of CCDC22 binds to the C-terminal coiled-coil region adjacent to Retriever, and this maintains the overall complex in a relatively compact state. The COMMD1-10 ring would be expected to be relatively flexible in its orientation, essentially free to rotate around the flexible pivots on either side. As outlined in the text, all key interfaces have been experimentally validated by mutagenesis, co-immunoprecipitation and cellular rescue studies.

**Supplementary Figure S14.**
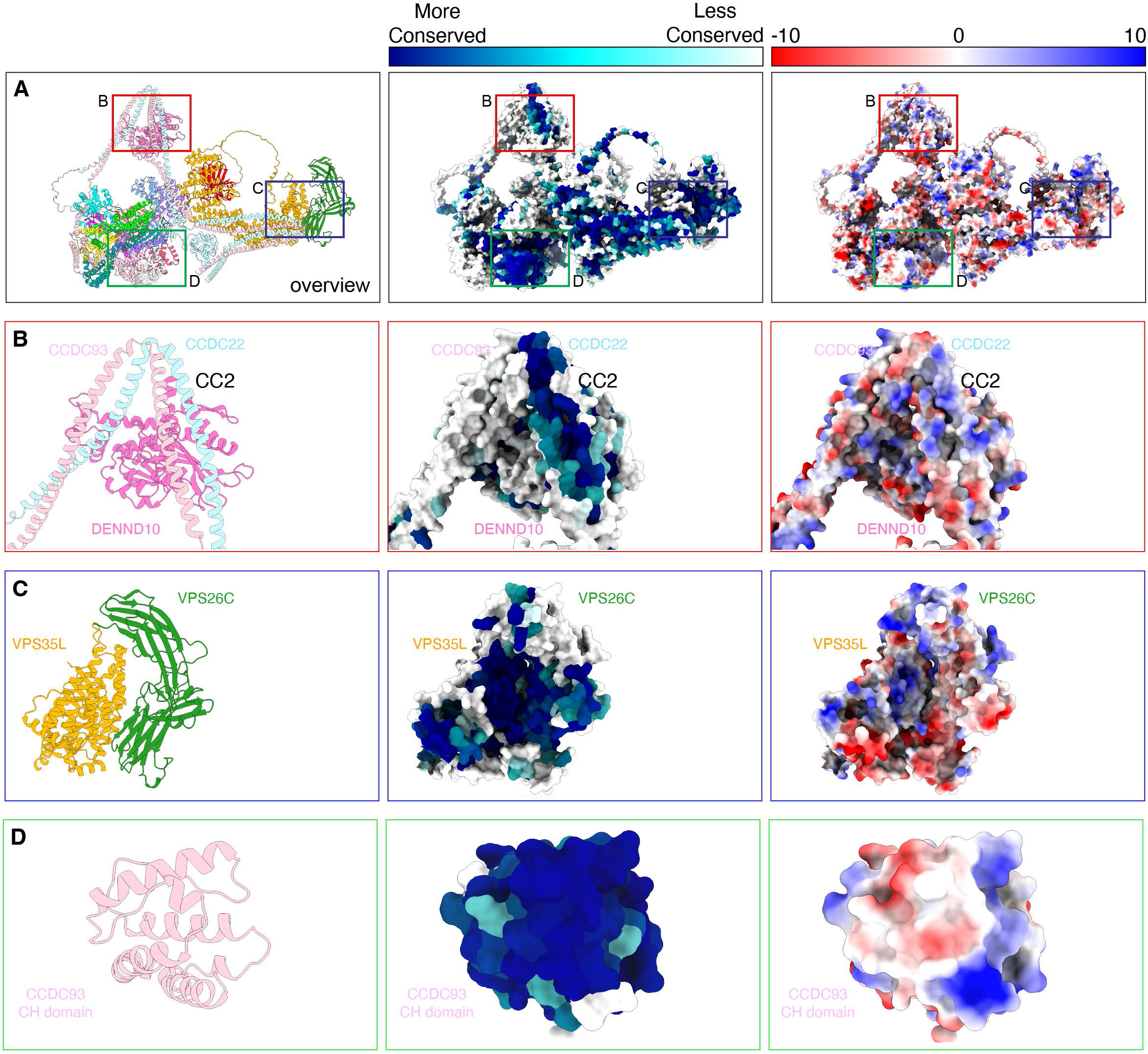
Conserved and electrostatic surfaces of the modelled Commander complex. (A) An overview of the Commander complex shown as a ribbon diagram (left), with conserved surfaces mapped with Consurf (middle) (134) and with electrostatic surface potential calculated with ChimeraX (127). (**B**) Close up view of the CCDC22-CCDC93-Dennd10 interface where a conserved surface aligns with the previously identified region for binding of the WASH complex subunit Fam21 (30). (**C**) Close up of the interface between VPS35L-VPS26C where a conserved pocket is found. Previous studies indicate that VPS26C is required for SNX17 interaction (6). (**D**) Close up of the of the CCDC93 N-terminal CH domain showing its very highly conserved surface properties.

**Supplementary Figure S15.**
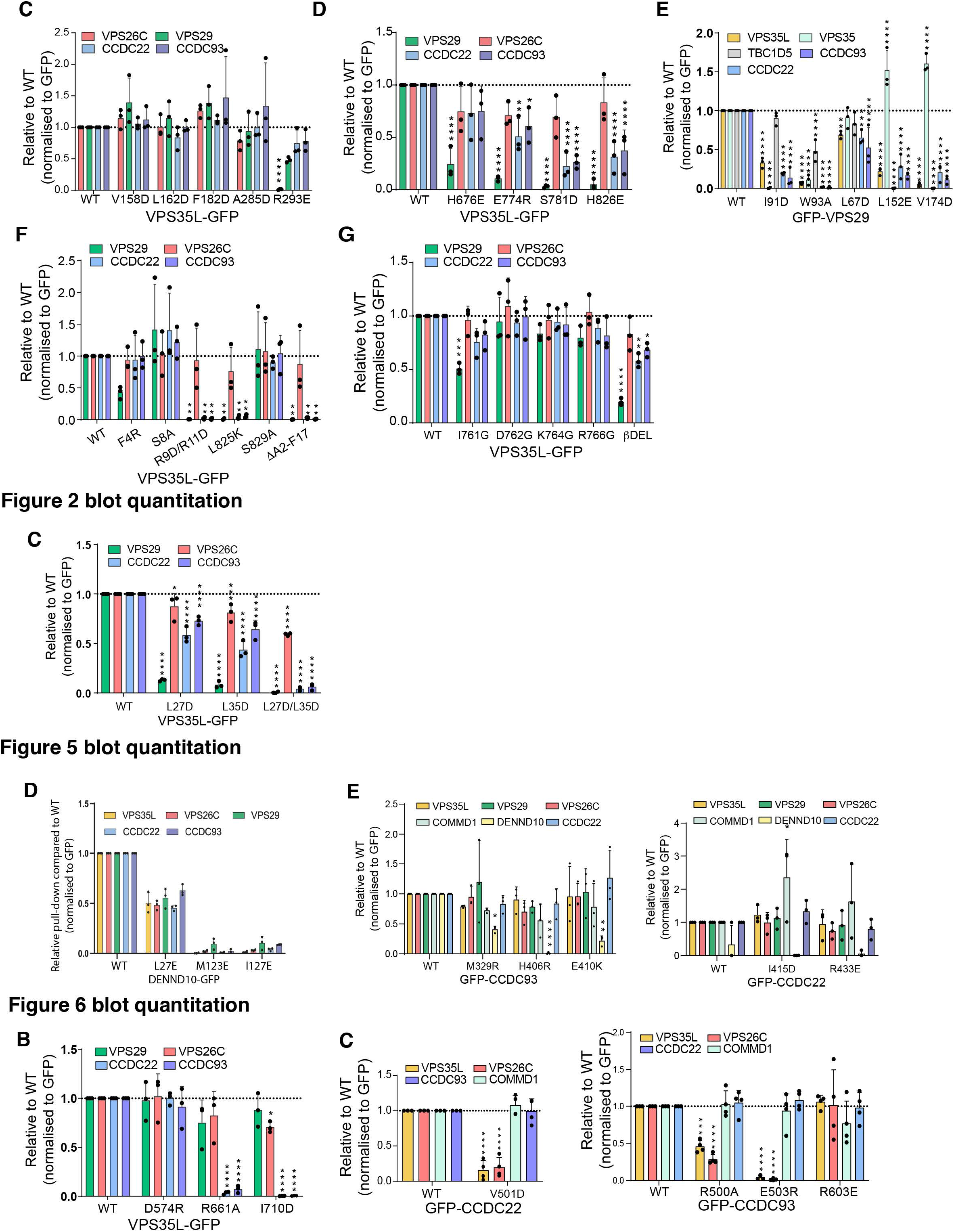
Quantified band intensities from all western blots in this study. Band intensities were quantified using Odyssey software and normalized to GFP expression. 2-way ANOVA with Dunnett’s multiple comparisons test. Error bars represent the mean, s.d.. *P < 0.05, **P < 0.01, ***P < 0.001, ****P < 0.0001.

**Supplementary Figure S16.**
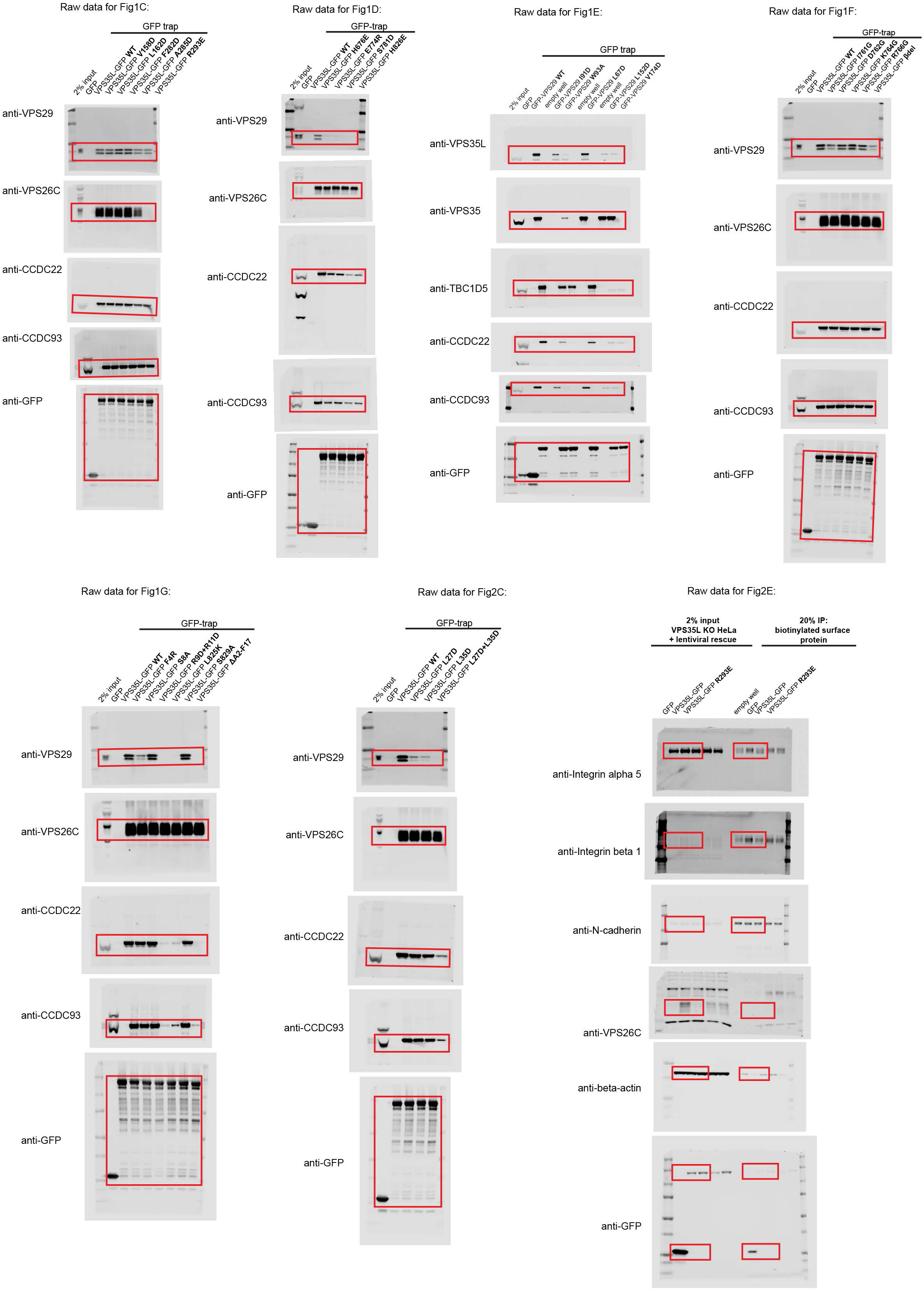
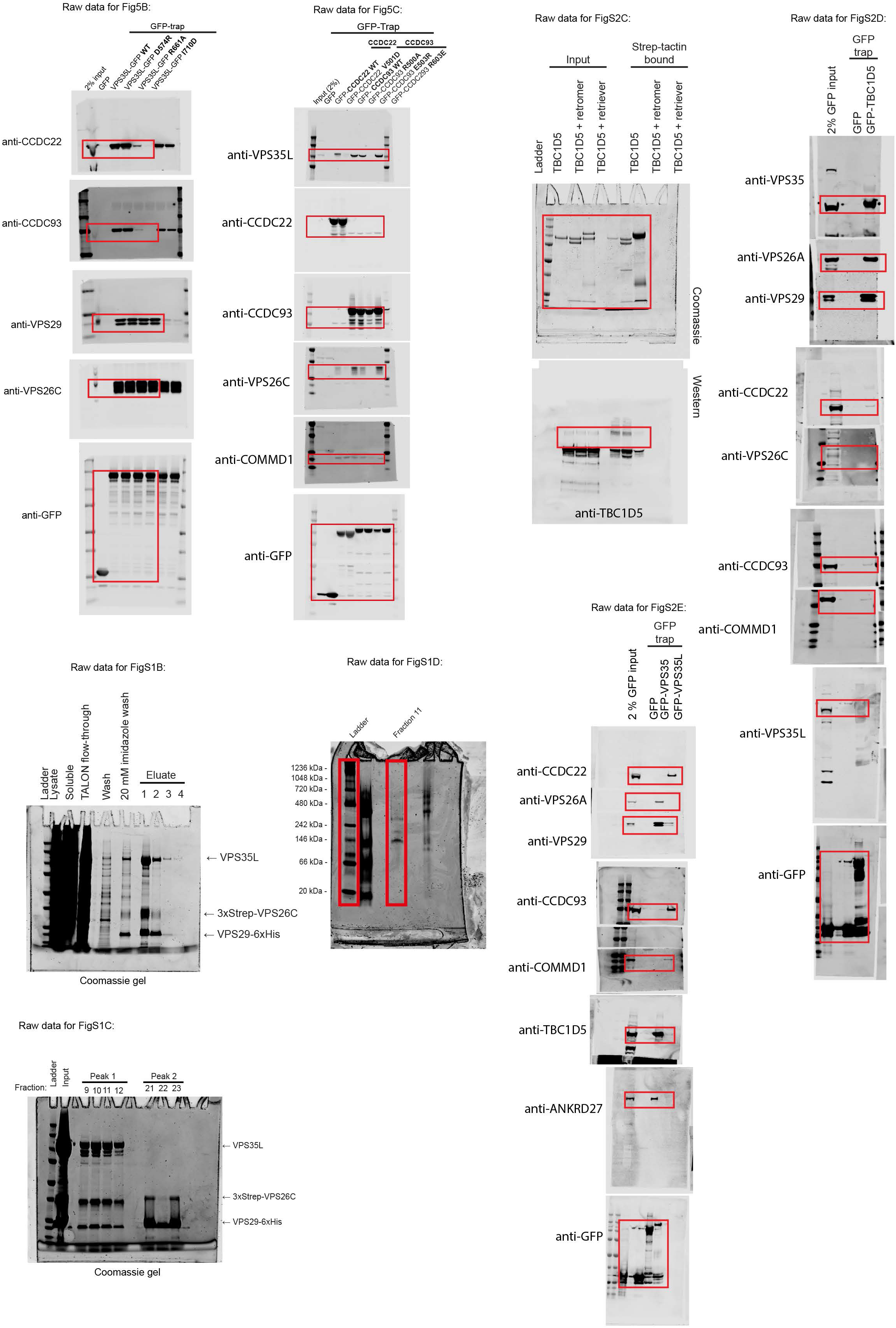
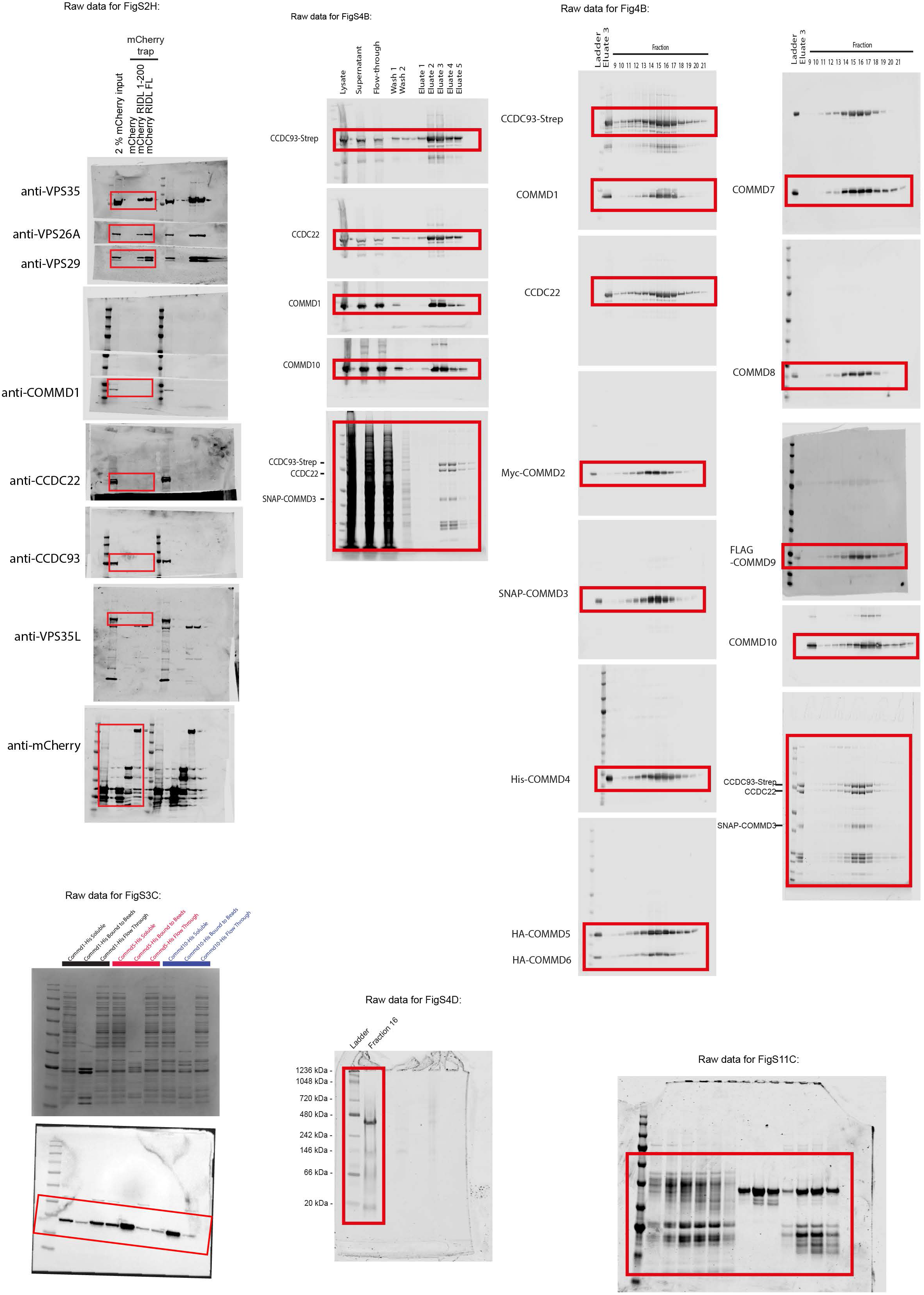
**Raw images of gels and western blots included in this study.**

**Table S1.** Cryo-EM data collection, refinement, and validation statistics.

|  | **Retriever**<br>VPS35L-VPS26C-VPS29 | **CCC**<br>COMMD1 to COMMD10-CCDC22-CCDC93 |  |
| --- | --- | --- | --- |
| **Data collection and processing** |  |  |  |
| Magnification | 130,000 | 75,000 |  |
| Voltage (kV) | 200 | 300 |  |
| Electron exposure (e<sup>-</sup>/Å<sup>2</sup>) Defocus range (µm) | 61.7 e<sup>-</sup>/Å<sup>2</sup><br>-2, -1.6, -1.2, -0.8 µm | 43.55/42.73 e<sup>-</sup>/Å<sup>2</sup><br>-2, -1.8, -1.6, -1.4, -1.2 µm |  |
| Pixel size (Å) | 0.525 | 1.084 |  |
| Symmetry imposed | C1 | C1 |  |
| Programme used | RELION/cryoSPARC | cryoSPARC | RELION |
| Initial particle images (no.) | 297,077 | 2,666,858 | 1,321,587 |
| Final particle images (no.) | 119,564 | 153,104 | 20,034 |
| Map resolution (Å) | 4. | 3.12 | 3.53 |
| FSC threshold | 0.143 | 0.143 | 0.143 |
| **Refinement** |  |  |  |
| Initial model used |  | Colabfold | CryoSPARC/Colabfold |
| Model resolution range (Å) |  | 3.49 | 3.98 |
| FSC threshold |  | 0.5 | 0.5 |
| Map sharpening *B* factor (Å<sup>2</sup>) |  | 83 | - 24 |
| Model composition |  |  |  |
| Non-hydrogen atoms |  | 29276 | 36574 |
| Protein residues |  | 1837 | 2298 |
| R.m.s. deviations |  |  |  |
| Bond lengths (Å) |  | 0.005 | 0.002 |
| Bond angles (°) |  | 1.118 | 0.561 |
| Validation |  |  |  |
| MolProbity score |  | 1.41 | 1.49 |
| Clashscore |  | 6.01 | 6.24 (90<sup>th</sup> percentile) |
| Poor rotamers (%) |  | 0.12 | 0 |
| Ramachandran plot |  |  |  |
| Favored (%) |  | 97.63 | 97.18 |
| Outliers (%) |  | 0.17 | 0 |
| EMDB ID |  | EMD-28825 | EMD-28827 |
| PDB ID |  | 8F2R | 8F2U |

**Table S2.** Crystal structure data collection, refinement and validation statistics^a^.

|  | COMMD5-COMMD7-COMMD9-COMMD10 | VPS29-VPS35L peptide |
| --- | --- | --- |
| **Data Collection** |  |  |
| Space group | C2 | P212121 |
| Unit cell dimensions (a,b,c) (Å) | 215.19 74.89 63.63 | 43.72 55.28 82.64 |
| Unit cell angles ($\alpha,\beta,\gamma$) (°) | 90.00 96.40 90.00 | 90.00 90.00 90.00 |
| Wavelength (Å) | 0.95372 | 0.95365 |
| Resolution range (Å) | 48.02-3.33 (3.60-3.33) | 45.95-1.35 (1.37-1.35) |
| Total reflections | 102614 (20743) | 277842 (11606) |
| Unique reflections | 14773 (2987) | 44459 (1992) |
| Multiplicity | 6.9 (6.9) | 6.2 (5.8) |
| Completeness (%) | 99.0 (97.6) | 99.1 (91.6) |
| Mean I/$\sigma$(I) | 8.9 (1.3) | 11.3 (1.0) |
| R-merge | 0.108 (1.088) | 0.07 (1.47) |
| R-meas | 0.127 (1.278) | 0.077 (1.61) |
| R-pim | 0.048 (0.480) | 0.031 (0.643) |
| CC(1/2) | 0.999 (0.732) | 0.998 (0.481) |
| **Refinement** |  |  |
| R-work | 0.243 (0.318) | 0.219 (0.400) |
| R-free | 0.287 (0.404) | 0.246 (0.382) |
| Number of non-hydrogen atoms | 4157 | 1770 |
| Number macromolecule atoms | 4157 | 1576 |
| Number water atoms | 0 | 194 |
| RMS bonds (Å) | 0.002 | 0.023 |
| RMS angles (°) | 0.519 | 1.95 |
| Ramachandran favored (%) | 96.5 | 97.9 |
| Ramachandran allowed (%) | 3.5 | 2.1 |
| Ramachandran outliers (%) | 0 | 0 |
| Clashscore | 2.97 | 9.8 |
| Overall Molprobit score | 1.11 | 1.7 |
| Average B-factor | 141.4 | 26.7 |
| PDB ID | 8ESD | 8ESE |

a. Statistics for the highest-resolution shell are shown in parentheses.

Table S3. Primary antibodies used in this study.

**Table S3. Primary antibodies used in this study.**
| Primary antibody | Species | Use and dilution | Supplier | Catalogue number |
| --- | --- | --- | --- | --- |
| CCDC22 | Rabbit | WB 1/1000 | Proteintech | 16636-1-AP |
| CCDC93 | Mouse | WB 1/1000 | LS Bio | C336997 |
| CCDC93 | Rabbit | WB 1/1000 | Proteintech | 20861-1-AP |
| EEA1 | Mouse | IF 1/200 | BD Biosciences | 610457 |
| FAM21 | Rabbit | WB 1/1000<br>IF 1/1000 | Gift from Dan Billadeau | N/A |
| GFP | Mouse | WB 1/2000 | Roche | 11814460001 |
| Integrin α5 | Mouse | WB 1/1000 | BD Biosciences | 555651 |
| Integrin α5 | Rabbit | IF 1/200 | Abcam | EPR7854 |
| Integrin β1 | Mouse | WB 1/1000 | BD Biosciences | 610467 |
| LAMP1 | Mouse | IF 1/400 | Developmental Studies Hybridoma Bank | H4A3 |
| VPS26A | Rabbit | WB 1/1000 | Abcam | ab137447 |
| VPS26C | Rabbit | WB 1/2000 | Merck Millipore | ABN87 |
| VPS29 | Rabbit | WB 1/500 | Abcam | ab98929 |
| VPS29 | Mouse | WB 1/200 | Santa Cruz | 398874 |
| VPS35 | Rabbit | WB 1/1000 | Abcam | ab157220 |
| VPS35 | Rabbit | IF 1/200 | Abcam | ab97545 |
| VPS35 | Goat | IF 1/200 | Abcam | ab10099 |
| VPS35L | Rabbit | WB 1/1000<br>IP 2μg | Abcam | ab97889 |
| VPS35L | Rabbit | IF 1/400 | Pierce | PA5-28553 |
| WASH1 | Rabbit | WB 1/1000<br>IF 1/1000 | Gift from Dan Billadeau | N/A |
| TBC1D5 | Rabbit | WB 1/1000 | Abcam | 203896 |
| ANKRD27 |  |  |  |  |
| GFP | Mouse | WB 1/2000 | Roche | 11814460001 |
| N-cadherin | Mouse | WB 1/1000 | Cell Signalling | 14215 |
| N-cadherin | Rabbit | WB 1/1000 | Cell Signalling | 13116 |
| $\beta$ actin | Mouse | WB 1/2000 | Sigma | A1978 |
| COMMD1 |  |  | Proteintech | 11938-1-AP |
| COMMD1 | Mouse | WB 1/1000<br>IF 1/700 | Sigma | WH0150684M1 |
| COMMD2 | Rabbit | WB 1/1000 | Millipore Sigma | HPA044190 |
| COMMD3 | Rabbit | WB 1/1000 | Abcam | ab176583 |
| COMMD4 | Rabbit | WB 1/1000 | Abcam | ab115169 |
| COMMD5 | Mouse | WB 1/1000 | Proteintech | 67043-1-Ig |
| COMMD7 | Rabbit | WB 1/1000 | Abcam | ab96091 |
| COMMD7 | Rabbit | WB 1/1000 | GeneTex | GTX112076 |
| COMMD8 | Rabbit | WB 1/1000 | Proteintech | 25237-1-AP |
| COMMD8 | Mouse | WB 1/1000 | Santa Cruz | sc-3973869 |
| COMMD9 | Rabbit | WB 1/1000 | Abcam | Ab121303 |
| COMMD10 | Mouse | WB 1/1000 | Santa Cruz | sc-398798 |
| COMMD10 | Rabbit | WB 1/1000 | GeneTex | GTX121488 |
| HA-tag | Mouse | WB 1/1000 | Insight Biotechnology | A02040 |
| Strep-tag | Rabbit | WB 1/1000 | GeneTex | GTX128061 |
| 6x His-tag | Rabbit | WB 1/1000 | Abcam | Ab18184 |
| Myc-tag | Rabbit | WB 1/1000 | GeneTex | GTX115046 |
| FLAG-tag | Mouse | WB 1/1000 | Sigma | F3165 |
| SNAP-tag | Rabbit | WB 1/1000 | GeneTex | GTX54523 |

**Table S4.** Secondary antibodies used in this study.

| **Secondary antibody** | **Species** | **Use and dilution** | **Supplier** | **Catalogue number** |
| --- | --- | --- | --- | --- |
| AlexaFluor 680 anti-mouse IgG | Donkey | WB 1/20000 | Life Technologies | A10038 |
| AlexaFluor 800 anti-rabbit IgG | Donkey | WB 1/20000 | Life Technologies | W10824 |
| Goat anti-rabbit IgG | Goat | WB 1/5000 | Life Technologies | AB2533967 |
| AlexaFluor 488 anti-mouse IgG | Donkey | IF 1/300 | Life Technologies | A21202 |
| AlexaFluor 568 anti-mouse IgG | Donkey | IF 1/300 | Life Technologies | A10037 |
| AlexaFluor 647 anti-mouse IgG | Donkey | IF 1/300 | Life Technologies | A31571 |
| AlexaFluor 488 anti-rabbit IgG | Donkey | IF 1/300 | Life Technologies | A21206 |
| AlexaFluor 568 anti-rabbit IgG | Donkey | IF 1/300 | Life Technologies | A10042 |
| AlexaFluor 647 anti-rabbit IgG | Donkey | IF 1/300 | Life Technologies | A31573 |
| AlexaFluor 488 anti-goat IgG | Donkey | IF 1/300 | Life Technologies | A11055 |
| AlexaFluor 568 anti-goat IgG | Donkey | IF 1/300 | Life Technologies | A11057 |
| AlexaFluor 647 anti-goat IgG | Donkey | IF 1/300 | Life Technologies | A21447 |

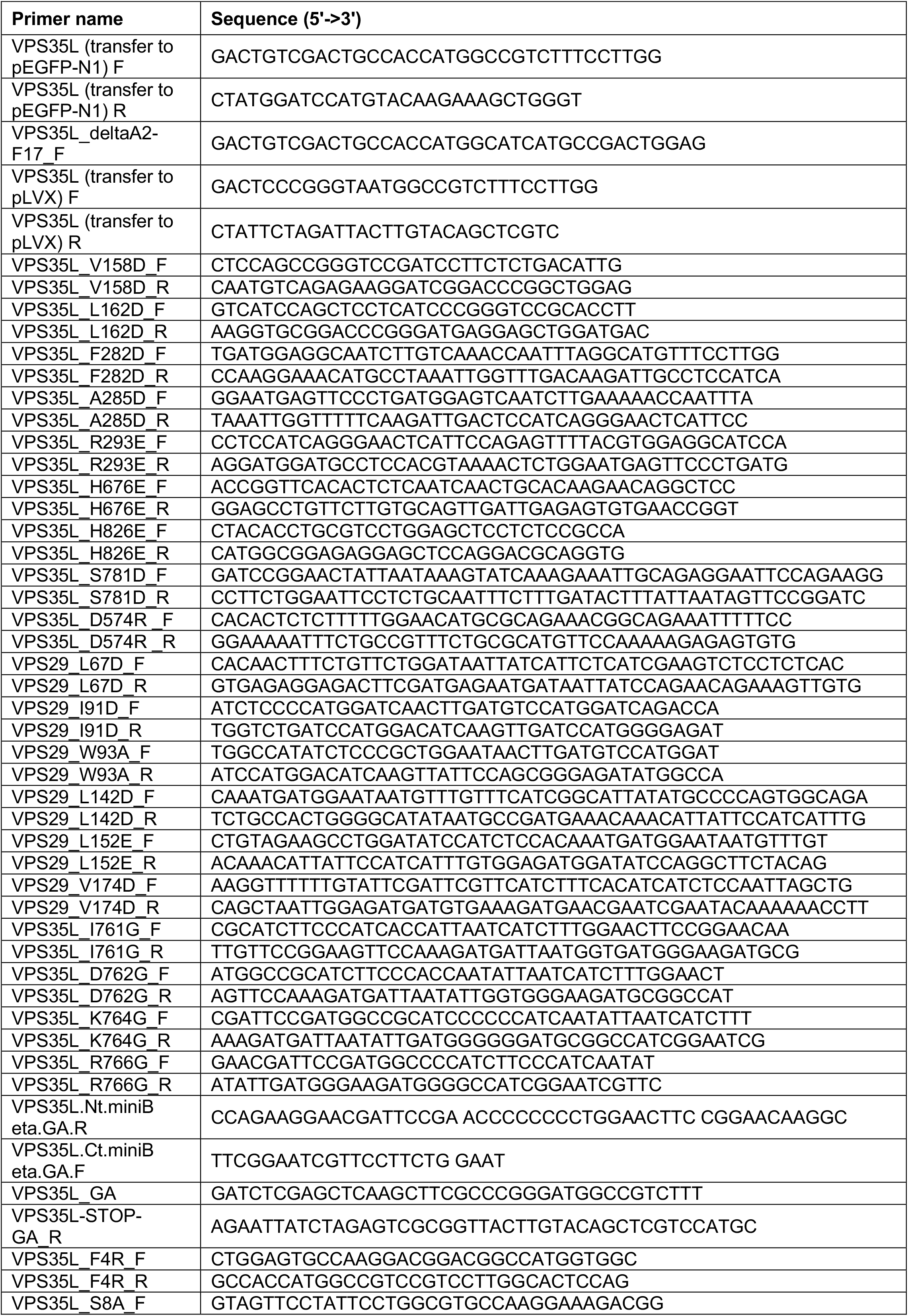
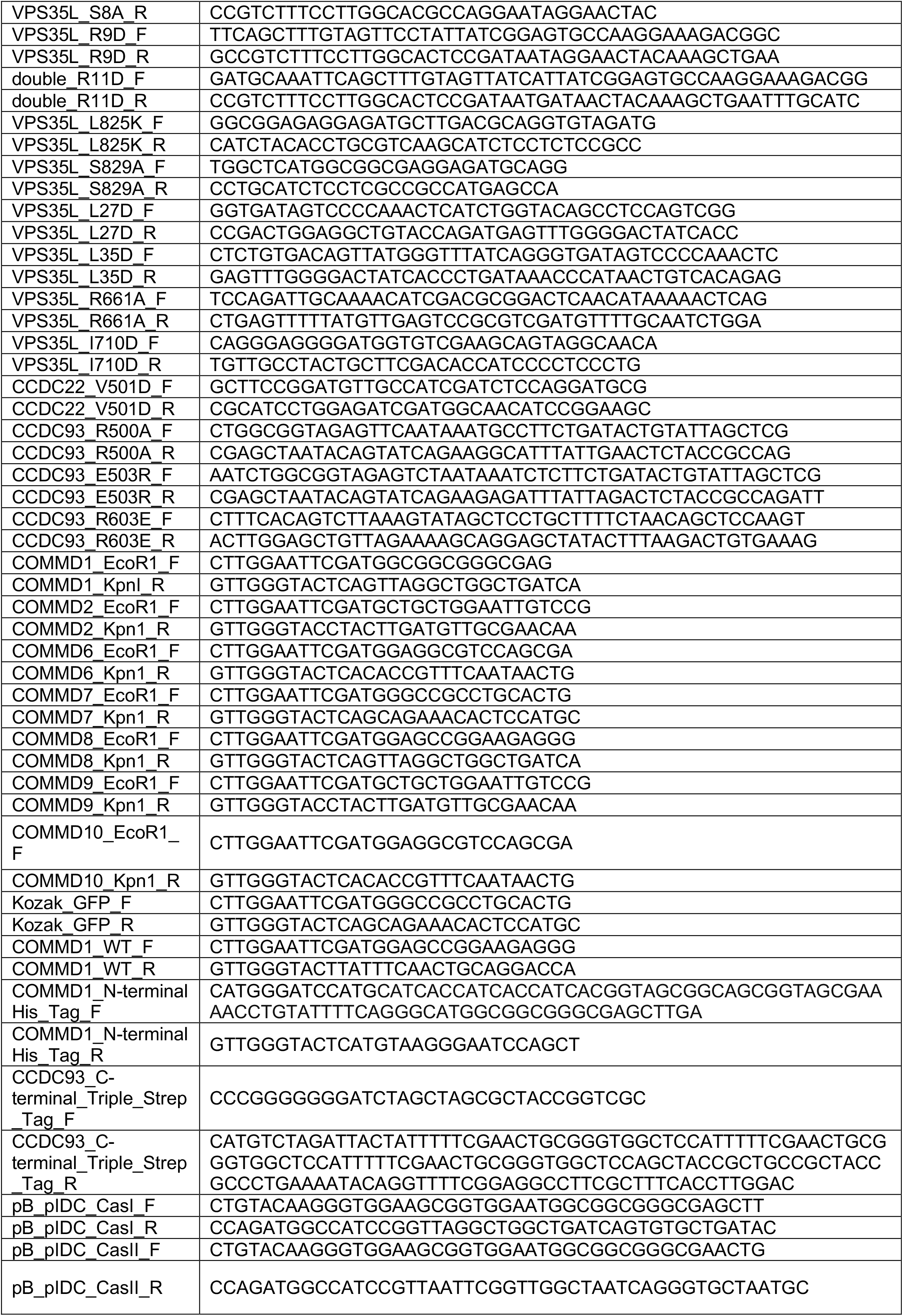
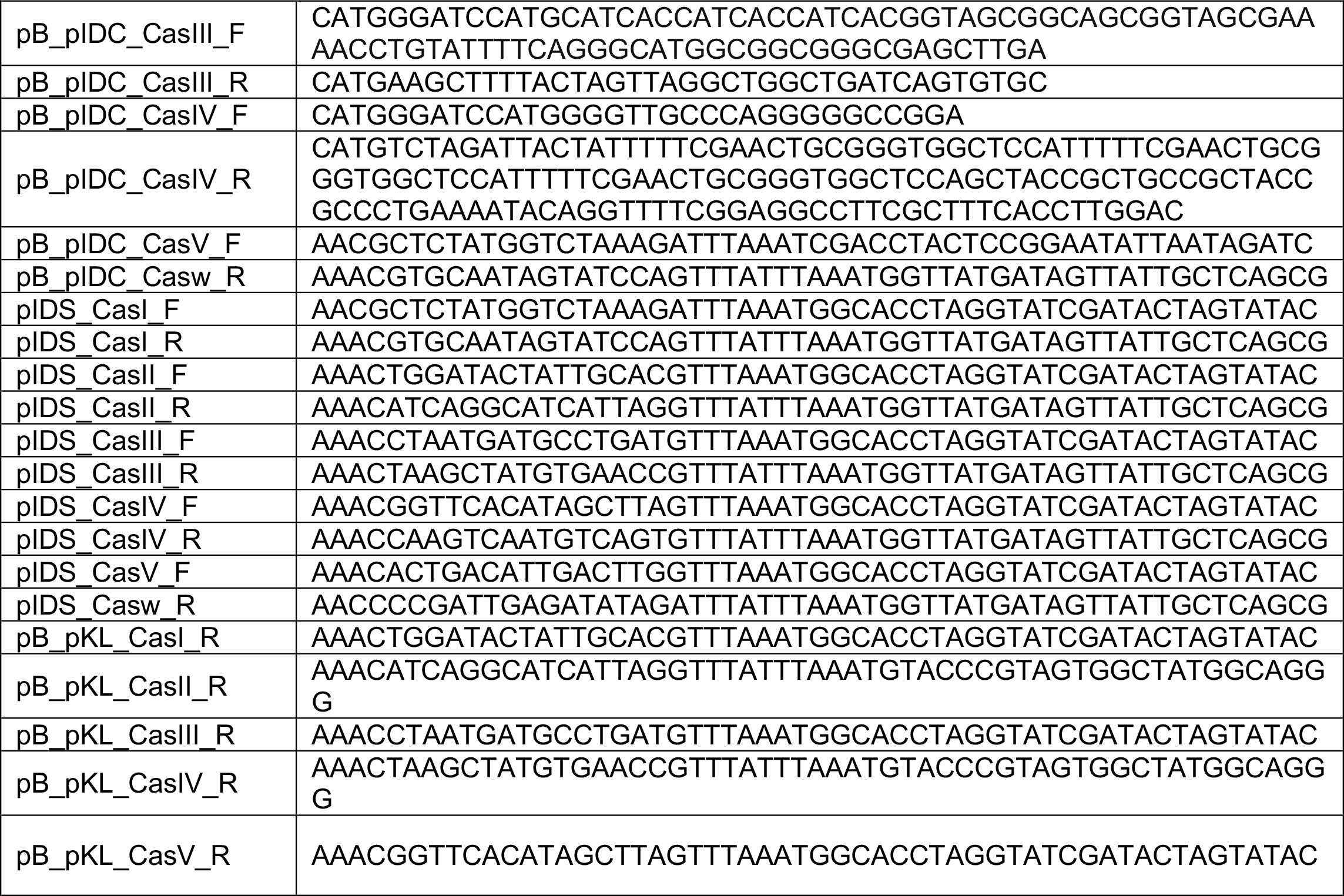
Table S5. Primer used in this study.

**Table S6.**
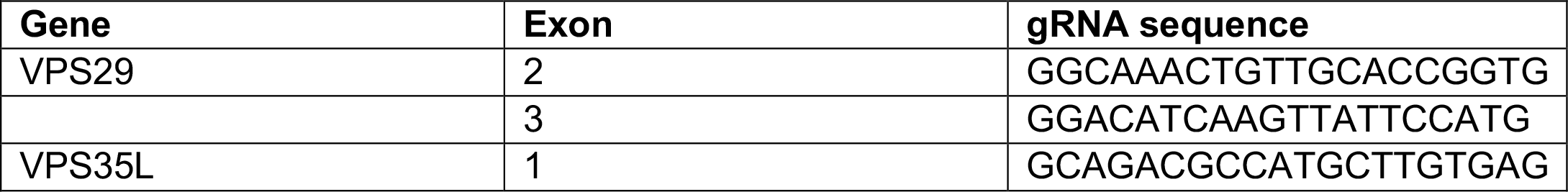
gRNA sequences for HeLa KO lines

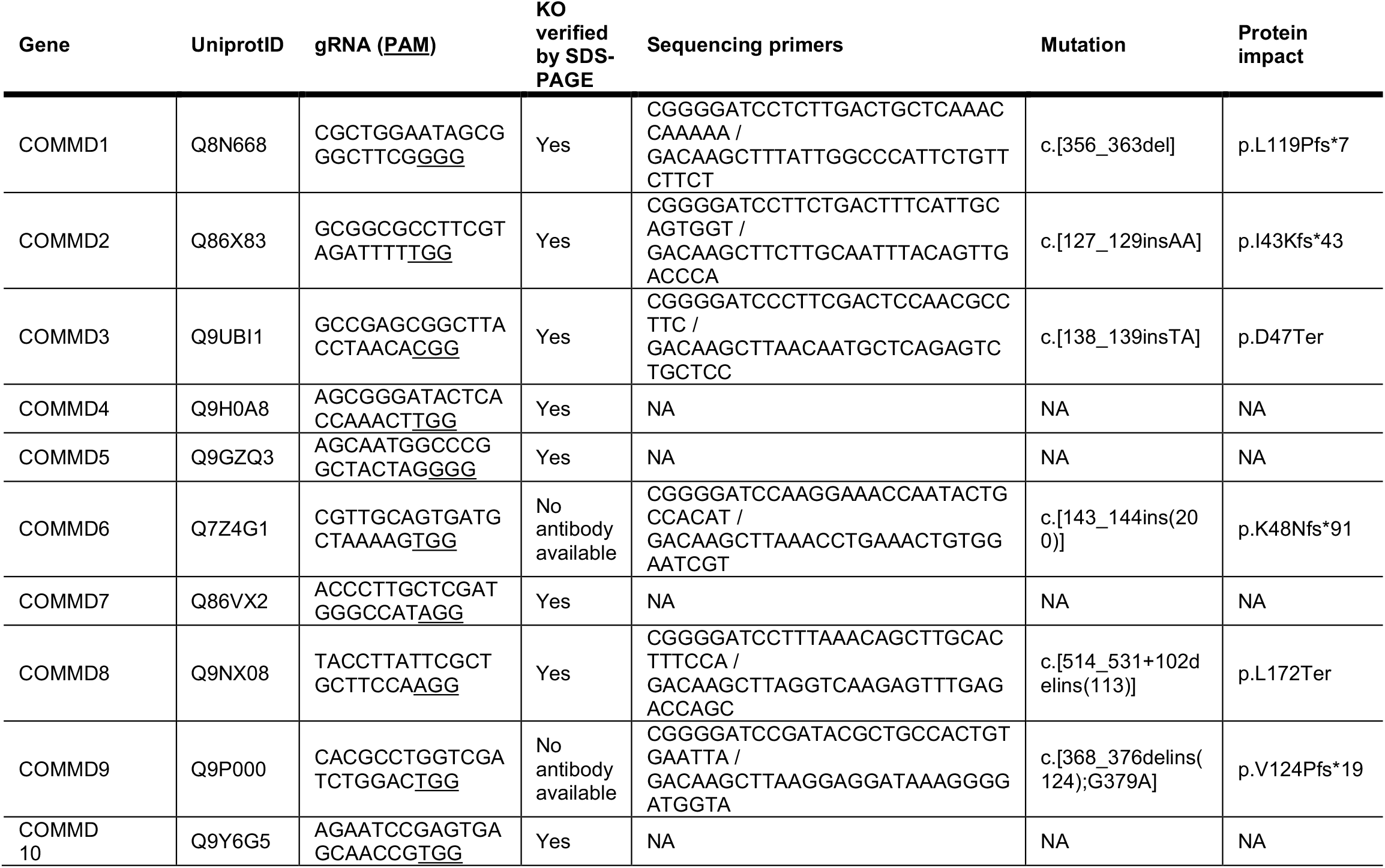
Table S7. Generation of eHAP KO COMMD lines.

**Table S8.** AlphaFold2 predicted complexes deposited in the ModelArchive^a^.

| **Model number** | **Species** | **Proteins** | **ModelArchive ID** |
| --- | --- | --- | --- |
| 1 | *Homo sapiens* | COMMD1-10 + CCDC22 (1-223) + CCDC93 (1-300) | ma-iplv4 |
| 2 | *Homo sapiens* | CCDC22 + CCDC93 | ma-9nv72 |
| 3 | *Homo sapiens* | VPS35L+ VPS26C + VPS29 | ma-3cag5 |
| 4 | *Homo sapiens* | VPS35L + VPS26C + VPS29 + CCDC22 (1-115) + CCDC22 (386-627) + CCDC93 (378-631) | ma-4097m |
| 5 | *Homo sapiens* | Combined models 1 + 2 + 3 | ma-ri7tb |
| 6 | *Danio rerio* | COMMD1-10 | ma-99f82 |
| 7 | *Salpingoeca rosetta* | COMMD1-10 | ma-xfc84 |

a. https://www.modelarchive.org

